# Integrating 12 Spatial and Single Cell Technologies to Characterise Tumour Neighbourhoods and Cellular Interactions in three Skin Cancer Types

**DOI:** 10.1101/2025.07.25.666708

**Authors:** P. Prakrithi, Laura F. Grice, Feng Zhang, Levi Hockey, Samuel X. Tan, Xiao Tan, Zherui Xiong, Onkar Mulay, Andrew Causer, Andrew Newman, Duy Pham, Guiyan Ni, Kelvin Tuong, Xinnan Jin, Eunju Kim, Minh Tran, Hani Vu, Nicholas M. Muller, Emily E. Killingbeck, Mark T. Gregory, Siok Min Teoh, Tuan Vo, Min Zhang, Maria Teresa Landi, Kevin M. Brown, Mark M. Iles, Zachary Reitz, Katharina Devitt, Liuliu Pan, Arutha Kulasinghe, Yung-Ching Kao, Michael Leon, Sarah R. Murphy, Hiromi Sato, Jazmina Gonzalez Cruz, Snehlata Kumari, Hung N. Luu, Sarah E. Warren, Chris McMillan, Joakim Henricson, Chris Anderson, David Muller, Arun Everest-Dass, Blake O’Brien, Huanwei Wang, Mathias Seviiri, Matthew H. Law, H. Peter Soyer, Ian Frazer, Youngmi Kim, Mitchell S. Stark, Kiarash Khosrotehrani, Quan Nguyen

**Author notes:** co-first authors (P. Prakrithi, Laura F. Grice, Feng Zhang).

## Abstract

Cutaneous squamous cell carcinoma (cSCC), basal cell carcinoma (BCC), and melanoma - the three major skin cancers - collectively comprise over 70% of all cancer cases. Despite their prevalence, much understanding of cellular interactions in the skin cancer microenvironment is needed, both in the outer skin layer where the cancer originates and at the deeper junctional and dermal layers into which it progresses. To address this gap, we integrated 12 complementary spatial and single-cell technologies to generate orthogonally-validated cell signatures, spatial maps, and interactomes for cSCC, BCC, and melanoma. Through comprehensive comparisons and integrating these spatial methods, we provided practical benchmarking guidelines for experimental design and analysis. By identifying keratinocyte cancer cells and melanomas, we found distinct signatures of these cells compared to non-cancer keratinocytes and melanocytes. Spatial integration of transcriptomics, proteomics and glycomics uncovered cancer niches enriched for cancer initiating cells (melanocytes or keratinocytes) and fibroblast and T-cell (MKFT) clusters, with altered tyrosine and pyrimidine metabolism. Ligand-receptor analysis across >700 cell-type combinations and >1.5 million interactions highlighted key roles for CD44, integrins, and collagens, with CD44-FGF2 emerging as a potential therapeutic target. Consistently, melanoma showed strong MFT interactions, validated by Opal Polaris, RNAScope, Proximal Ligation Assay and two additional single-cell spatial platforms (making a total of 14 technologies). For population-scale generalisation, genetic associations from >500,000 individuals were mapped onto spatial skin tissues, identifying SNPs enriched in domains containing melanocytes and T cells and their ligand-receptor pairs, shedding light on functional mechanisms linking genetic heritability to cells within cancer tissue. We built an interactive multiomics resource for exploring spatially-resolved molecular signatures and cellular crosstalk in skin cancer, available at https://skincanceratlas.com.

## Background

Although skin cancer is the most common neoplasm, its major subtypes, basal cell carcinoma (BCC), squamous cell carcinoma (SCC), and melanoma, differ markedly despite arising within the similar anatomical environment of the epidermal and dermal layers. Keratinocyte cancers BCC (∼75-80%) and cutaneous SCC (cSCC; ∼20%), are the most prevalent skin malignancies. Although associated with low mortality, their high incidence imposes substantial public health burdens, with an estimate of ∼3.5 million cases treated annually in the U.S., costing ∼$4.8 billion (Rogers et al., 2015; Guy et al., 2015). Still, about a third of skin cancer mortality is attributed to keratinocyte cancer (Czarnecki, 2024). By contrast, melanomas comprise <10% of diagnosed skin cancers but carry a markedly higher case fatality rate, constituting the majority of skin cancer-related deaths (Siegel et al., 2022). A deeper understanding of the tissue microenvironment and cellular interactions would explain the phenotypic differences among these skin cancer subtypes.

To understand early cancer progression and harness the therapeutic potential of immunotherapies for skin cancer, it is essential to understand the cellular composition of tumours and the cancer-immune interactions that underlies carcinogenesis. Approximately 30% of melanomas originate from benign naevi (Pampena et al., 2017), whereas most arise *de novo* from isolated melanocytes (Marks et al., 1990; Weatherhead et al., 2007). BCCs typically proliferate slowly and rarely metastasise, while cSCCs are more proliferative and a subset of cSCC (∼5%) is highly metastatic (Weinberg et al., 2007). Only ∼1% of pre-cancerous actinic keratoses progress to cutaneous cSCC, and determinants distinguishing progressor from non-progressor lesions remain unknown (Werner et al., 2013). Only 30-50% of advanced-stage melanoma patients respond to immunotherapies (George et al., 2017; Herrscher et al., 2020) and ∼50% of advanced-stage cSCC patients respond to immunotherapy treatment (García-Sancha et al., 2021).

Single-cell RNA sequencing (scRNA-seq) studies have investigated the cellular composition of BCC (Guerrero-Juarez et al., 2022; Yerly et al., 2022, Huang et al., 2023; Ganier et al. 2024), cSCC (Ji et al., 2020; Yan et al., 2021; Zou et al., 2023) and melanoma (Tirosh et al., 2016; Jerby-Arnon et al., 2018; Karras et al., 2022; Pozniak et al., 2024). Available melanoma single-cell datasets often contain relatively few cells (Tirosh et al., 2016; Zhang et al., 2022) and predominantly focus on acral melanoma or uveal melanoma, or on *in vitro* melanoma cell lines (reviewed by Lim and Rigos, 2024). A comprehensive comparison between the three cancer types at single-cell resolution and in the spatial context remains lacking.

This study applied state-of-the-art single-cell and spatial technologies to map skin cancer cells, cell-cell interactions, and microenvironments, comparing tumour micro-environment of keratinocyte cancers with melanomas. To enable cross-validation and comprehensive multi-modal views, we integrated 12 technologies, leveraging the complementary information in resolution, sensitivity, and throughput, adding spatial context to scRNA-seq data and spatially quantify both RNA, protein and glycan modalities. Spatial communities and single-cell gene signatures were systematically defined, and spatial cell-cell interactions (CCIs) were compared. The comprehensive analyses revealed shared and unique CCIs contributing to the divergent progression across the three skin cancer types. We presented a cross-validated spatial multiomics analytical framework to find 1) cancer cells; 2) gene signatures of cell types; 3) ligand-receptor based interactions; 4) community/neighbourhood composition; 6) interacting cell pairs and ligand-receptor pairs; 7) multimodal evidence for CCIs; and 8) genetic association signals with cell types in spatially defined regions. This framework is adaptable to different biological systems beyond skin cancer.

## Results

### Ultraplex, multimodal, multiplatform, single cell and spatial omics data resource for skin cancer

12 single cell and spatial imaging and sequencing technologies were applied to deeply investigate samples from 24 skin donors, consisting of patients diagnosed with cSCC (n = 7), BCC (n = 4), and melanoma (n = 10) and from non-cancer donors (n = 3) (**Table S1**). Controls include matched healthy skin samples from non-sun exposed skin of cSCC patients (n = 5) and samples from non-cancer donors (n = 3). External datasets are used for validations. Each biopsy was measured by up to 12 technologies as shown in **Fig 1**. Additional Curio Seeker and STomics data for melanoma samples were used for validation, so in total, data from 14 technology platforms were analysed. In the following sections, we present the results in two focal topics: a single cell and spatial atlas (**Fig1** to **Fig 4**) and an interaction atlas (**Fig 5** to **Fig 7**), followed by the integration with population genetics data of >500,000 individuals (**Fig 8**). Our database, **skincanceratlas (**https://skincanceratlas.com/), enables interactive exploration of the single-cell and spatial atlas and interactome **(Fig S1)**.

**Figure 1.**
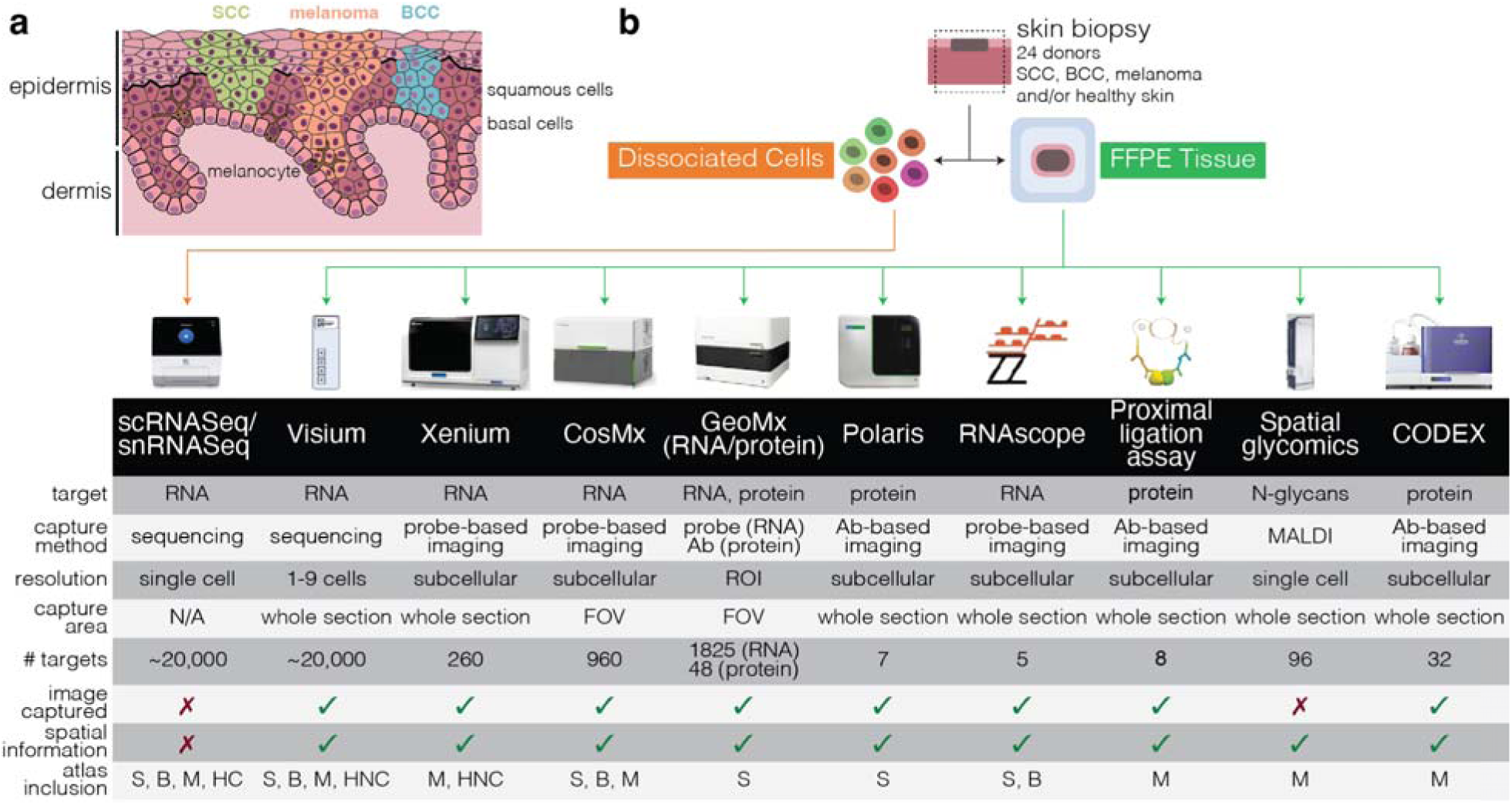
Integrating 12 single cell and spatial technologies to characterise cell type, spatial communities and cell-cell interactions in skin cancer. *(a)* Simplified cross-section of the human epidermis, highlighting the cellular origins of squamous cells, melanocytes and basal cells. Coloured regions represent the source cells of SCC (green), which originates from squamous cells, melanoma (orange), which originates from melanocytes, and BCC (blue), which originates from basal cells. Other cells and structures in the lower dermis layer are not depicted. *(b)* Overview of sample design and technologies used to generate data for this project. Technologies included are single cell RNA sequencing for fresh samples, single nuclei sequencing for formalin-fix d samples, Visium, Xenium, CosMX, GeoMX DSP for whole transcriptome, GeoMX DSP for proteins, Polaris, RNAscope, the proximal ligation assay, spatial glycomics and CODEX. Ab - antibody; FFPE - formalin-fixed paraffin-embedded; FOV - field of view; MALDI - matrix-assisted laser desorption/ionization; ROI - region of interest; S - cSCC; B - BCC; M - melanoma; HC - healthy (cancer patient); HNC - healthy (non-cancer patient donor).

### A comprehensive single-cell reference dataset of cSCC cancer with matched healthy and cancer samples

We first generated a high-quality single-cell cSCC reference dataset, then compared matched healthy and tumour samples to identify transcriptional shifts and cancer-specific gene signatures in cSCC cells. From 45,909 high-quality cells, we identified five major lineages (“Level 1” annotations), comprising endothelial cells, fibroblasts, melanocytes, keratinocytes, and immune cell types (**Fig 2a-c, Fig S2**). A second round of clustering defined eight major immune subsets (“Level 2” annotations; **Fig 2b-c**), and subclustering resolved 19 immune (sub)types (“Level 3” annotations; **Fig 2b, Fig S2a,c**), representing six T cell subsets, four macrophages, two NK clusters, two Langerhans clusters (LC), two DC clusters, a B cell cluster, a plasma cell cluster and a CD14+ monocyte cluster. Next, we characterised KCs, which comprised 70.6% of cells in the cSCC scRNA-seq dataset, and identified six subtypes: basal KCs (*KRT15*+, *KRT14*+), differentiating KCs (*PKP1+, KRT10*+), dysplastic KCs (KRT6B+, CD74+, S100A8+, S100A9+), KC interferon (*IFI27*+), KC cornified (*SBSN+, KRT2+, DSC1*+) and KC hair (combination of *KRT6B+, KRT17+, KRT16+, KRT5+*), (**Fig 2c, Fig S2b**). Mapping the cells defined by scRNA-seq data to the spatial CosMX skin tissues provides evidence for accurate cell type, displaying consistent annotation with anatomical regions of the skin (**Fig S2d**).

**Figure 2.**
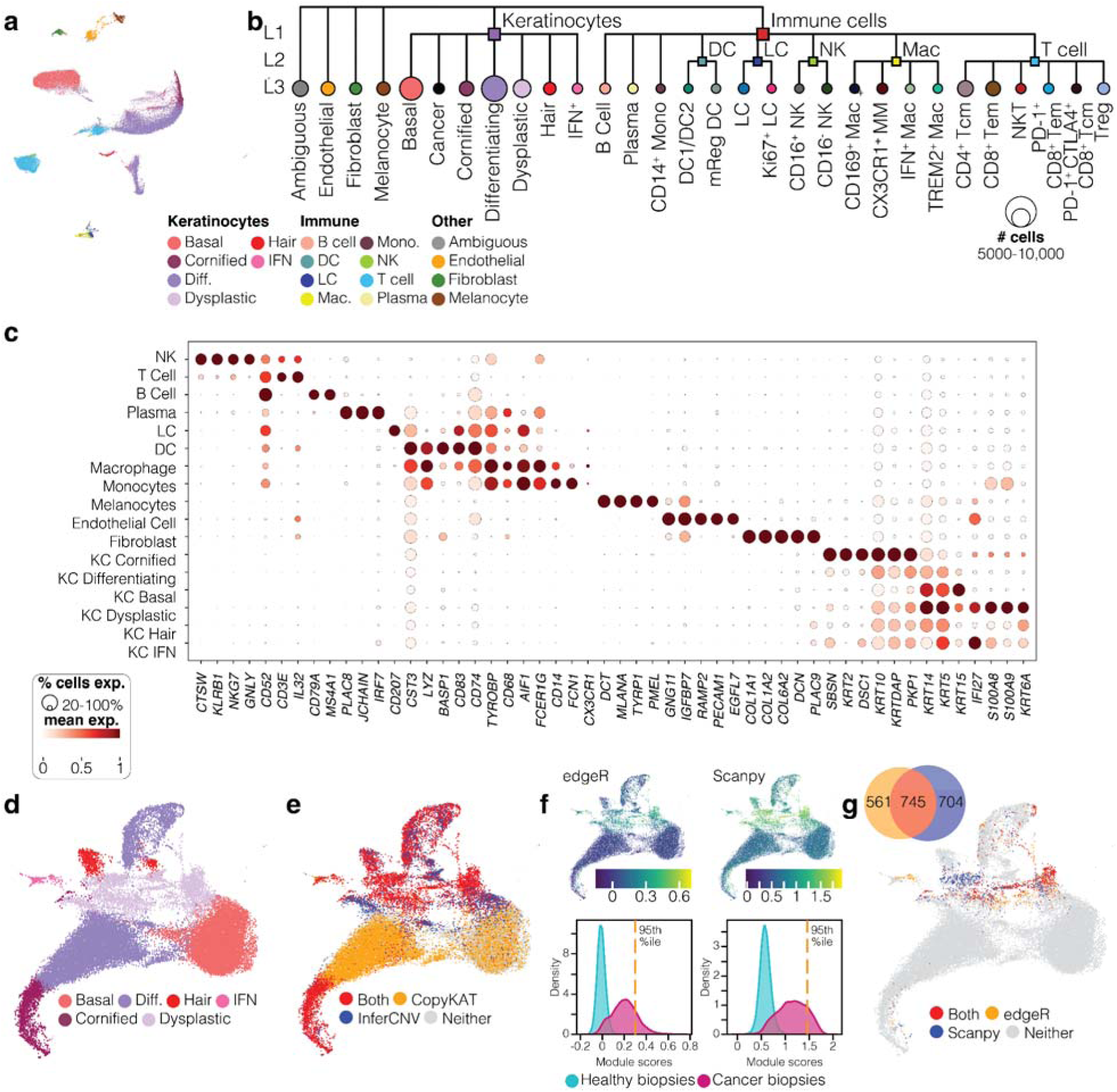
A single-cell reference dataset covering both cSCC, BCC cancer and healthy samples. *(a)* UMAP plot showing the integration of 45,909 healthy and cSCC cells from 11 samples of five patients, indicating results of Level 2 cell type annotation. Seventeen cell types were identified – eight immune cell clusters, six KC clusters, endothelial cells, fibroblasts, and melanocytes - plus an additional cluster of ambiguous cells. Diff - differentiating keratinocytes; IFN - IFN^+^ keratinocytes; DC - dendritic cell; LC - Langerhans cell; Mono. - monocyte; NK - natural killer cell; Mac - macrophage. *(b)* Dendrogram showing the three-level (L1 - L3) cell classification hierarchy, including L3 annotation of immune cells. MM - monocyte/macrophage; Tcm - central memory T cell; Tem - effector memory T cells. *(c)* Distinguishing markers of 17 Level 2 cell types. Markers are a combination of predicted markers for each cluster, plus known canonical markers for each cell type. *(d)* Subclustering of keratinocytes showing the six Level 2 subtypes. (e-g) Classification of cancerous KC cells. Candidate cells were first classified as being aneuploid (red) if both InferCNV and CopyKat predicted them to be as such (e). Cells were then assigned n “cSCC score” (Module score calculated based on the cumulative expression of genes differentially expressed in KCs in the cancer samples as compared to those from the normal samples) using differentially expressed genes identified using two different methods, edgeR and scanpy (f). Finally, cells were classified as KC Cancer (g) if they were classified as aneuploid (e) and also received an cSCC score above the 95th percentile of all cell scores (f). The venn diagram indicates the number of cells passing the module score threshold by edgeR and scanpy.

### High-confidence identification of cancer keratinocytes using gene signatures and inferred copy number variation

Single-cell efforts to define transcriptional signatures for KC cancer cells remain limited, particularly those that cleanly separate malignant KCs from non-malignant KCs and other lineages. We developed a stringent analysis pipeline to identify high-confidence cancer keratinocytes based on consensus aneuploidy from two CNV inference methods and elevated tumour-associated gene module scores (**Fig. 2d–g**). 745 high-confidence KC cancer cells were identified (with one cell from the normal sample, suggesting a 0.1% of possible false positive detection of KC cancer cells). Although we did not *a priori* restrict KC cancer cells to belong to a specific KC subtype, the majority (82.6%) of these cells were classified among dysplastic KCs, reinforcing malignant identity through three convergent lines of evidence. Additional evidence derives from a validation assessment using an external dataset, which contains a tumour-specific keratinocyte (TSK) population (Ji et al., 2020). We found a moderate but statistically significant overlap in marker genes (p = 1.7 × 10⁻²L) and reciprocal module scoring further demonstrated enrichment of TSK gene signatures within our KC cancer cluster and enrichment of our KC cancer gene set in the Ji et al. TSK cluster **(Fig S3)**. The high-confidence KC cancer cells were used for downstream analyses.

### A single-cell transcriptional reference resource for melanocytic lesions, spanning a disease spectrum from benign to dysplastic naevus and invasive melanoma

Limited single-cell datasets for cutaneous melanoma were available (Tirosh et al., 2016). Some recent work reported scRNA-seq data for a mixed acral and cutaneous melanoma study (Zhang et al., 2022) and for uveal melanoma and *in vitro* melanoma cell lines (as reviewed by Lim and Rigos, 2024). In this work, we produced a reference of melanoma cell types using single-nuclei sequencing of formalin fixed tissues, enabled by recent advances in highly sensitive probe-based gene counting protocols. Three patient samples were selected from a cohort of 50 patients, based on high-confident annotation by 23 pathologists. One sample was definitely invasive melanoma (5,591 cells), another as benign naevus (3,250 cells) and one severely dysplastic naevus sample (1,906 cells). From 10,747 single nuclei, we identified 11 immune cell types and five KC types, as well as endothelial, fibroblast, pericyte, Schwann cell, and sweat gland clusters (**Fig 3a-c, Fig S4**). Importantly, we combined multiple filtering layers to distinguish melanoma cells from melanocytes **(Fig 3d-g)**. Similar to the approach to define highly confident KC cancer cells in cSCC, we integrated CNV analysis, module scores, and spatial mapping of melanocytes to identify 118 “pure” melanoma cells (**Fig 3d-g**). These cells were used to find “clean” signatures of confidently defined melanomas compared to melanocytes and cSCC as described below.

**Figure 3.**
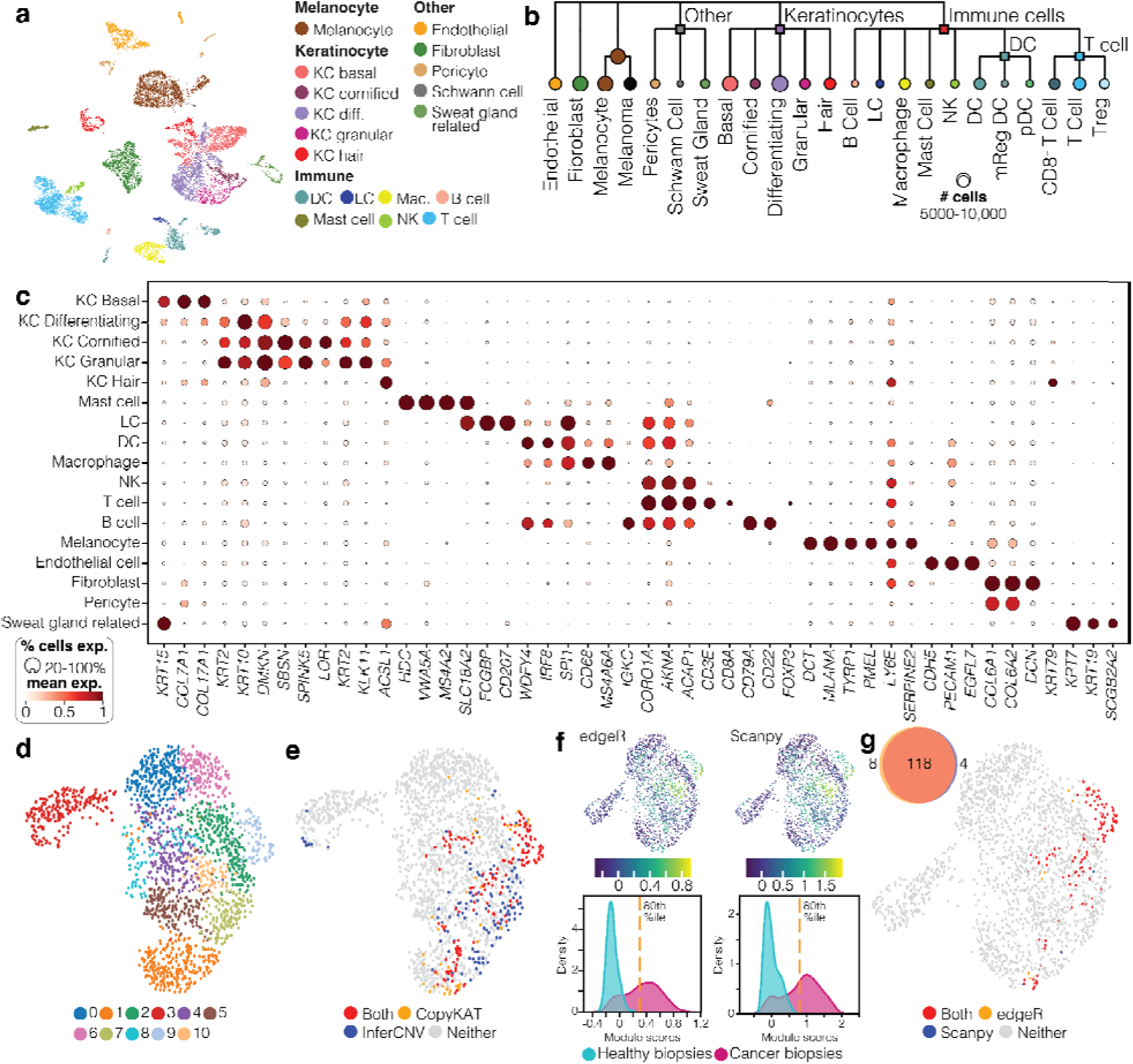
A single-cell reference dataset of melanoma cell types across the malignancy spectrum. *(a)* UMAP plot showing the integration of 10,747 melanoma cells from three patient samples, indicating results of Level 2 cell type annotation. Eighteen cell types were identified - melanocytes, seven immune cell clusters, five KC clusters, and five other cell types. *(b)* Dendrogram showing the Level 2 cell classification hierarchy. *(c)* Distinguishing markers of 18 Level 2 cell types. Markers are a combination of predicted markers for each cluster, plus known canonical markers for each cell type. *(d)* Result of Level 2 reclustering and cell type annotation for melanocytes. (e-g) Results for classification of cancerous melanoma cells. Melanocytes from the patient with the malignant tumour were classified as likely melanomas if they were both predicted to have aneuploid genomes (red) by both InferCNV and CopyKat (e). Cells were then assigned a “melanoma score” (f). Specifically, a module score was computed using genes upregulated in the melanoma sample compared to the benign sample using both edgeR pseudobulking and scanpy non-parametric test. For the sample from melanoma patient, a majority of the cells with a score >80th percentile cut-off were from the Melanoma sample cluster (Clusters 9 and 10), (g) and finally the cells inferred ‘Aneuploid’ by the CNV analysis and with a high module score by both the aforementioned methods are labelled as malignant melanocytes (red) as shown in the UMAP. The venn diagram indicates the number of cells passing the module score threshold by edgeR and scanpy.

**Figure 4.**
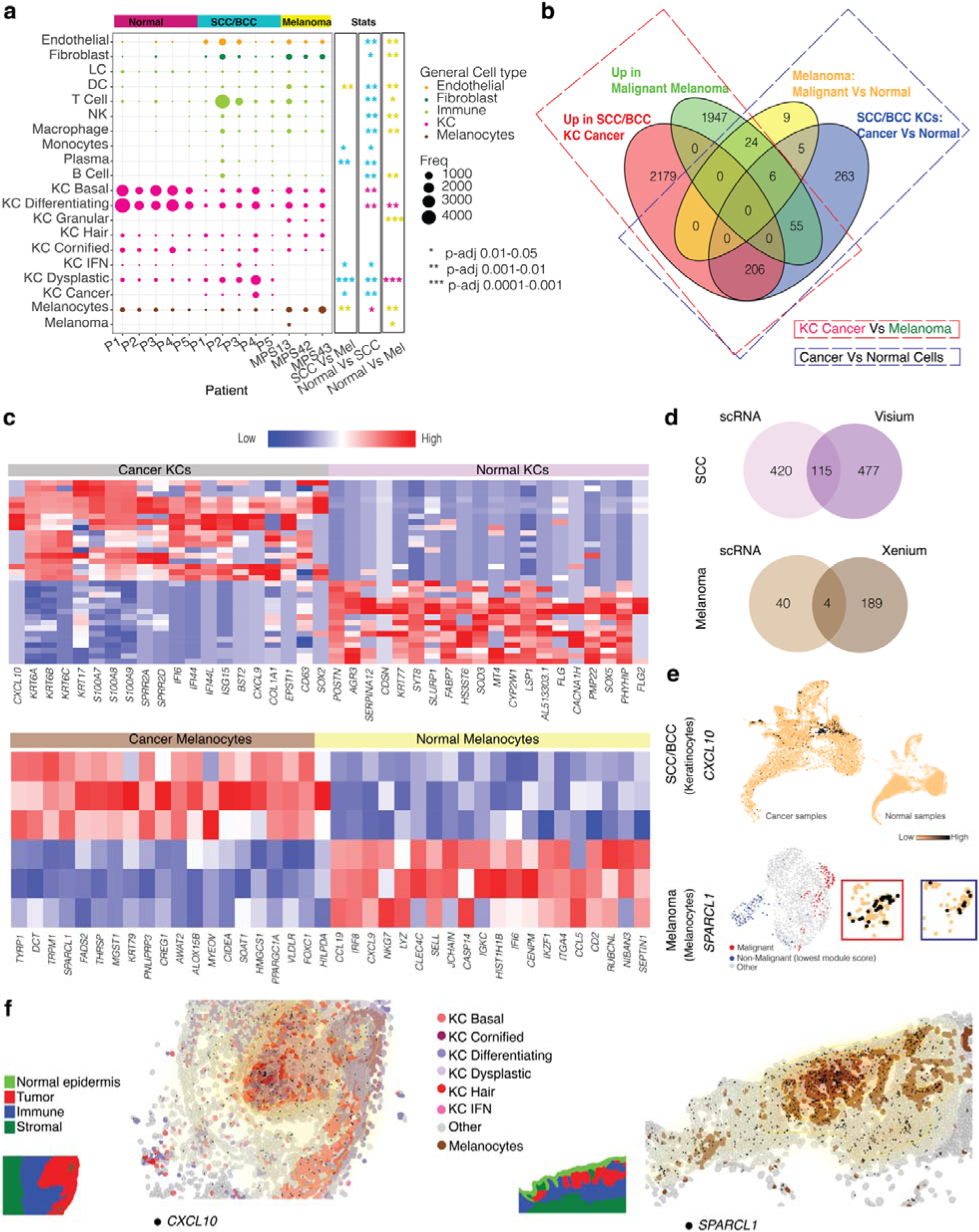
Transcriptional distinctions between cSCC, BCC and melanoma revealed by the comprehensive scRNA-seq datasets. *(a)* Dot plot showing the percentage of each Level 2 cell type within patient samples. Dots are coloured by cell type category and dot size indicates their percentage within each sample; all columns sum to 100. Results of differential abundance statistical tests are shown to the right, comparing abundance in cSCC vs melanoma, cSCC vs healthy skin, and melanoma vs healthy skin. Asterisks indicate the sample in which the cell type was found to be more abundant, either healthy skin (pink), cSCC-BCC (blue) or melanoma (yellow). *(b)* A venn diagram of the top significant upregulated genes across cancerous and non-cancerous KCs and melanocytes. (red) Upregulated in cSCC/BCC KC Cancer cells compared to Malignant Melanocytes from melanoma samples, (green) Upregulated in Malignant Melanocytes from melanoma samples compared to cSCC/BCC KC Cancer cells, (yellow) Upregulated in Malignant melanocytes compared to other melanocytes in melanoma samples, (blue) Upregulated in cancer KCs compared to other KCs in cSCC/BCC sample. *(c)* Heatmaps showing top 50 differentially expressed genes across Cancer vs Normal KCs (top left - The genes highlighted in red are consistently upregulated genes across platforms.), Melanocytes vs Melanoma (bottom - The genes highlighted in red are involved in cytokine interactions, and in blue for those in immunoregulatory pathways). Each row of the heatmap indicates a pseudo-bulked pool. *(d)* Integrative, multiple platform analysis of differentially expressed genes. From left to right, the Venn diagram shows the overlap between DE genes between cSCC cancer KCs vs normal KCs and Melanoma cells across scRNA-seq and for KCs in cancerous tissues compared to those from the normal tissues from non-cancer donors with spatial datasets of Visium, Xenium and CosMX. *e)* UMAP plot for scRNA-seq data showing the expression of CXCL10 in cancer vs non-cancer samples (top), which matches the location of KC cancer cells in UMAP shown in Fig 2. The UMAP below is that of the all the melanocytes as shown in Fig 3, with the malignant melanocytes highlighted in red and an equal number of melanocytes in the same sample that have the lowest module score for malignant melanoma signatures. The expression of SPARCL1 is shown in these two groups of cells marked by colour-matched boxes. *f)* Tissue gene expression plot of CosMX data showing two of the five shared markers SOX2 and LAMP3 (black dots). Pathological annotation of the region is shown on the left. The yellow lines and regions indicate a contour map of transcript density. A higher density is is observed in the tumour region.

**Figure 5.**
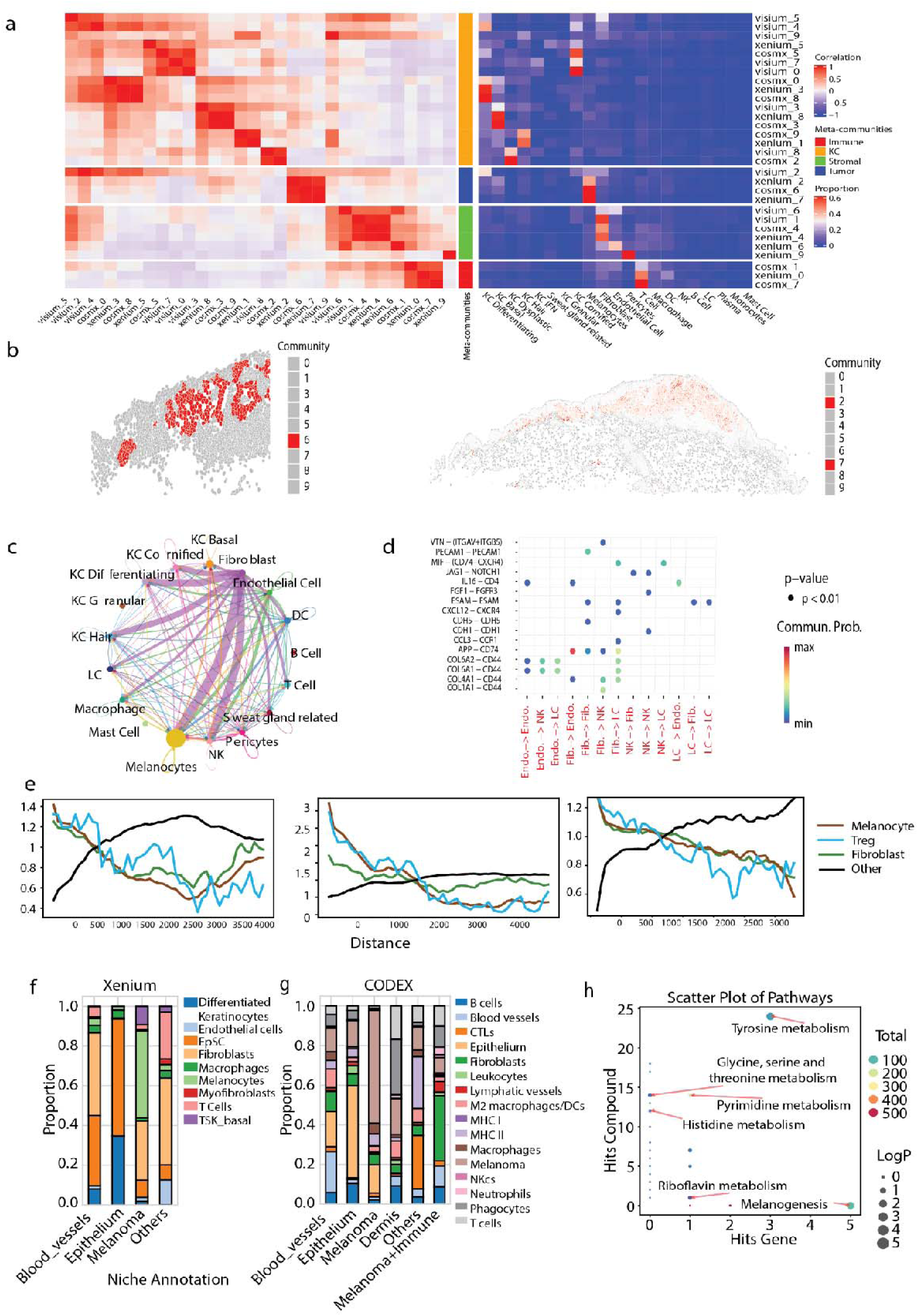
Integrative analysis across multiple patients and different technologies to reveal important pathways and interactions within robust spatial communities. *(a)* Cross-modality comparison of the ten communities identified for each of Visium, CosMx and Xenium. The 4 coloured bars represent super-communities (or meta-communities), which group the 10 finer communities based on their dominant cell type composition. Each row shows a community identified from one of the three spatial platforms. The left heatmap shows similarity across communities within and between technologies, measured by pairwise Pearson correlation values between communities based on their cell type composition. This allows similar communities across technology platforms and samples to be grouped to form meta-communities. The right heatmap shows the cellular makeup of each community (i.e. proportion of each cell type per community), providing information to label the groups of communities. The central annotation shows the broad classification of communities into immune, KC, stromal or tumour-related communities, based on the cellular makeup of each. *(b)* Spatial localisation of cells belonging to communities CosMx_6 (left) and Xenium_2 and Xenium_7 (right). Together with Visium_2, these communities form a meta-community that is enriched for melanocytes as shown in panel a. *(c)* Inter-community communication within melanoma CosMx_6. The chord plot visualises cell-cell communication mediated by Collagen signaling pathways, using the CellChat pathway database. Lines connect communicating cell types; line thickness represents greater communication between cell pairs. *(d)* Ligand-receptor interactions between pairs of cell types within the melanoma community CosMx_6. Top significant L-R pairs and corresponding cell type pairs are shown. *(e)* Cell type co-occurrence in CosMx samples between melanocytes and either other melanocytes (brown), Treg cells (blue), fibroblasts (green) or other cells (black). Each line plots the co-occurrence score (y-axis) between melanocytes and the test cell type calculated over increasing spatial distances (x-axis). The samples from left to right are melanoma 23346-105P, 30037-07BR and 6475-07FC. (f-g) Cell type proportions of communities identified in Xenium (f) and CODEX (g) for adjacent sections from the same sample (48974-2B). The melanoma community in both datasets is enriched with melanocytes and fibroblasts. (h) Joint pathway analysis using upregulated genes or proteins of the melanocyte communities in Xenium and CODEX data (shown in f and g), and highly expressed glycans of the melanocyte community in MALDI data (shown in Fig S13a). The proteins, genes, metabolites are mapped to KEGG metabolic pathways. The X-axis shows the number of genes/proteins from Xenium and CODEX data found in the pathway, while the Y-axis shows glycans in the same pathway.

### Overall shifts in fibroblast and T cell composition between non-cancer KC and cancer KC revealed by scRNA-seq analysis

Comparing matched healthy and cancer scRNA-seq samples from five patients, comprising 20,827 cells from cancer biopsies and 25,082 cells from healthy samples (**Fig S5a-b**), we found upregulation of tumour-associated genes such as *S100A7* and *KRT6B*, (**Fig S5c, d**) and increase in immune, fibroblast, and endothelial compartments in cancer (**Fig 4a**). The fibroblast proportion was consistently higher in cancer tissue, corroborating the role of fibroblasts in cSCC development (Schütz et al., 2023). The increase of both the lymphoid (T, B and NK cells) and myeloid (monocytes, macrophages and dendritic cells) in malignant skin matched the elevated expression of immune gene signature consistently found across patients (**Fig S5d, f-g**). T cells displayed the highest differences in abundance between cancer and non-cancer biopsies (**Fig 4a, Fig S5e**). We found core cSCC gene signatures, with 57 and 98 genes up- and down-regulated in cSCC, respectively (**Fig S5f**). These genes were enriched for immunological process GO terms, such as T-cell mediated cytotoxicity and antigen processing (**Fig S5g**). Basal and differentiating KCs were more prominent in healthy skin, whereas dysplastic and IFN KCs were enriched in cSCC samples, and no significant difference in the proportion of KC hair and KC cornified. The healthy skin sample signature was enriched for homeostatic processes, including “establishment of skin barrier” (GO:0061436) with genes like *CLDN1*, *IL18*, *KLF4*, *KRT1*, *NFKBIZ*, and *TP63*, suggesting potential loss of normal balance between proliferation and differentiation and skin integrity in cSCC (**Fig S5g**).

### Distinct gene signatures of cancer cSCC, suggesting roles of interferon gamma signalling, microenvironment remodelling and T cell recruitments

Using the scRNA-seq dataset, we identified 535 genes significantly upregulated in KC cancer cell compared to KC non-cancer cells in matched samples (**Fig 4b, c**). These genes highlighted chemokine signalling pathway and cytokine-cytokine-receptor interactions (*CXCL10, CXCL9, CCL20, CCL21, CCL8, IL32, CCL5, CXCR4*), interferon signalling (*IFI6, IFI27, IFI35, IFIT1, IFIT2, IFIT3, IFIT, IFITM1, ISG15, ISG20*), extracellular matrix remodelling (*MMP1, MMP2, MMP3, COL1A2*), de-differentiation (*SOX2),* and *IL-17/TNF* inflammatory signature *(S100A7, S100A8*, *S100A9, TNFSF10, TNFSF13B, TNFRSF21)*. To test the robustness of these signatures, we integrated results from scRNA-seq and Visium data, which add independent samples from non-skin cancer donors (**Fig 4c, d**). Consistent between scRNA-seq and Visium data, we identified 115 upregulated genes in KC cancer cells, enriched in epidermal remodelling and differentiation (*KRT6B, KRT16, GJB6, GJB2, and CTSC*) and interferon gamma signalling (IFI6, IFIT1, IFIT3, IFITM3, IGFB2, *ISG15, ISG20, CXCL10, CXCL11, CXCL9, IL15, CCL2, ISG20*). A visual example of one of the 115 shared genes, CXCL10, is shown uniquely expressed as in UMAP plots (**Fig 2g** vs. **Fig 4e**) and in spatial tissue plots (**Fig 4f**) consistent with histologically defined cancer regions show high specificity of this gene to KC cancer cells. *CXCL10 and CXCL9* were the most elevated in cSCC compared to normal skin (**Fig 4c**). It was shown that CXCL9 and CXCL10 from keratinocytes and fibroblasts can bind to CXCR3 expressed T cells and direct the recruitment of T cells to the skin (Kersh et al., 2024; Tuong et al., 2019).

### Single-cell -defined gene signatures of melanomas

Comparing transcriptional profiles between non-cancer and cancer cells of the same skin cancer type and comparing SCC cancer cells with melanoma, we found more variation between cancer cells from different lineages (2385 genes higher in cSCC than in melanomas, and 2032 upregulated in melanocytic lesions than in cSCC) than within a lineage, (44 upregulated genes in melanoma cells compared to melanocytes and 535 genes elevated in KC cancer compared to non-cancer keratinocytes) (**Fig 4b**). The 2032 genes higher in cSCC compared to melanomas were enriched for Myc targets, E2F targets, G2-M checkpoints, mTORC1 signalling and DNA repair pathways, whereas the 2385 genes higher in melanomas were most enriched in KRAS pathway, UV responses, EMT pathway, IL-2/STAT5 pathway, and Notch pathway (**Fig S6**).

By stringently defining melanoma cells as described above, we identified key melanoma marker genes associated with immune regulation and cancer invasion distinguishing melanomas from melanocytes (**Fig 4b-f, Fig S4, Fig S6**). The scRNA-seq data showed 44 genes highly expressed in melanomas compared to melanocytes, with a strong enrichment for melanogenesis and tyrosine metabolism (*TYRP1, OCA2* and *DCT*), (**Fig 4d, Fig S6**). Four shared genes were consistently found upregulated in both scRNA-seq and Xenium data, including *TYRP1*, *DCT*, *SPARCL1* and *TRPM1*. A visual example of a melanoma-specific gene, *SPARCL1*, is shown in **Fig 4e,f**. *SPARCL1* encodes an extracellular matrix protein, reported to be an upregulated marker in a melanoma population resistant to immunotherapy (Quek et al., 2024). *LGI3* and *SPIRE2* are in the 44 upregulated genes and are known to influence melanoma cell migration, invasion and melanosome transport. We noted that the scRNA-seq data lacked replicates and so the pseudobulk approach may not be sufficiently powered. Nevertheless, the three-way integration between scRNA-seq, Xenium and CosMX provided cross-validations (**Fig S7**). Moreover, the Xenium data derived from a ‘ground-truth’ baseline of skin samples from non-cancer donors in a different cohort strengthened the comparisons (**Fig S7)**.

### Spatial distribution of cells in stroma and cancer immune compartments shows enrichment of T cells adjacent to cancer cores in cSCC

To independently validate cell composition defined by the scRNAseq data, we applied GeoMX Cancer Transcriptome Atlas (CTA, 1820 oncogenes) and GeoMX immune-oncology protein panel (48 proteins) for cancer (panCK+) and stromal-immune regions (CD45+), (**Fig S8; Supplementary Note 1**). Samples from three cSCC patients with matched scRNA-seq data were used (R01, P04 and B18; **Fig S5**). With the 48-protein GeoMx panel, we confirmed the presence of cell subtypes in the Level 2 and Level 3 annotation, including M2 macrophages (CD163, CD68), B cells (CD20), CD8+ T cells (CD8), CD4+ T cells (CD4), DCs (CD11c), fibroblasts (FAP-alpha, Fibronectin), and Treg cells (FOXP3, CD25), (**Fig S9, Fig S10**). The cell type deconvolution results for GeoMX CTA data show the heterogeneity between patients, while highlighting the high proportion of T cells across immune-stromal regions adjacent to cancer, with the highest abundance of the CD4+ T cell, followed by Treg and relatively fewer CD8+ T cells (**Fig S9f**).

### A spatial reference dataset of 21 cell types cross-validated using Visium, GeoMX, CosMX, Xenium, CODEX and spatial glycomics data

The 21 cell-type signatures for cSCC, BCC and melanoma from scRNA-seq/snRNA-seq data were mapped to the spatial tissues in the Visium data, including 5x cSCC, 3x BCC and 4x melanoma samples across 9x patients (**Fig S11a**, **Table S1**). High consistency between histological features, cell types and gene markers was observed (**Fig S11a,b**), suggesting an accurate data resource for downstream analyses. The spatial transcriptomics reference dataset covers three resolutions, including at the regions of interest level (GeoMX WTA and GeoMX protein, often >100 cells), spot resolution (Visium, ∼10 cells), and single-cell/subcellular resolution (CosMX, Xenium, CODEX). For example, the projection of scRNA-seq cell-type signatures to spatial CosMX data demonstrated an accurate distribution of cell types such as keratinocytes, melanocytes, fibroblast and immune cells to distinct layers in the skin, corroborating the accuracy of our scRNA-seq cell-type annotation (**Fig S2**, **Fig S11b**). With the subcellular resolution in CosMx data, we observed correspondence between the single-cell and single molecular localisation of RNA markers for KC cells, for examples *S100A8* and *KRT17* (**Fig S11**). The tumour region was consistently and independently mapped by Xenium (**Fig S11c**), CODEX (**Fig S11e, d**) and glycomics data (**Fig S11f).** More detailed visualisation and list of markers are shown in **Fig S14-S16.**

### Single-cell spatial heterogeneity analysis suggested a more complex tumour community in melanomas than in cSCC and BCC

With the spatial mapping of CosMX single cell data, we characterized cellular diversity of the cancer microenvironment across the three major skin cancer types. Rao’s quadratic entropy score suggested a high level of heterogeneity correlated with the mixed distribution of diverse immune cell types, for example an FOV with B cells, plasmacytoid dendritic cells (pDCs), myeloid and T cells (**Fig S12a, b).** Indeed, we observed the highest heterogeneity scores in immune-rich FOVs (**Fig S12d**). To compare cell type heterogeneity across the different skin cancer subtypes, we grouped scores across all FOVs by cancer type, including 30 FOVs from four melanoma patients, 24 FOVs from two BCC patients, and 27 FOVs from three cSCC patients (**Fig S12c, d**). We detected a significant increase in cell type heterogeneity score in the melanoma samples compared to in cSCC cancer (**Fig S12c, d**). However, we noted that heterogeneity assessment would require further validation in independent cohorts.

### Reproducible cancer communities across multiple samples and spatial transcriptomics platforms highlight the importance of fibroblast and T cells proximal to cancer cells

A key challenge in identifying functional tissue neighbourhoods (communities) is that they represent novel patterns derived from spatial analysis, making it unclear whether they are reproducible across biological replicates and technological platforms. To address this, we integrated three spatial transcriptomics datasets (Visium, CosMX, and Xenium), consolidating neighbourhood information for each cell/spot into a shared matrix to identify meta-communities, consisting of communities with similar cell type composition (**Fig 5a**, see Methods). In particular, we identified a meta-community comprising Visium_2, Xenium_2, Xenium_7, and CosMX_6, all enriched for melanocytes surrounded by fibroblasts, basal KC, and T cells across all samples, herein refer as MKFT community (**Fig 5a**). The MKFT meta-community was shared between samples and technologies. Comparing across cancer types it appeared to be significantly more abundant in melanoma samples than in BCC and cSCC (**Fig S17a, b**). We then characterised the meta-community by interactions between the cells within the communities through ligand-receptor coexpression (**Fig 5c, d**) or cell-cell colocalization (**Fig 5e**). Overall, fibroblast interactions were particularly dominant, especially Fibroblast with melanocytes (**Fig 5c**). Within the melanoma-associated CosMX_6 community, interactions were highly enriched for collagen signalling, especially between collagens and CD44 (including: *COL6A1, COL6A2, COL4A1, and COL1A1*) (**Fig 5d**). CD44 marks antigen-experienced T cells and promotes their adhesion to and infiltration of the tumor microenvironment. The co-localization analysis in the melanoma communities in three Xenium samples shows that Treg and Fibroblasts have a high co-occurrence probability with melanocytes **(Fig 5e).**

### Cross-modality community analysis characterised metabolic signatures in the melanoma microenvironment

We investigated the molecular signatures of the melanoma community across multiple analytes (i.e. RNA, protein and metabolites) in spatial context to gain a holistic insight into cell and tissue-level functions that may not be fully captured with a single modality (**Fig 5f-h**). The Xenium (RNA), CODEX (protein) and mass cytometry imaging (MALDI MSI for spatial glycomics) data independently defined melanoma community, with distinct molecular markers, as shown in **Fig S13**. The single cell Xenium data defined 17 cell types where their locations matched the pathological annotation of the tumour and immune cells (**Fig S11c, Fig S14a**), including all key KC cell types. In contrast, the CODEX data could not map KC cells, due to lack of protein markers for these cell types, but could clearly pinpoint additional immune cell types such as the M2 Macrophages and Neutrophils (**Fig 5g, Fig S11d**). CODEX data stratified melanoma communities into two, including one more infiltrated with immune cells (**Fig 5g**, **Fig S15c**). Notably, both CODEX and Xenium data defined melanoma communities with high abundance of fibroblast (**Fig 5f, g**). Although the cell-type clustering using glycomics is less defined, it is clear that melanoma/melanocytes exhibited unique metabolomic signatures of 28 consistent metabolic markers compared to other cell types in the remaining clusters (**Fig S11f, Fig S13a, Fig S16**).

Importantly, joint pathway analysis of genes/proteins with metabolites provided three lines of evidence that Tyrosine metabolism was enriched (supported by 3 genes/proteins and 24 metabolites), (**Fig 5h**). Tyrosine is a critical precursor for melanin production in melanocytes and dysregulation in this pathway can contribute to melanoma development or progression (Najem et al., 2021). This analysis also showed the upregulation of Pyrimidine metabolism in the melanoma community, a pathway fundamental for DNA and RNA synthesis and is linked to sunlight-associated melanomas, melanoma progression and treatment resistance (Edwards et al., 2016; Santoriello et al., 2020), (**Fig 5h**).

### Non-cancer KC and cSCC displayed differential interactions between immune cells, fibroblast and keratinocytes

The matched scRNA-seq cSCC dataset showed overall more L-R interactions in cancer samples than the matched non-cancer samples, with core sets of L-R pairs shared across patients, enriched for immune-related functions including MHC class II complex assembly (**Fig S18d**). While KCs were the dominant ligand contributors in healthy samples, we observed a consistnet increase in the total number of predicted L-R interactions and increased signalling from fibroblast and immune cells (**Fig S19**).

### A robust pipeline to spatially define and compare CCI

To improve accuracy of scRNA-seq predictions of L-R interactions, we performed spatially-constrained two-level permutation (SCTP; **Supplementary Note 2),** (Pham et al., 2023). This analysis revealed interactions that were predicted by scRNA-seq for L-R pairs that were not colocalised in the spatial tissue, suggesting possible false detection (e.g., XCL1-XCR1). SCTP also found L-R interactions that were missed from scRNA-seq data analyses, but were detected by Visium (e.g. *WNT5A-ROR1*), (**Fig S20a**). Indeed, we found cases where the scRNA-seq missed the interactions, while all three platforms, Visium, CosMX and Xenium strongly supported the interactions both by statistical significance test and by visual co-expression of the L-R pairs between neighbour pairs (e.g., *CXCL12-CXCR4*, *CCL9-CCR7*) (**Fig S20b**). To identify and compare highly confident interactions, we then applied a multi-platform, multi-sample CCI (MMCCI) algorithm to integrate the data to find interactions consistent across biological replicates (Hockey et al., 2024). MMCI was then used for differential interaction analysis at cell-type network levels (comparing edges connecting two interacting cell types) or at L-R levels (comparing L-R interaction scores).

### Differential ligand-receptor interactions specific for BCC, cSCC or melanoma show important roles of angiogenesis, integrins, and fibroblast growth factors

Applying MMCCI for the integrated CosMX and Visium datasets, we consistently found 16 L-R pairs highly expressed in BCC, 17 in cSCC, and 37 in melanoma (**Fig 6a**). For BCC, among the L-R pairs enriched, we found strong interactions involving interleukins (*IL6-IL6R, IL6-IL6ST, IL1B-IL1RN-IL1R2*), chemokines (*CXCL2-CXCR1, CCL2-ACKR4*) and fibronectins (*FN1-ITGB8* and *FN1-ITGB6*), all having roles in angiogenesis (Villani et a., 2021). Interactions in the canonical Wnt signalling pathways (*WNT5A-FZD7* and *WNT5A-FZD8*; also supported by *EPCAM-EPCAM* interactions) were found more active in BCC than in cSCC and melanoma. The Wnt pathway is crucial in coordinating with the Hedgehog signalling in BCC to maintain the proliferative state of BCC cells and sustain cancer stem-like cells in BCC (Yang et al., 2008).

**Figure 6.**
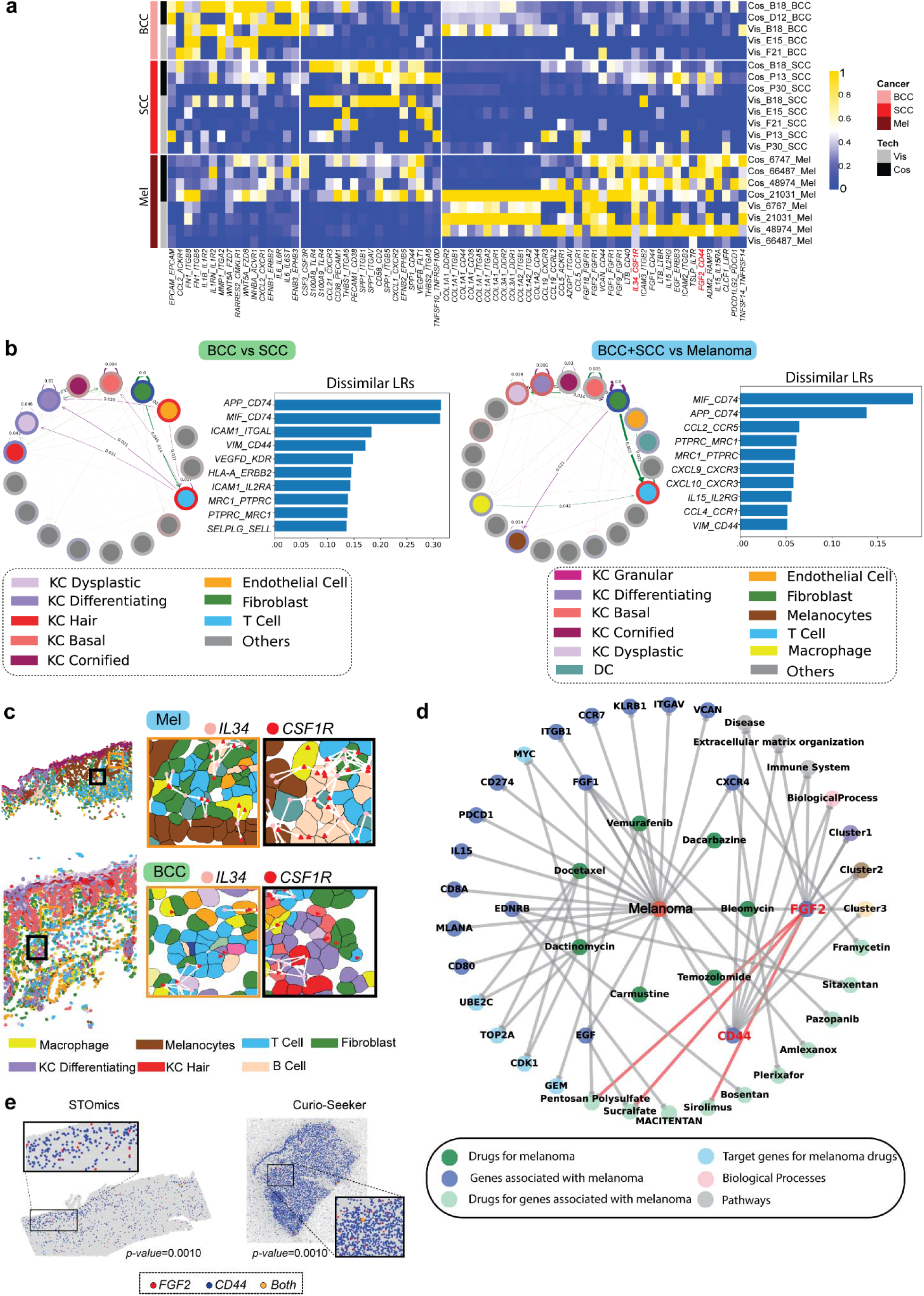
Differential cell-cell interaction using integrative analysis of Visium, CosMx, Xenium, Stereo-seq and Curio data. *(a)* Heatmap of L-R scores for L-R pairs enriched per cancer type, with a consistent trend across samples and the two Visium and CosMx platforms. Differentially expressed L-R pairs were calculated comparing each cancer type vs the others using a pseudobulked L-R scores with 3 pools per sample. Each heatmap row is a distinct CosMx or Visium sample. The two L-R pairs specific for melanoma IL34-CSF1R and FGF2-CD44 were used for experimental validations. *(b)* Comparing differential interactions at cell type level for BCC vs cSCC and for the integrated BCC-cSCC vs Melanoma using all L-R pairs. For cSCC vs cSCC BCC comparison, purple arrows show more interactions in BCC and green arrows indicating more in cSCC. For BCC-cSCC vs Melanoma, the purple arrows show more in melanoma and green arrows show more in cSCC-BCC. *(c)* Spatial mapping of cancer type-enriched L-R pairs in CosMx data. One of the L-R pairs that was significantly different between cancer types across technologies in Panel a, namely IL34-CSF1R (higher in melanoma) is shown. It is visualised in FOVs from melanoma sample (top) and BCC sample (bottom). For each cancer type, the cell type annotation of the FOV is shown (top left) with orange and black boxes indicating the highlighted regions (top right). Magnified boxes (top right) show the presence of the ligand (pink) and receptor (red), with white arrows showing the connections between ligands and receptors of nearby cells. *(d)* Melanoma drug target graph integrating multiple biological and pharmacological knowledge types. Nodes represent genes, drugs, and biological functions. Level 1 connections show melanoma-associated genes and drugs targeting melanoma. Level 2 links display drugs targeting the melanoma-associated genes from Level 1 and a broader gene set targeted by drugs in the network. All genes in the graph are either upregulated or have high ligand-receptor scores. Clusters 1, 2, and 3 are pathways enriched with genes shown in the graph. *(e)* A significant co-localization of cells expressing FGF2 and CD44 is replicated for melanoma samples across different high-resolution technologies.

**Figure 7.**
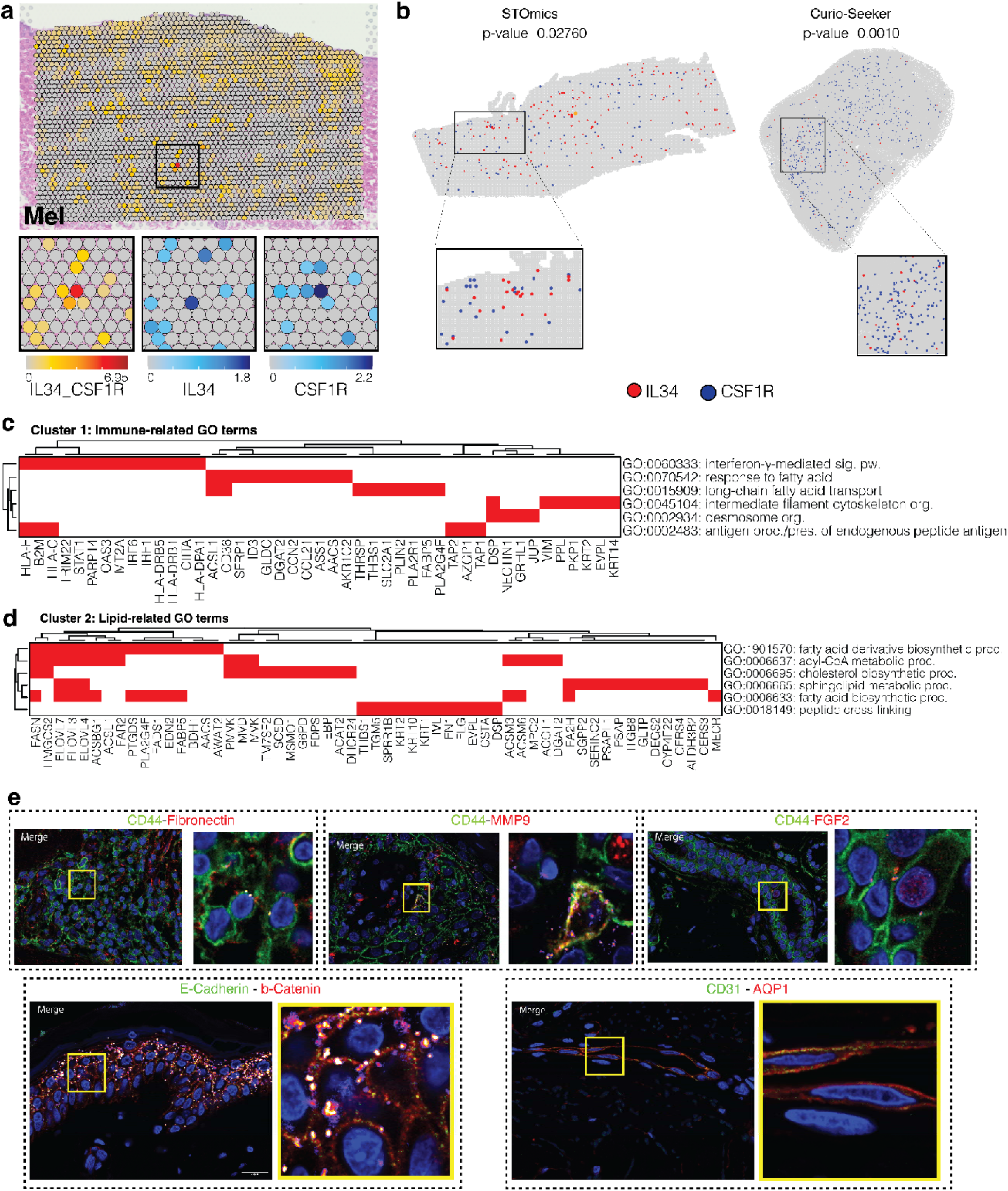
Multi-omics validations of L-R interaction *(a)* Exemplar spatial plots showing the L-R score for IL34-CSF1R from patient 48974. The black box indicates a region highlighted below the main image. Here, zoomed-in boxes show the IL34-CSF1R L-R score (left) and IL34 (middle) and CSF1R (right) gene expression for the same tissue region. *(b)* High-resolution spatial transcriptomics for melanoma samples using STOmics and Curio-Seeker, showing cells expressing IL34 and CSF1R and a significant co-localization of these cells. (c-d) Heatmaps indicating grouped GO terms and associated genes that are enriched in IL34-CSF1R positive spots in melanoma samples compared to IL34-CSF1R negative spots. GO term groups were calculated by k-means clustering (k = 3) of GO semantic similarity scores; two such groups are shown here. The full heatmap is shown in **Fig S10b**. *(f)* Proximal ligation assay (PLA) for validating CD44 interactions in melanoma (top). A merged image of signal for the ligand and the receptor and a zoom-in window highlighting the interaction on the cell membrane. A positive PLA signal is visible if two interacting proteins are in a proximity less than 20 nm. The bottom panels show signals for positive (E-Cadherin-b-Catenin) and negative (CD31-AQP1) controls.

**Figure 8.**
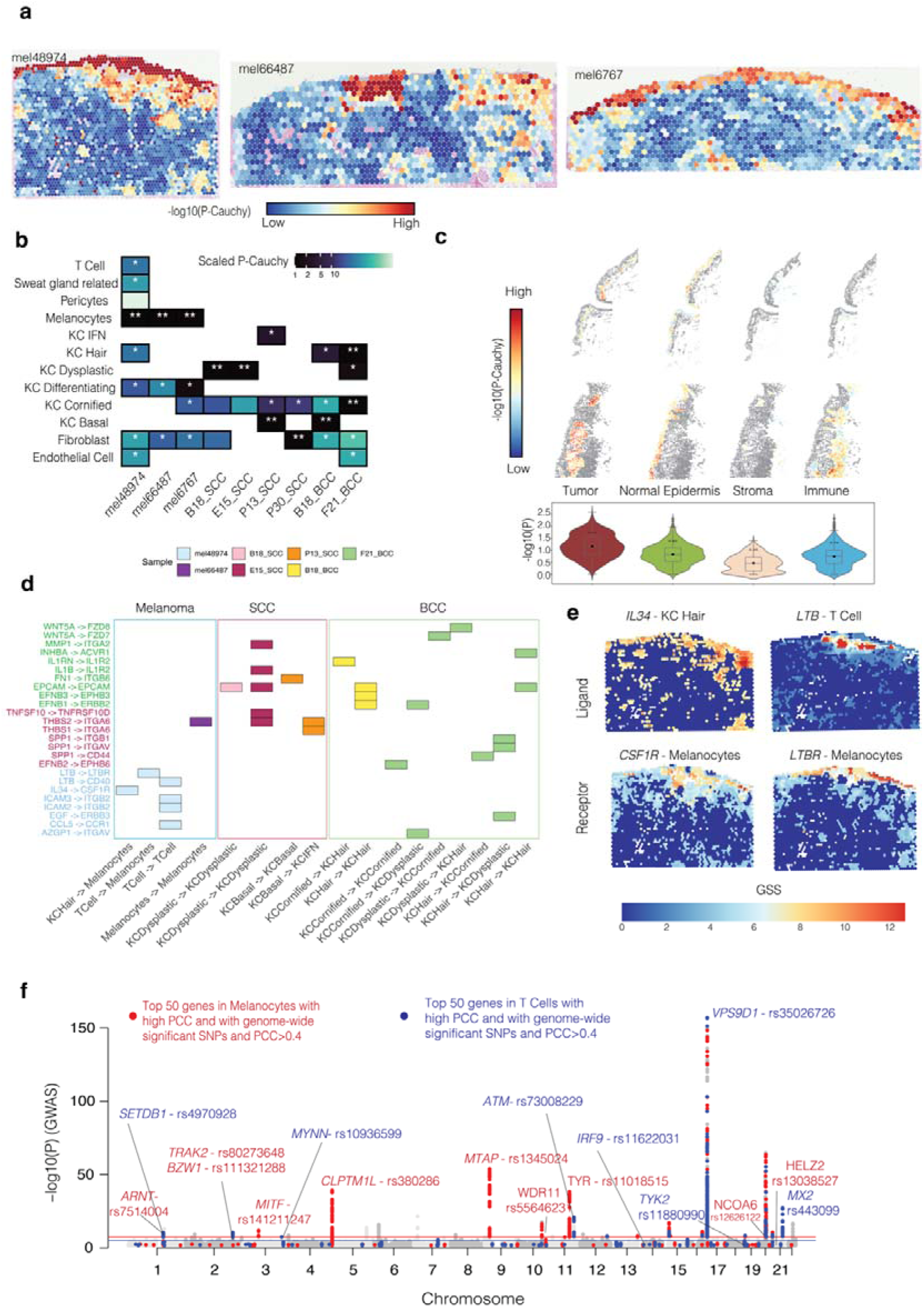
Mapping genetics effects from genome-wide association studies for melanoma, cSCC, and BCC to spatial domains and cell types. *(a)* Gene specificity score (GSS) and association of spatial spots with skin cancer heritability. GSS score for each gene in a spot/cell represents the enrichment of the gene as a top rank most abundant gene in the spot/cell and its neighbour spots/cells in an anatomical region, a spatial domain, or a cell type. The p-value shows the spatial heritability enrichment significance of a spot with a trait based on SNPs mapped to the genes with high GSS scores (one-sided Z-test for stratified coefficient different to 0). The p-value is more significant if the SNPs that are mapped to the high GSS genes explain a higher proportion of heritability for the trait. *(b)* Cell types with the highest enrichment of heritability explained by SNPs tagged to GSS genes of cells in a cell type. The white asterisks indicate the significantly enriched cell-type for heritability of cutaneous melanoma, cSCC and BCC traits. Non-significant data is shown as empty. The P-Cauchy values are scaled such that the most significant (**) values are ‘1’ and the other significant cell types are indicated by *. The scaling was done as Scaled_P_Cauchy = P_Cauchy / min(P_Cauchy). *(c)* gsMAP significance spatial heritability enrichment is shown at single-cell resolution across the tissue (upper tissue plots) or per annotated skin regions (lower violin plots) from the cosMx data of the sample mel48974, suggesting significance in tumour regions. *(d)* L-R pairs with significant association with SNP heritability explained by the corresponding cell types. The rectangles show cases where both L and R genes had PCC >0.3 between GSS of the gene and the gsMAP P-values (the significance level for the LD stratified coefficients for the spot bigger than 0). The results suggest which L-R pairs are related with the heritability of a cell type pairs. *(e)* GSS of two L-R pairs showing specificity of the L and R genes to tissue regions at the immune-rich dermal layers and the epidermis of the skin. *(f)* Manhattan plot showing top significant GWAS SNPs co-localizing with genes in melanocytes (red) and T cells (blue) that had the highest Pearson correlation between GSS and the gsMAP trait association P-value or associated with SNPs with genome-wide significance. The Y-axis shows the - log(P-value) from GWAS analysis.

For cSCC, strong interaction of Osteopontin (*SPP1*) was found, particularly with the pairs *SPP1-ITGB1* and *SPP1-ITGAV* (uniquely expressed in cSCC), while *SPP1-CD44* and *SPP1-ITGB5* were stronger compared to in BCC and Melanoma. Through interactions with integrins and *CD44*, *SPP1* may activate intracellular signalling pathways like *PI3K/Akt, MAPK/ERK*, and *FAK* to promote cell survival, proliferation, and growth (Anborgh et al., 2010). The MAPK/ERK pathway and PI3K/Akt pathways can be activated by *FLT1* and *CSF3-CSF3R* interactions. The interactions associated with angiogenesis like *CD38-PECAM1, VEGFB-FLT1* (*VEGFR-1*), and *CXCL1-CXCR2* were also higher in cSCC, suggesting that angiogenesis was upregulated in both BCC and cSCC, but through different regulation pathways. In addition, SCC samples also showed more interactions of calprotectin (*S100A8* and *S100A9*) with *TLR4* on immune cells, contributing to inflammation in the tumour microenvironment.

For melanoma, we found marked enrichment of collagen interactions (e.g., *COL1A1*, *COL1A2*, *COL3A1*; making 15 out of all the 37 L-R pairs that were specific for melanoma compared to BCC and cSCC). Particularly, type I collagen (*COL1A*, *COL1B*) and integrin receptor (*ITGA*, *ITGB*) families displayed strong interactions in melanoma. These included *COL1A1-ITGA2*, *COL1A1-ITGB1*, *COL1A1-ITGA5*, *COL1A2-ITGA2*, and *COL1A2-ITGB1* (**Fig 6a**). Changes in integrin activity have been implicated in differential metastatic and invasive risks in melanoma (Xu et al., 2017). The collagen-integrin interactions play key roles in microenvironment remodelling, creating pro-tumorigenic and immunosuppressive niches, promoting angiogenesis, and activating the MEK/ERK signalling pathways (Hayashido et al., 2014). Among the top 12 melanoma-specific collagen interactions, four L-R pairs were involved in DNA damage responses through interactions with *DDR1* and *DDR2*. The second most common ligand-receptor pairs that were uniquely increased in melanoma involve fibroblast growth factor, with six out of 37 interactions, including *FGF1*-*CD44*, *FGF2-CD44, FGF2-FGFR1, FGF18-FGFR1, FGF1-FGFR1, FGF9-FGFR1*. FGF signalling can enhance melanoma cell proliferation by activating downstream signalling pathways such as MAPK/ERK, PI3K/AKT, and JAK/STAT, which play key roles in cell cycle progression and survival, and some are therapeutic targets for melanoma (e.g., FGF2/FGFR signalling).

### Different interactions between cell-cell pairs highlight the roles of fibroblast, T cells and melanocytes/keratinocytes

For interactions at cell type level, MMCCI multiplatform and multimodal analysis showed that cSCC and BCC were more similar to each other than to melanoma (**Fig S21a**). The most common interactions across the three cancers involve fibroblasts with immune cells and KC cells (**Fig S21a**). We observed stronger fibroblast to T cells interaction in BCC compared to in cSCC (**Fig 6b** - left) and similarly in cSCC-BCC compared to in melanoma (**Fig 6b** - right), whereas the fibroblast to melanocyte interaction was higher in melanoma (**Fig 6b** - right). In contrast, cSCC and BCC had more interactions between fibroblast and KC cells than in melanoma, especially the interactions with differentiating KC (**Fig 6b** - right). Overall, this analysis suggests the importance of fibroblast within the TME across all three cancer types, while highlighting that the interactions between fibroblast and T cells were negatively correlated to the level of cancer severity (from BCC to SCC and melanoma).

At L-R pair level, *MIF-CD44* was among the top two L-R pairs that changed the most between cancer types (**Fig S21b)**. This pair displayed strong interactions between macrophages with three cells types, including KC cells, fibroblast and T cells (**Fig S21c**). The canonical L-R target for immunotherapy, *CD80-CTLA4,* was also at the top most active L-R pairs, suggesting the power of this analysis pipeline (**Fig S21d**). The most significantly interacting cell-cell pairs for this pair in melanoma were fibroblast with melanocyte and fibroblast with T cell (**Fig S21e**). Overall, differential L-R pairs specific for a cancer type were enriched for key cancer-related pathways such as EMT (**Fig S21d**). The top L-R pairs associated with T cell and melanocytes are shown in **Fig S21f**. The global shortlisting of L-R pair interactions suggests the dominance of *CD44* participation, which is a druggable target (**Fig 6d**). New in-house data using different technologies (STOmics and Curio-Seeker) and external datasets confirmed the colocalisation of cells expressing the melanoma specific L-R pairs **(Fig 6e, Fig S22a).** The external dataset by Ji *et al*. confirmed SCC specific L-R pair colocalization **(Fig S22b).**

### Multiple lines of evidence suggest the important role of IL34-related antigen-presenting pathways in melanoma

Among all possible interactions between >2000 known L-R, the *IL34-CSF1R* consistently appeared in the top interacting pairs across spatial modalities and was higher in melanoma samples than in cSCC and BCC (**Fig 6a**). IL34 is a cytokine, predominantly produced by keratinocytes, whose receptor *CSF1R* activates immune cells, in particular macrophages and Langerhans cells (Wang et al., 2012; Stanley et al., 2014). High *IL34* and *CSF1R* coexpression correlates with tumor progression in lung cancer (Baghdadi et al., 2016 and 2018). To validate the interaction, we used RNAScope assay and confirmed the co-localisation of *IL34* and *CSF1R* (**Fig S23**), consistent with spatial single-cell gene expression data (**Fig 6c**). Low spatial resolution, transcriptome-wide Visium showed the colocalization of *IL34-CSF1R* in the dermis layer **(Fig 7a)**. Cells expressing the two genes were visualized on STOmics and Curio-Seeker data for melanoma samples and appeared to be in spatial proximity **(Fig 7b)**. Additionally, pathway comparisons for interacting Visium spots with those spots without interactions show enrichment of the antigen processing pathway and lipid metabolism (**Fig 7c, d**). We detected 531 genes upregulated in *IL34-CSF1R* positive spots in melanoma, and 758 genes for BCC. Only the melanoma gene list, not SCC-BCC lists, was found to show any enrichment for GO terms, including pathway terms associated with immune- (**Fig 7c**) and lipid-related (**Fig 7d**) functions, suggesting the important roles of the upregulation of the *IL34-CSF1R* interactions in melanoma.

### Spatial multimodal validations of ligand-receptor interactions highlight the role of CD44 and fibroblast

Beyond using independent transcriptomics technologies, we also validated ligand-receptor interactions based on colocalization between neighbour cells (**Fig S24**). Using our STRISH computational pipeline, we scanned through the whole tissue section and identified image tiles containing double-positive CD8+ PD-1+ immune cells and PanCK+ PD-L1+ cancer cells (**Fig S23**), (Tran et al., 2022). We next applied proximal ligation assay to detect pairs of proteins within a 20 nm distance (**Fig 7e**, **Fig S24**). We tested three L-R pairs shortlisted as described in **Fig 6**, focusing on CD44, a dominant receptor found with distinctively more common interactions in melanoma compared to BCC and cSCC (**Fig 7e**). CD44 acts as an MMP9 docking receptor that localises MMP9 to the cell surface, where it can degrade components of the ECM such as collagen to enhance tumour invasion (Yu and Stamenkovic, 1999). CD44 was reported to bind fibronectin (FN) to anchor cells to their surrounding ECM, potentially supporting invasion. CD44 interacts with fibroblast growth factor (FGF1 and FGF2) in melanoma, possibly enhancing tumour initiation and migration. Here we validated CD44-MMP9, CD44-FN1, and CD44-FGF2 interactions. The PLA signal clearly suggests the interactions occurred (**Fig 7e**). A similar approach can be used to validate more pairs. Moreover, we showed that these L-R pairs had significant prognostic values when applying to the public TCGA data (**Fig S25**), further suggesting the roles of cell-cell interactions to cancer phenotypes.

### Spatial-genetics integration of population-scale data highlights the role of T cell interactions

We next integrated spatial transcriptomics data with genome-wide association study (GWAS) data to genetically map skin cancer risk heritability to spatial cell types and domains for cSCC, BCC, and melanoma. Mapping involved linking SNPs to spatially enriched genes and testing their significance against non-associated SNPs (Song et al., 2025; refer to Methods). We used data from 10,557 cases and 537,850 controls for SCC and 36,479 cases and 540,185 controls for BCC (Seviiri et al., 2022). For melanoma data from 30,143 clinically-confirmed melanoma cases and 81,405 controls was used (Landi et al., 2020).

From Visium data, we found that spatial regions enriched with genetic association for BCC, cSCC and melanoma were localised to the epidermis. The association signal was, in some cases, specific to locations, rather than continuous or evenly distributed in the outer layer of the skin (**Fig 8a**). Cauchy aggregated significance for cell types show that top association for melanoma included melanocytes and KC differentiating. Fibroblast consistently displayed a strong association signal across melanoma and cSCC and BCC samples (**Fig 8b**). This was consistent with the spatial cell-cell interaction analysis, which suggested the important roles of fibroblast in interactions with melanocytes, keratinocytes and immune cells. Cell types most associated with cSCC and BCC cancer are KC dysplastic (more for cSCC), KC hair (more enriched for BCC), and KC cornified (similar level to BCC and cSCC, and more than melanoma), (**Fig 8b**). Furthermore, we also mapped genetic association signals to single cell resolution CosMX data (**Fig 8c**). The cells in the tumour regions (based on histological annotation) exhibited the strongest spatial heritability explained by SNPs linked to GSS genes in cancer region, while were not significant in other regions (**Fig 8c**).

Next, we identified ligand-receptor genes with significant correlation between the GSS of the L-R genes and Cauchy P-value for the cell type, suggesting possible mechanisms on how the genetic association for a spatial region or a cell type may be explained through interactions. Again, we observed strong immune interactions in melanoma, especially those involved in T cells, more compared to in SCC or BCC (**Fig 8d**). Visualisation of genetics association signals between T cells with Melanoma via *IL34-CSF1R* and *LTB-LTBR* suggests tissue regions, where the association with the melanoma were most strong (**Fig 8e**). The intersected regions with the enrichment of both L and R genes (and corresponding cell types) were at the junction of the epidermis and dermal regions, suggesting the roles of cancer and immune interaction at the invasive front. Across the whole genome, the top genetic association with spatial expression patterns is consistent with mapping key melanoma markers such as *MITF*, *TYR*, and *MX2,* suggesting that candidate genes detected here are highly relevant (**Fig 8f**). Moreover, interacting ligand and receptor genes, such as *CSF1R*, *LTB* and *LTBR,* were in the top 50 genes with the highest correlation between expression specificity for melanoma and spatial GWAS enrichment (**Fig 8f**). Top genome-wide significant SNPs are associated with those genes having the highest specificity of T cells or melanoma. This suggests that the heritability of melanoma cancer risk may be exerted from effects on T cells and melanocytes (**Fig 8f)**. Overall, our integrative analyses of spatial data with GWAS suggests functional interpretation of SNPs mapped to genes, cellular interaction and cancer regions.

### The skincanceratlas database allows users to browse gene expression and L-R data from three omics technologies

We have created a comprehensive, interactive database called skInteractive that allows users to explore our high-throughput single-cell and spatial data atlas and interactome (**Fig S1**). The Atlas section of the database shows cell type clustering and annotation results from scRNA-seq, Visium and CosMx data. The Gene Explorer section allows the user to browse genes and L-R pairs at single-cell and/or spatial resolution in select samples from cSCC, BCC and melanoma patients. No coding or data downloads are11equireed, making the **skincanceratlas** database an accessible and user-friendly way to browse this resource. The skInteractive database can be accessed at https://skincanceratlas.com.

## Discussion

We present cell types, gene signatures, and differential interactomes of the three major skin cancer types, which collectively comprise approximately 70% of all cancers in populations of European ancestry. Using 12 complementary technologies, we generated comprehensive single cell cSCC, BCC and melanoma datasets, resulting in an integrated multiomics reference atlas with complementary resolution, sensitivity, and throughput. By leveraging spatial proximity, orthogonal platforms, and multimodal integration, followed by extensive experimental and external-data validation, we constructed high-confidence interaction networks. Accordingly, this interaction atlas addresses a long-standing gap in understanding the roles of CCI in cancer progression, which remains not well characterized (Feller et al., 2016; Thieu et al., 2013). In total, 14 orthogonal technologies were assessed to provide guidelines on experimental design to address limitations and combine strengths to cross-validate and gain more biological information than is attainable through any single technology.

For cell-cell interaction at transcriptome-wide analysis, we integrated scRNA-seq and Visium to map ligand-receptor based interactions. In this combination, the scRNA-seq data become a reference for deconvolution in Visium data, allowing cell-type specific inference of L-R interactions using Visium spot-based, deconvolution data to infer both autocrine interaction within one cell or cell types and paracrine interaction between two cell types (Pham et al., 2023). Single-cell resolution spatial transcriptomics data from Xenium and CosMX platforms was then used for validating the L-R pairs predicted by Visium and scRNA-seq. In this way, we found L-R pairs and cell-type pairs in tissue niches most active in one cancer type than the other two or in cancer compared to non-cancer samples. We applied our MMCCI method that enables within and cross-sample statistical comparisons (Hockey et al., 2024). The cell-type specific L-R analysis from Visium, CosMX and Xenium data suggested that epithelial-mesenchymal transition was the main interaction pathway that differs between melanoma and cSCC-BCC, with the stronger interaction between fibroblasts and melanocytes in melanoma samples and more interaction of fibroblasts with T cells in BCC-cSCC samples. The enriched colocalization of melanocyte, fibroblast and T-regulatory cells (MKFT niches) was consistently found in CODEX and Xenium data within the melanoma community. Given that fibroblasts are essential in initiation and progression of both keratinocyte cancer and melanoma (Wang et al., 2012; Sasaki et al., 2018; Van Hove et al., 2022), the differences in fibroblast interactions between the three cancer types may contribute to differences in their metastatic potential. Increasing evidence suggests that interactions between mutated melanocytes and fibroblasts are associated with melanoma progression (Flach et al., 2011; Kim et al., 2013; Ayuso et al., 2021; Wang et al., 2016).

To discover transcriptional gene signatures of KC cancer cells, we compared cell-type specific DE analysis results from scRNA-seq data with that of CosMX, Xenium and Visium data. Reproducible markers for KC cancer and melanomas were consistent across platforms and samples. These are highly confident candidates for further experimental perturbation studies. In BCC-cSCC, comparing 11 healthy and cancer samples with scRNA-seq, we found an enrichment of the immune response with CD4 and regulatory T cells, consistently higher in cancer samples across all five patients. At the gene level, 39 genes were upregulated for cancer samples across the entire cSCC dataset, including genes related to progression like *S100A7* and *KRT6B (*Chen *et al., 2019).* Comparisons between cancer and healthy regions of each sample identified a consistent shift towards fibroblast-based signalling in skin cancer samples. With spatial data, we obtained insights into fibroblast interactions specific to partner cell types, such as melanocytes and T cells in the MKFT niches.

To find reproducible spatial patterns specific for melanoma, we mapped melanoma communities across CODEX, Xenium and Glycomics modalities and performed joint analysis of spatial omics data for this community, showing the multimodal evidence for the upregulation of tyrosine and pyrimidine metabolism pathways. The community-based colocalization analysis added more evidence for the interaction of melanoma/melanocytes with fibroblast and T cells in the MKFT niches. The ligand-receptor based analysis of the melanocyte-enriched community suggested stronger interaction in melanoma samples compared to in BCC, specifically with strong interactions involving collagen with CD44 and Integrins, mostly produced by fibroblasts, activating T cells and cancer cells respectively. Inflammatory and mesenchymal fibroblast cells and T cells create a protumorigenic microenvironment that may be associated with survival (Schütz et al., 2023; Zhang et al., 2022).

Multimodal validation of ligand-receptor interactions provided strong evidence for the L-R pairs and interacting cell types across modalities (RNA or protein) and resolution (local or nano-scale distance). Computational validation of interactions was through comparing results across platforms, where the broad discovery of interactions using scRNA-seq and Visium can be confirmed at a single cell resolution by the colocalisation of ligand and receptor signals in the Xenium and CosMX. Further validation was performed by targeted hybridization using RNAscope, with highly sensitive chemistry and single-molecule resolution. Changes in downstream signalling pathways of the L-R pairs also contributed to validations. Moreover, we extended the validation to the protein level, with Opal TSA chemistry for highly sensitive detection of protein colcalisation. However, the colocalisation detected by Opal data lacks the distance resolution to find exact interactions. We, therefore, used PLA assay to detect interactions within 20 nm distance, specifically mapping the interactions to the cell membrane. This way, we validated CD44-FGF2 interaction, which is druggable, demonstrating a pipeline for drug discovery.

Extending beyond a typical small cohort of spatial data, our integration analysis involved the mapping of genetic association signals derived from >500,000 people. We mapped SNP heritability tissue regions and showed the genetic association of melanocytes for the melanoma trait and of KC dysplastic and KC cornified for cSCC and BCC, also highlighting the roles of T cells. This analysis demonstrates that while each of the skin cancers arises from a specific type of cell, their shared and independent risk genes and SNPs act across a range of skin cells. The result suggests that one may need to consider that at least some risk loci may be mediated by cis-regulation in keratinocytes, which are involved in tightly controlling melanocyte proliferation and invasion. The analysis also provided evidence on the roles of genetics association in ligand-receptor interaction, such as the *IL34-CSF1R* pair.

Together, by using a spatial multi-omics approach with new data from 12 independent technologies and two additional technologies for validation, as well as using external datasets, our study represents the comprehensive comparison of spatial cellular signatures across the three skin major cancer types, BCC, cSCC and melanoma. Although the limitation in the number of sample size exists, the cross-validation by 14 independent experimental technologies provided high-confidence results for the 24 individuals. Further validation through integration with GWAS data of >500,000 patients strengthened generalizability. The highly integrated spatial multi omics dataset is available through our skin cancer website (https://skincanceratlas.com/), which is accessible to the broader research community for visualisation and analysis without requiring coding. The data is expected to be useful in multiple contexts, for example, by providing new insights into pathways in which GWAS DNA mutations have been reported but whose functional consequences at the single-cell and spatial levels remain poorly understood. The interacting cell types and L-R pairs identified here represent promising therapeutic targets for skin cancer treatment, including immunotherapies.

## Methods

### Patient material and ethics

All samples (**Table S1**) were collected with informed patient consent and approved for research use under ethics approval numbers 2018000165 and 2017000318 by the University of Queensland’s Human Research Ethics Committees and 11QPAH477 by the Metro South Human Research Ethics Committee. All formalin-fixed, paraffin-embedded (FFPE) blocks were previously prepared following a standard fixation procedure in 10% formalin, processed in ethanol and xylene and embedded in paraffin wax. The four melanoma samples (patients 6747-085P, 21031-08TB, 48974-2B, 66487-1A) were collected during 2008-2018 and all blocks were stored at room temperature. Full patient IDs are abbreviated as 6747, 21031, 48974 and 66487 throughout this manuscript.

For BCC and cSCC samples from eight patients (B18, E15, F21, P30, P13, P04, R01, D12), all fresh-shaved biopsies were obtained in accordance with the approved ethics protocol (11QPAH477). Patients presented at the Princess Alexandra Hospital Dermatology Department between October 2018 and February 2020. Among these patients, three patients (B18, E15, F21) were diagnosed with both cSCC and BCC and biopsies of both cancer types were collected. Five patients (B18, P30, P13, P04, R01) kindly consented to participate in the collection of 4 mm punch biopsy samples of non-sun exposed non-cancer skin for paired scRNA sequencing experiments. Lesion identity was confirmed by pathological inspection. Portions of each sample from these patients were also preserved with 10% formalin as described for FFPE samples above. To process fresh samples for scRNA sequencing, briefly, fresh-shaved biopsies were collected in DMEM for immediate tissue dissociation. Tissue was incubated in 10 mg/mL Dispase II (cat. No. 04942078001, Roche, Darmstadt, Germany) for 45 min at 37°C, snipped into small pieces with scissors, and incubated in 0.25% Trypsin for 2 min. The cells were disrupted gently with a pipette and filtered through 70 µm and 40 µm cell strainers, taken up in culture medium and spun down at 350 rcf. Resuspended cells were collected in PBS containing Foetal Calf Serum for single-cell sequencing.

scRNA-seq was performed on 14 samples representing both cancer and non-cancer cSCC and melanocyte lesions. Healthy and cSCC biopsies were paired, from patients B18, P30, P13, P04 and R01 (**Table S1**). The cancer biopsy from patient P13 was identified as being intra-epidermal carcinoma (IEC), also known as Bowen’s disease, a more superficial subtype of cSCC which occurs in the upper epidermal layer. Patient B18 was diagnosed with both cSCC and BCC, and tissue from both cancer lesions was pooled prior to library preparation. Patients E15 and F21 were also diagnosed with both cSCC and BCC, but only cSCC tissue was used for scRNA-seq. Two separate samples were collected for P30; data were pooled after sequencing. Melanoma used for CODEX, Xenium and spatial glycomics were archival samples from a retrospective patient group with thin melanomas (Stage I, Breslow depth <1mm). For snRNAseq, three archived melanoma samples representing three diagnosis types (by 23 pathologists), including malignant, intermediate (dysplastic) and benign melanocyte lesions (MPS13, MPS42, MPS43). Skin samples from healthy volunteer donors aged from 25-45 without skin cancer were collected from the forearm of the donors and were preserved in FFPE format.

Data generation, pre-processing and cell type annotation methods are described in detail for all technologies in the **Supplementary Methods**.

### scRNA-seq data analysis

scRNA-seq data was generated, processed, integrated and annotated as described in the **Supplementary Methods.** L-R analysis for scRNA-seq data was performed using CellChat (Jin et al., 2021), using normalised gene expression for all patients and all genes as input. Analysis was performed as per the detailed CellChat vignette. Circos plots were generated using the R package circlize (Gu et al., 2014). Significant L-R pairs present in ≥3 samples were visualised in a heatmap using ComplexHeatmap (Gu et al., 2016). GO analysis was performed as described above for the core gene suite analysis. L-R pairs were split into their composite genes prior to analysis.

Cancerous KC cells in cSCC samples were identified based on two intersecting criteria. Copy number variation analysis was performed using both InferCNV (Tickle et al., 2019) and CopyKat (Gao et al., 2021) using default parameters. We make use of CopyKAT’s ability to predict ‘Aneuploid’ cells and InferCNV’s de-noising and QC filtering approach to retain only the cells that are likely to be ‘Aneuploid’. Candidate KC cancer cells passed the first round of filtering if they were predicted to be aneuploid by both tools. Next, genes differentially expressed between KC cells from cancer and healthy biopsies were identified using either edgeR (Robinson et al., 2010) or Scanpy *(Wolf et al., 2018)* and used to calculate an “cSCC score” using a custom python script equivalent to Seurat’s AddModuleScore function. Cells receiving a cSCC score in the ≥95th percentile of all scores for both methods passed the second round of filtering. Therefore, KC cancer cells were those found to be both abnormal in ploidy and enriched for genes associated with cancer biopsies. KC cancer cells were compared to TSK cells from Ji et al. (2020) as described in the **Supplementary Methods.**

Melanoma cells were annotated in a similar way to the KC cells, with the module score calculated using DE genes between the malignant melanoma sample and the benign ones in the similar manner and with an ≥80th percentile threshold used for the second step of filtering based on the observed module score distribution and the number of cells.

### Visium data analysis

Data generation, processing and integration, plus cell type deconvolution and CCI analysis were performed described in the **Supplementary Methods.** For the IL34-CSF1R analysis, raw L-R scores from stLearn were used to classify spots as either IL34_CSF1R-positive (i.e. with a L-R score > 0 for this pair) or -negative (i.e. with a L-R score = 0). Integrated Visium Seurat objects for each cancer type in turn were used to perform differential gene expression analysis using Seurat’s FindMarkers function (Wilcoxon test with parameters min.pct = 0.25, logfc.threshold = 0.25, adjusted p-value threshold ≤0.05) to compare gene expression between positive and negative spots. GO enrichment analysis was performed as described above. GO terms associated with upregulated genes in melanoma were split into functionally related groups (**Fig 8f-g**) by calculating pairwise semantic similarity values between GO terms using GOSemSim (Yu et al., 2010). K-means clustering (k = 3) was used to cluster the resulting semantic similarity values into three groups of related GO terms. Genes associated with each GO term in each group were plotted using Complex Heatmap (Gu et al., 2016).

### CosMx data analysis

CosMx data was generated, processed, integrated and annotated as described in the **Supplementary Methods.** We first analysed CCI within individual FOVs. We used our SCTP method in stLearn (Pham et al., 2023) for CCI prediction, because this tool incorporates information about L-R pairs, cell types, and physical distances, thus maximising data usage and providing spatially meaningful (and therefore more biologically-meaningful) results. Briefly, for a given cell, SCTP defines a neighbourhood as the set of cells within a predefined spatial distance of that cell. For each L-R pair and each cell in an FOV in turn, L-R scores are calculated as the sum of the mean ligand expression and the mean receptor expression across a given cells’ neighbourhood. The L-R score is further corrected by neighbourhood cell type diversity, which is known to positively correlate with the likelihood of CCI (Rieckmann et al., 2017; Hou et al., 2020). stLearn uses a permutation test to determine the null distribution of L-R scores for hypothesis testing. It defines significant cells and L-R pairs. We performed cell type-specific CCI analysis to examine significant L-R interactions between pairs of cell types, using the outputs from the cell level analysis described above. Briefly, SCTP generates a CCI_LR_ matrix by counting the number of cells with significant L-R score signalling from one cell type to another for a given L-R pair. Like the cell level analysis, SCTP uses a permutation analysis to test whether these counts are significantly different from random.

To quantify the spatial heterogeneity of each FOV, we first constructed cell-cell neighbourhood networks by applying Delaunay triangulation to cell spatial coordinates, resulting in one network per FOV. Next, we applied Rao’s quadratic entropy to each cell in each network to measure the cell type heterogeneity. We elected to use Rao’s quadratic entropy scoring for this purpose because it can consider both the probability of two neighbouring cells (i.e. two cells sharing an edge) being different cell types, and the spatial distance between each member of the neighbouring pair. As the natural entropy score is often used as a measure for connected graphs, we fed the customised Delaunay cell neighbourhood network through the ATHENA local quadratic scores function (Martinelli et al., 2002). For cross-cancer type comparison, we aggregated the entropy scores of cells from the same FOVs and grouped the FOVs by cancer subtype. The entropy score was calculated as a product of the spatial distance of cells and the cell type probability; these remain as constant units across FOVs, so normalisation is not required. We performed pairwise comparison of the distribution of entropy scores from each cancer subtype using a Wilcoxon rank sum test.

### Xenium data analysis

Formalin-fixed paraffin-embedded (FFPE) tissue blocks were sectioned at a thickness of 5 μm and mounted onto Xenium slides, in accordance with the FFPE Tissue Preparation Guide (10x Genomics, CG000578, Rev B). In situ hybridisation was carried out overnight using 260 probes from the pre-designed Xenium Human Skin Panel (10x Genomics). DAPI staining was used to label nuclei, which were used for the estimation of cell boundaries (10x Genomics, CG000582, Rev D). Following completion of the run, H&E staining was conducted on the same tissue region. Each Xenium sample was preprocessed individually using Seurat version 5.0. During the quality control (QC) step, cells with zero expression across all genes were filtered out. Normalisation was performed using the *SCTransform* function, followed by principal component analysis (PCA) using the top 30 principal components. To annotate cell types for the Xenium cells, Seurat’s label transfer workflow was employed. The melanoma single-cell RNA-seq dataset processed in the previous section was used as the reference. Anchor points between the reference and Xenium datasets were identified using the *FindTransferAnchors* function. Cell type annotations were then transferred to the Xenium data using the *TransferData* function, applying the level 2 annotation labels from the reference dataset.

### CODEX data analysis

Cell segmentation for CODEX QPTIFF data was done using Cellpose as an implementation function in the Sopa package. Signal intensity for each protein channel was then mapped to the Cellpose boundaries. Outlier cells with data lower than 0.05 quantile or higher than 0.95 quantile were removed from the raw protein expression intensity matrices. The data was transformed with arcsinh and scaled to mean 0 and standard deviation 1. The cell type identification was performed based on the protein markers included in the panel using z-scores. For spatial community analysis (niche detection), we used two methods, NeighbourhoodCoordination and MonkeyBread neighbourhood clustering. Both methods clustered cells based on the cell type proportion of a neighbourhood tissue area as squared tiles (windows) or a circle of a given radius. Colocalization between cells of two cell types was computed based on distance. A network connecting cell types and communities was drawn using the network approach in the Sopa package.

### Spatial Glycomics data analysis

Following general pre-processing, data from three MALDI samples were individually analysed using the R-based package *SpaMTP V1.0* (Causer et al., 2024). Mass peaks were initially binned at a resolution of 250 ppm, resulting in 5433 detectable m/z peaks. Samples were annotated against the Lipid Maps database implemented in *SpaMTP*, with the *AnnotateSM* function. Principal component analysis was run and dimensionality reduction was performed for each sample using the first 30 principal components. Louvain clustering was implemented using a resolution of 0.3, resulting in samples containing between 9 and 13 clusters. Pseudo-bulking differential metabolite abundance analysis was performed per cluster using the *SpaMTP FindAllDEMs* function. The top 10 m/z values per cluster, per sample, were then combined and hierarchical clustering was implemented to group similar clusters together based on pseudo-bulked expression. Melanoma clusters were identified based on spatial location and confirmed by hierarchical grouping. Differential abundance analysis was again performed to identify all significantly abundant metabolites within the melanoma cluster compared to all other clusters of each sample. Common metabolites that were differentially abundant within the melanoma cluster, across all three samples, were then identified and spatially plotted across each tissue sample.

### Integrative Cell Neighbourhood/Community analysis

For each data type, Visium, Xenium, and CosMX, we applied the NeighbourhoodCoordination method to map communities of nearby cells that had similar neighbourhoods as assessed based on cell type composition (Schürch et al., 2020). A neighbourhood matrix, where the proportion of cell types within a neighbourhood (window) were calculated for each cell in each Visium sample, or Xenium sample or CosMX FOV using the same setting. The windows for all samples and FOV within one technology (e.g., from all Visium samples) were merged into one matrix per technology. The windows were then clustered using K-mean clustering, with K=10 for all samples, for consistency.

To find robust communities, we combined together 30 communities, representing Visium, Xenium, and CosMX. Each community was represented by the proportion of cell types within the community. The matrix of combined communities was used to group similar communities into functional categories (considered as meta communities) such as tumour, stromal, immune or KC. This way, communities across cancer types and platforms can be compared. For example, we compared the cancer community (e.g., CosMX_6) in cell-cell interactions across BCC, cSCC and Melanoma samples. We also compared the cancer heterogeneity at community level (e.g., CosMX_6 and Visium_2 for the cancer sample), where each community may be more or less abundant in one cancer type compared to the other cancer types.

### Integrative Cell-Cell interaction analysis

We implemented our MMCCI cell cell interaction analysis pipeline to integrate data from multiple samples and multiple platforms. MMCCI takes inputs as CCI results from individual samples calculated by spatially-aware interaction scores and P-values from stLearn. Prior to integration, the interaction scores were normalised to take into account differences in the number of cells/spots across samples. The integration process resulted in two main outputs, the strength of interactions between two cell types (total number of interacting cells) and the integrated P-value for the interaction (using Stouffer’s method to calculate the inverse cumulative distribution of all P-values).

### Integrative DE and pathway analysis across four platforms scRNA-seq, Visium, CosMX and Xenium

Pseudobulking following EdgeR DE analysis pipeline with quasi likelihood ratio test were applied across platforms. Shared DE genes, consistently found in all orthogonal spatial technologies, were highly confident DE genes that can be considered as promising markers of a cancer type or a biological pathway differentially regulated among cancer types. For each modality, significant abundant analysis was performed between clusters/spatial communities. Using cell type annotation, a spatial community/neighbourhood commonly found across modalities can be jointly analysed.The differential markers (genes, proteins or metabolites) derived from comparing one community with the remaining other communities were input into MetaboAnalyst for joint pathway analysis.

### Spatial Datasets with single-cell resolution (STomics and Curio-Seeker)

Some of the interacting L-R pairs were visualized on high resolution spatial transcriptomics data. STOmics and Curio-Seeker (Now acquired by Takara Bio, USA) are both spatial technologies offering a single-cell resolution. Processed data for one melanoma sample from Curio was obtained from the company. The STOmics data was generated in-house, where one melanoma and two colorectal cancer samples (FFPE tissues) were profiled by Stereo-seq OMNI technology. The data was processed using the SAW pipeline and the counts data was used for visualizing cells expressing the L-R genes.

### Spatial multiomic validation with Proximity Ligation Assay (PLA), Opal Polaris and RNA scope

We assessed multiple approaches to validate ligand-receptor interactions, including RNAscope, Opal Polaris and Proximal Ligation Assay for validating cell-cell interactions. In the case of IL34-CSF1R, antibodies for IL34 were not readily optimised and so we applied RNAscope to examine the colocalization at single molecule and single cell resolution (**Fig 7**). Similar to Xenium or CosMX assay in resolution, the RNAscope produces an additional advantage with the Z-probe amplifier chemistry leading to a high detection sensitivity. Our data show consistent results between RNAscope and CosMX data that prove the colocalization within neighbouring cells of IL34 and CSF1R mRNA (**Fig S23, S24**). Using spatial transcriptomics data, we can also validate downstream pathways that change specifically associated with the L-R pairs co-expressed in the spatial spots. For example, the functional downstream consequences of IL34_CSF1R signalling were identified based on genes that were differentially expressed between IL34_CSF1R-positive (L-R score >0) and -negative (L-R score of 0) spots for each cancer type (**Fig 7a, c-d**).

At the protein level, we detected colocalization of the protein pairs with Vectra Polaris and Proximal Ligation Assay. The PLA was performed to validate ligand-receptor interactions identified through spatial transcriptomics data. FFPE melanoma tissue sections were deparaffinized, rehydrated, and subjected to antigen retrieval. Primary antibodies specific to the ligand and receptor of interest were applied and incubated overnight at 4°C. Following the manufacturer’s instructions for the NaveniFlex Tissue MR Atto647N kit (Navinci). Fluorescent signals indicative of close-proximity interactions were generated through ligation and amplification steps. After PLA single is generated, the tissue sections were stained with anti-mouse/rabbit secondary antibodies conjugated to Alexa Fluorophore for 1 hour at room temperature for visualisation of target proteins. Subsequently, tissues were counterstained with DAPI, and imaged using the STELLARIS Confocal Microscope (Leica). Our PLA results clearly showed the specific signals on the cell membrane in the positive control (E-cadherin and b-Catenin) and no signal in the negative control (CD31-AQP1), (**Fig S24**).

### Integrating spatial transcriptomics with GWAS data

We used the summary statistics from a cSCC GWAS study of 10557 controls and 537850 controls and for BCC GWAS with 36479 cases and 540185 controls (Seviiri et al., 2022) and Melanoma GWAS with 30,143 clinically-confirmed melanoma cases and 81,405 controls (Landi et al., 2020). We applied gsMAP method to map genetic signals to spatial gene expression in the three skin cancer datasets (Song et al., 2025). First, tissue domains were determined by finding similar spots/cells using a graph attention autoencoder network which generated latent representations, which in turn were used to find pairwise cosine similarity between spots/cells. Next, gene specificity scores (GSS) for each spot were computed by aggregating information between similar spots/cells (domain), and rank enrichment information for top abundant genes from its homogeneous spots. The expression specificity of a gene within a focal spot was assessed by calculating the geometric mean of its expression rank across the tissue region (microdomain identified by graph attention, or cell type) of the focal spot, divided by the geometric mean of its expression rank across all spots in the ST data. Genes with a ratio higher than 1 and expressed in more spots/cells in the region/cell-type than in overall sample(s) were considered specific for the region or cell type. A high GSS score for a gene suggests that the gene was higher/enriched for the region/cell-type than most other genes in that region/cell-type.

Based on proximity to the nearest transcription start sites, GWAS SNP are assigned to GSS genes for each focal spot/cell. Given the set of assigned SNPs to each spot/cell, the SNP effects in the GWAS summary statistics for BCC, cSCC, and cutaneous melanoma and LD scores (from the 1000 Genomes Project Phase 3) were used for LD score regression analysis. Given the total set of SNPs assigned to a spot/cell, the SNP-level trait heritability (Chi-square association with the skin cancer trait of a SNP from the summary statistics) is partitioned into the SNP effect of a focal spot or cell and the effect of the SNP given the baseline SNPs that are not assigned to GSS genes. The enrichment p-value is calculated based on the partitioned regression coefficient, using one-sided Z-test for bigger than 0. To compute p-value for the association of a spatial region or a cell type across the whole sample, we aggregated P values of individual spots/cells within the spatial regions (or cell types) using Cauchy combination.

### Skincanceratlas database

We built the skInteractive database in the form of a visualisation dashboard with a Shiny v1.7.2 application (Sievert et al., 2020). The database has two main sections, the Atlas, which shows cell types and clustering results, and the Gene Explorer, which allows the user to browse gene expression and/or L-R interaction scores for the different datasets and modalities. The Atlas dashboard was constructed using Javascript (NuxtJS framework). We converted all plots to geo map components in Apache Echarts v5.3.1 (Li et al., 2018) that provided interactive features to work with the plots. For the Gene Explorer Shiny app, we implemented multiple tab themes and used Seurat v4.1.1 (Hao et al., 2021) to generate plots from the different data sets stored in SeuratObjects. The Shiny application included Visium (gene expression and L-R scores), CosMx (gene expression) and scRNA-seq data (gene expression). skInteractive Database will be ported to AWS cloud.

## Data availability

All of the sequencing data and accompanying H&E images for spatial transcriptomics both raw and processed will be deposited to ArrayExpress repository (https://www.ebi.ac.uk/arrayexpress/) and made publicly available according to human ethics regulations. All other experimental data (e.g. imaging data using RNAscope or by Polaris immunofluorescence) will be made available upon request. GWAS data used in this analysis is available as detailed in the relevant publications (Landi et al., 2020; Seviiri et al., 2022)

## Code availability

The code to reproduce analyses and figures presented in this paper is available at https://github.com/GenomicsMachineLearning/SkinCancerAtlas/tree/main

The public Github repository of the skInteractive database can be accessed at https://github.com/GenomicsMachineLearning/SkinCancerAtlasBackend and https://github.com/GenomicsMachineLearning/SkinCancerAtlasWebSite and users can explore the interactive functionalities without the need for customised coding at https://skincanceratlas.com/

## Author contributions statement

Q.N. conceived the project. Key contributors to data analysis include L.G. (scRNA-seq and Visium data), P.P. (scRNA-seq), F.Z. and G.N. and O.M (CosMx data), X.J. (scRNA-seq and Visium data), M.T. (Polaris, RNAscope, and CosMx data), O.M. (GeoMx data), E.E.K (CosMx data), M.T.G. (CosMx data), Z.R. (CosMx data). D.P., L.G., P.P. and A.N. built the skInteractive web tool. A.X., S.M.T, T.V., K.D., L.P., A.K., M.L., S.R.M., S.E.W., Y.K. conducted the experiments and generated the data. M.Z., H.S., J.G.C, S.K., H.N.L., H.P.S., M.T.L, K.M.B, M.M.I, M.S, M.H.L, I.F., Y.K., M.S.S., K.K., Q.N. contributed to data analysis and interpretation. P.S., M.S.S., K.K. and I.F. provided clinical samples. All authors contributed to writing the manuscript and have reviewed and approved the manuscript.

## Acknowledgements

This work has been supported by the Australian Research Council (ARC DECRA grant DE190100116 to Q.N), National Health & Medical Research Council (NHMRC Project Grant 2001514 to Q.N), NHMRC Investigator Grant (GNT2008928 to Q.N). We thank the staff at the University of Queensland sequencing facility and the School of Biomedical Science imaging facility for their support in spatial transcriptomics and single cell RNA sequencing. We thank all members in Nguyen’s Genomics and Machine Learning Lab for discussion and support in data generation and analysis. Acknowledgements for the melanoma meta-analysis data can be found in the supplemental data.

