## Supplementary Material for "Integrating 12 Spatial and Single Cell Technologies to Characterise Tumour Neighbourhoods and Cellular Interactions in three Skin Cancer Types"

^7^ Bruker Spatial Biology, NanoString^®^ Technologies, Seattle WA 98109, USA (EEK - Department of Laboratory Medicine and Pathology, University of Washington School of Medicine, Seattle, WA, USA; ZR - Paul G. Allen Research Center at Swedish Cancer Institute; MTG - Flagship Biosciences; LP - Duality Biologics; ML - Max Planck Research Unit for Neurogenetics; SRM - Mount Sinai, Center for Disease Neurogenomics, Department of Genetics and Genomic Sciences)

*^8^* School of Agriculture and Food Sciences, The University of Queensland, QLD 4072, Australia

^9^ Division of Cancer Epidemiology and Genetics, National Cancer Institute, National Institutes of Health, Bethesda, MD, USA

^10^ Leeds Institute for Data Analytics, University of Leeds, Leeds, UK

*^11^* UMPC Hillman Cancer Center & School of Public Health, University of Pittsburgh, Pittsburgh, PA, USA

^12^ School of Chemistry and Molecular Biosciences, The University of Queensland, Brisbane, QLD 4072, Australia

^13^ Australian Infectious Diseases Research Centre, School of Chemistry and Molecular Biosciences, The University of Queensland, Brisbane, QLD 4072, Australia

^14^ Department of Emergency Medicine in Linköping, and Department of Biomedical and Clinical Sciences, Linköping University, Linköping, Sweden

^15^ Department of Dermatology and venereology in Östergötland, and Department of Biomedical and Clinical Sciences, Linköping University, Linköping, Sweden^16^ Institute for Biomedicine and Glycomics, Griffith University, Gold Coast, QLD 4215

^17^ Sullivan Nicolaides Pathology, Bowen Hills, Qld 4006, Australia

^18^ QIMR Berghofer, Population Health program, Brisbane, QLD 4006, Australia

^19^ School of Biomedical Sciences, Faculty of Health, Queensland University of Technology, Brisbane, QLD, 4001, Australia

^20^ School of Biomedical Sciences, Faculty of Medicine, University of Queensland, Brisbane, QLD, 4072, Australia

^21^ Pfizer, 21621 30th Dr SE, Bothell, WA 98021

^#^co-first authors

### **Supplementary Figures**

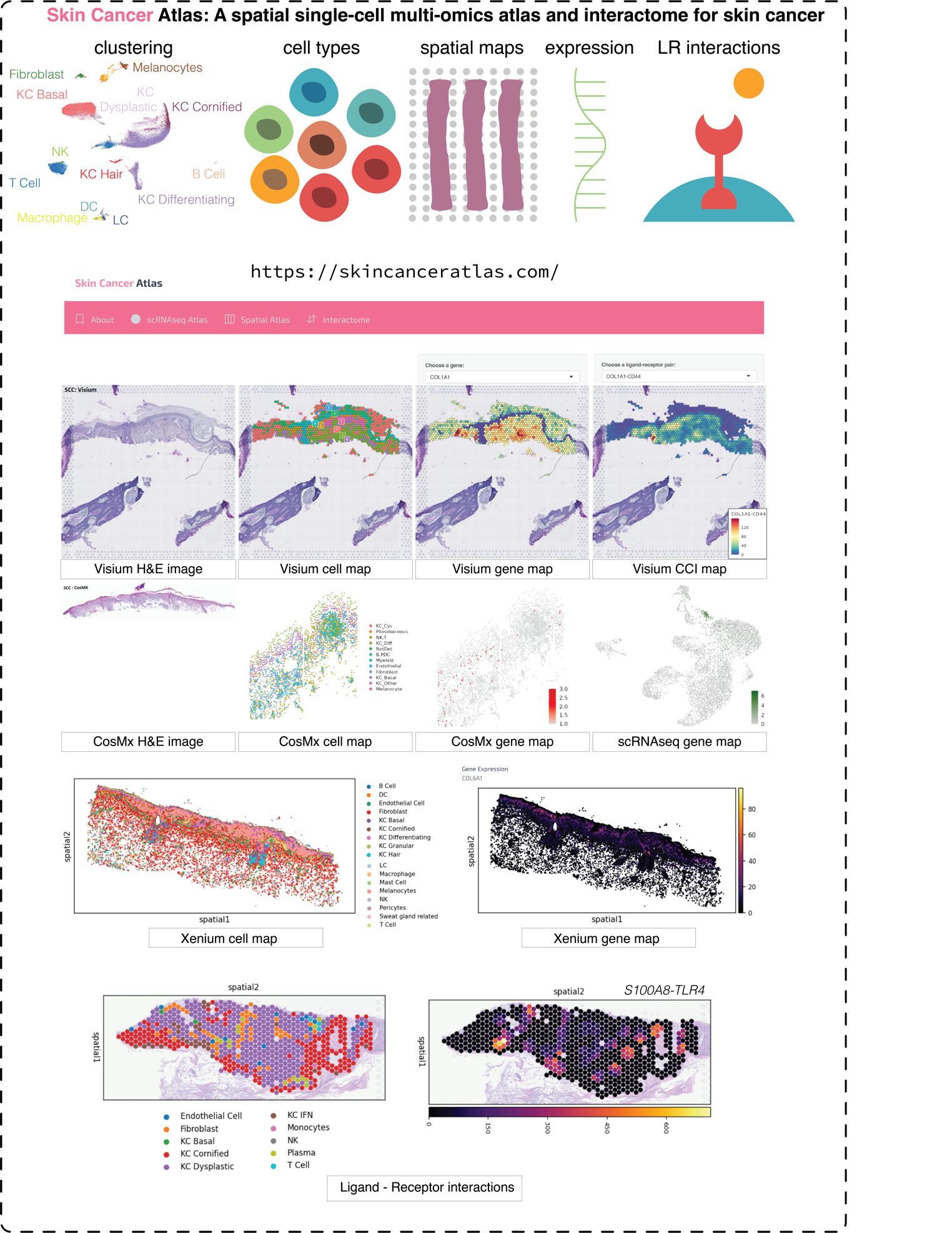

**Figure S1. The skincanceratlas database allows interactive analysis of genes, cells, interactions across the multi-omics skin cancer data resource.**

*The skincanceratlas function allows users to visualise clustering, cell type and spatial data for cSCC, BCC and melanoma. The Gene Explorer function (bottom) allows visualisation of gene expression and LR interactions at single cell and/or spatial resolution from data generated with Visium, CosMx and scRNAseq for all three skin cancer types.*

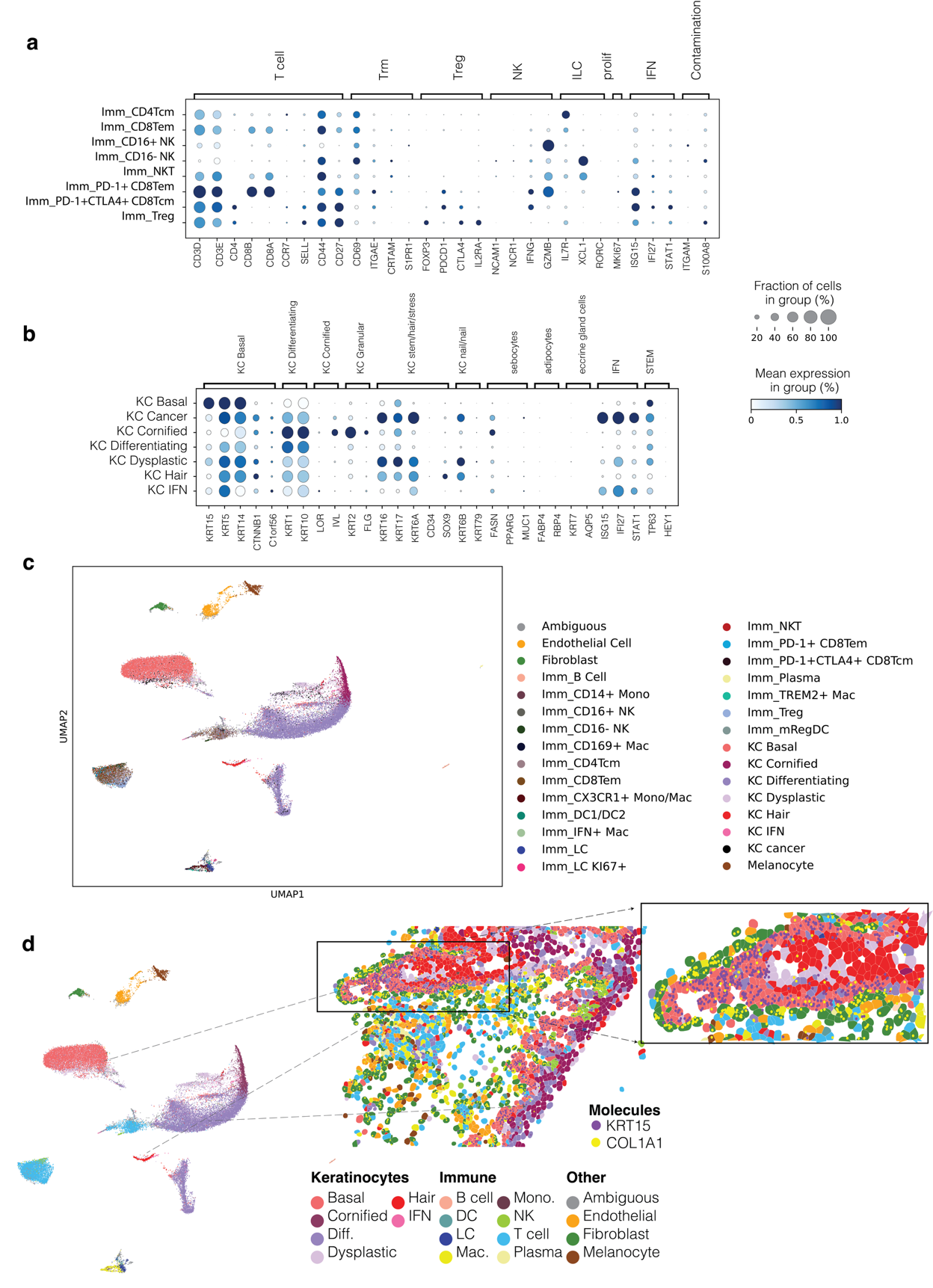

**Figure S2. Annotation of KC and Immune cells in BCC-cSCC.**

*(a) Marker expression for the level 3 annotation of immune cells.*

*(b) Marker expression for the level 3 annotation of KC cells.*

*(c) UMAP displaying the overall level 3 annotation of all 30 cell types.*

*(d) Visualisation of the single cell (sub)clusters that are mapped to the tissue using the CosMX data. scRNAseq data was used as the reference to transfer cell annotation labels to CosMX cells using the RCTD deconvolution method. The KC basal, KC hair, KC differentiating, KC cornified clearly form distinct layers in the cSCC skin sample B18. The cell types defined in scRNAseq data clearly mapped to distinct anatomical layers of the skin, providing cross-platform evidence for correct cell type annotation of the scRNAseq reference data, which did not have a spatial context.*

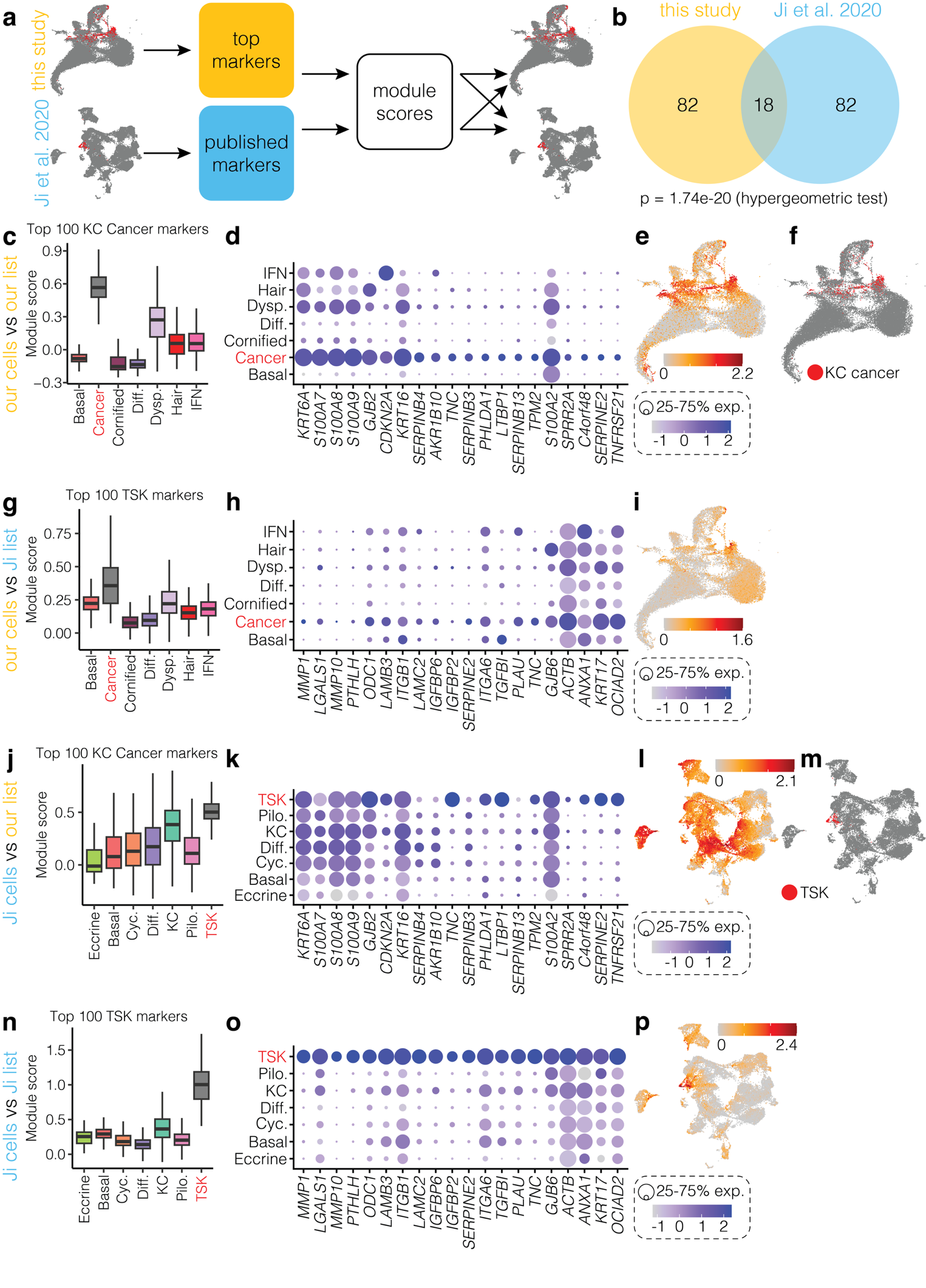

**Figure S3. Cross-dataset comparison of KC cancer cluster with published TSK keratinocytes.**

*(a) Schematic of the analysis workflow. Marker genes distinguishing KC cancer cells from other KCs in this study were identified and used to generate module scores across keratinocytes in both our dataset and the Ji et al. (2020) KC dataset. Conversely, the top TSK markers reported by Ji et al. were used to generate module scores in both datasets.*

*(b) Overlap between the top 100 markers of the KC cancer cluster and the TSK cluster; significance of the overlap was assessed by hypergeometric test.*

*(c-e) Expression of KC cancer marker genes in our KC dataset. shown as (c) a boxplot of module scores (top 100 markers), (d) a dotplot (top 20 markers), and (e) a UMAP of module scores (top 20 markers).*

*(f) UMAP plot of our dataset highlighting the KC cancer cluster in red.*

*(g-i) As in (c-e), expression of TSK markers in our dataset.*

*(j-l) As in (c-e), expression of KC cancer marker genes in the Ji et al. dataset.*

*(f) UMAP plot of the Ji et al. (2020) dataset highlighting the TSK cluster in red.*

*(n-p) As in (c-e), expression of TSK markers in the Ji et al. dataset.*

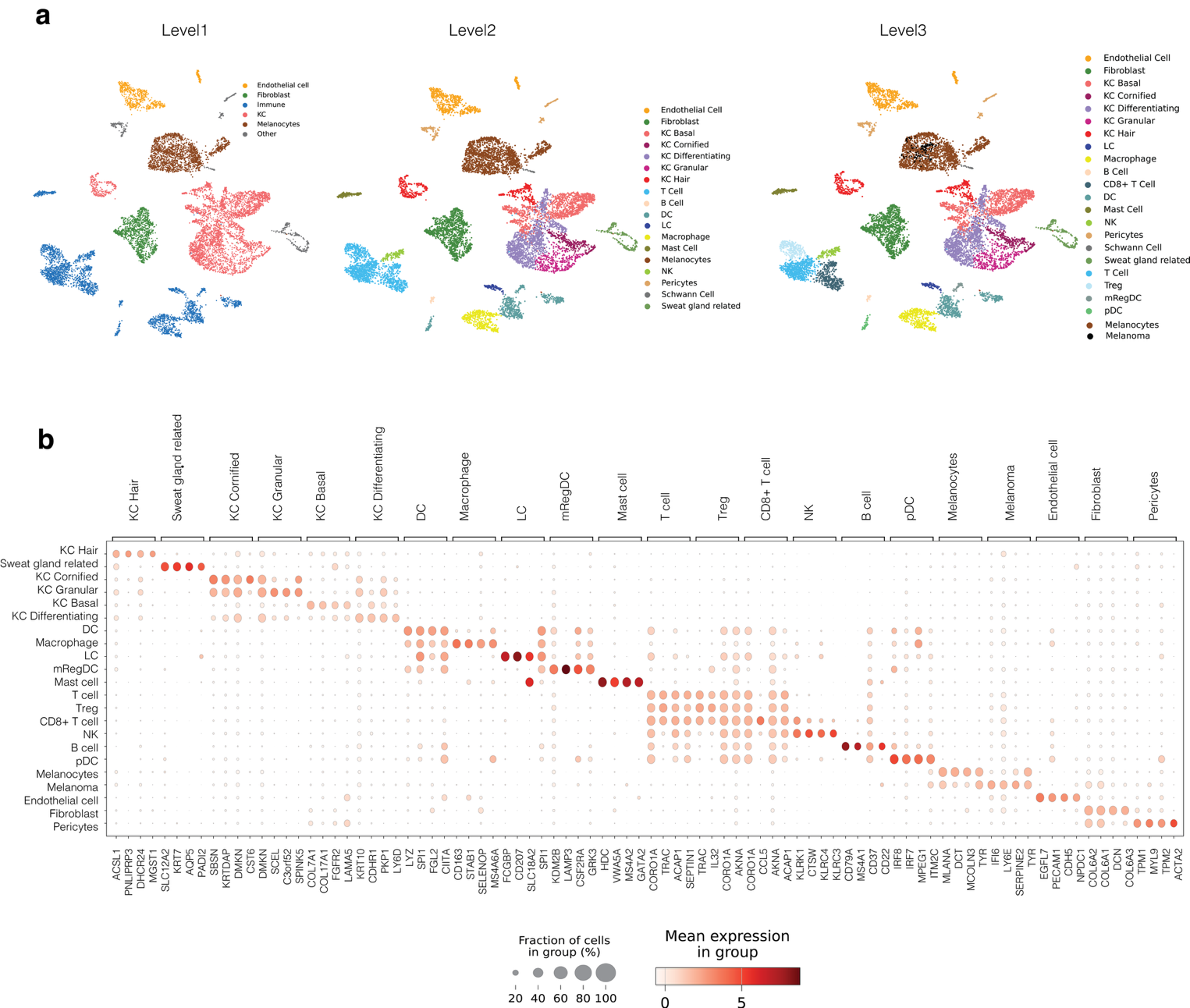

**Figure S4. Annotation of melanoma samples.**

*(a) UMAP shows the annotation of all the three annotation levels from common to specific cell types. UMAP showing Level 3 annotation also displays melanoma cells, which were not defined by Level 1 and Level 2 annotation.*

*(b) Heatmap showing marker expression for the level 3 annotation.*

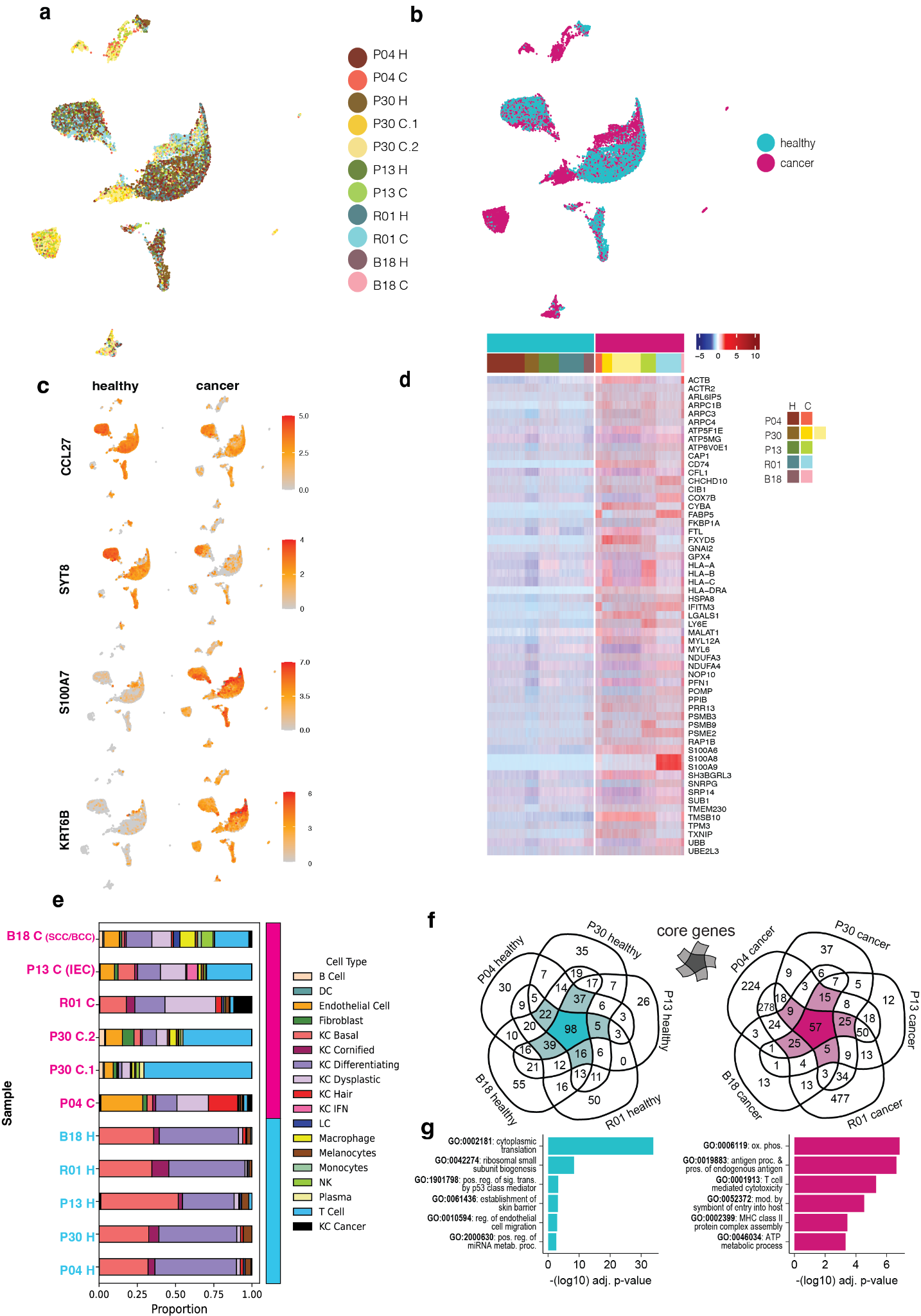

**Figure S5. A single-cell atlas of cSCC cell types.**

*(a-b) UMAP plots showing the integration of ~46,000 cells from 11 samples of five cSCC patients, showing cell distribution by patient sample (a) and cancer status (b).*

*(c) Expression of selected genes that are amongst the top 10 most differentially expressed genes between cancer and healthy cells when compared across the entire dataset. Each column shows only the healthy (left) or cancer (right) cells split from the full UMAP plot shown in (a-b).*

*(d) Heatmap showing expression in healthy and cancer samples of the 57 core cancer genes shared between all five patients (as in Panel F).*

*(e) Bar plot showing the distribution of each cell type across eleven samples.*

*(f) Genes overexpressed in healthy (left) or cancer (right) cells across individual patients. Core suites of genes that are detected across four or more of the five patients are highlighted in the centre of each diagram; suites contain a total of 217 healthy and 136 cancer genes respectively.*

*(g) The top six non-redundant gene ontology terms enriched in the two gene suites, ranked by -log10 adjusted P-value.*

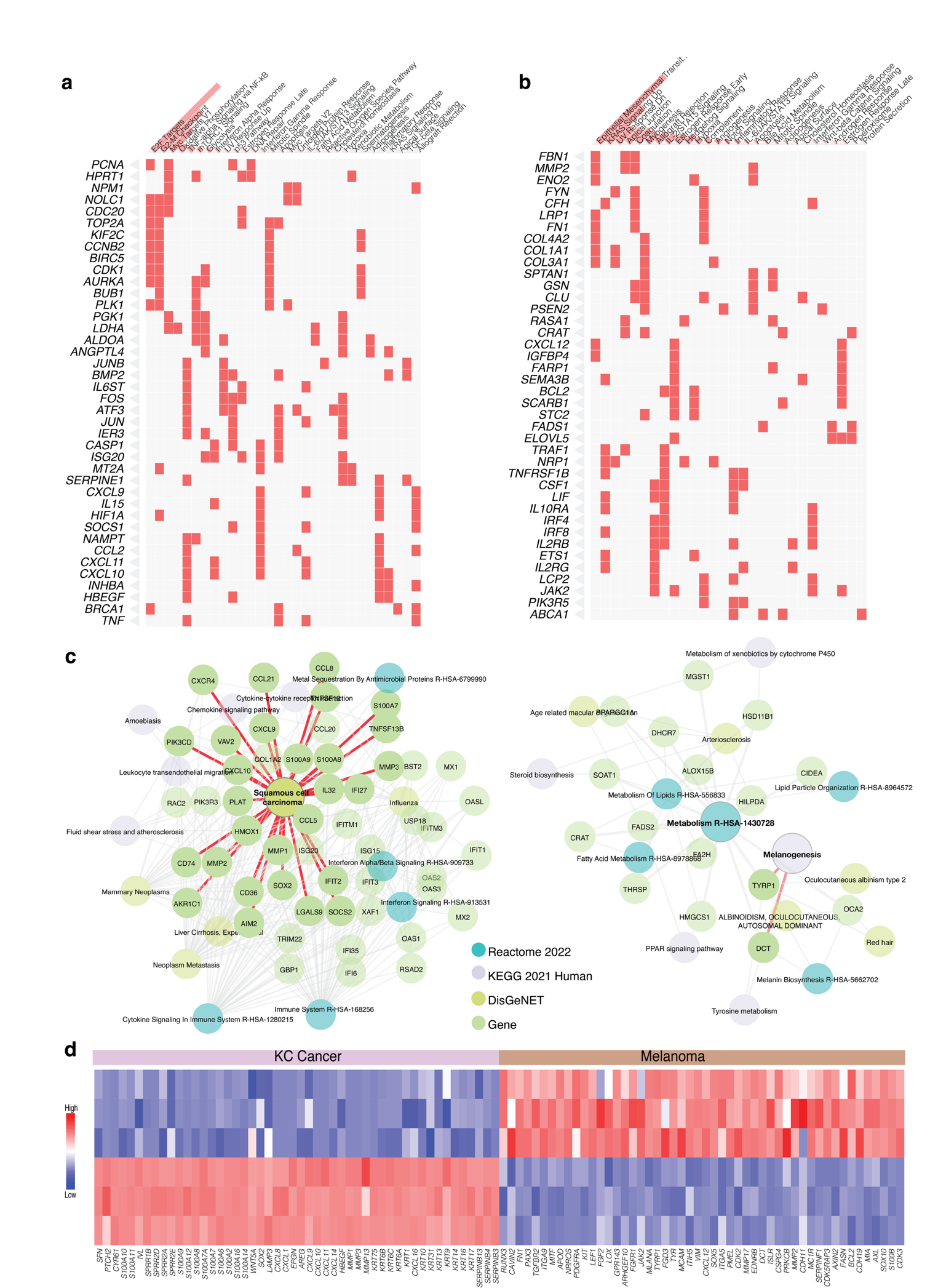

**Figure S6. Pathway analysis for gene markers of cSCC-BCC and melanoma.**

*(a) Heatmap showing enriched pathways tested against the MSigDB. All DE genes upregulated in Cancer cells in cSCC-BCC compared to normal KC cells (left) and in melanomas compared to normal melanocytes were used for enrichment analysis. Pathway names are shown in columns and genes in these pathways are shown in rows. The red bars indicate -log(P-value) and the terms are sequentially ordered with the most significant pathway on the left.*

*(b) MSigDB pathways for 2713 genes up in Melanoma Vs KC Cancer.*

*(c) Integrated pathway analysis, using KEGG and Reactome databases for genes higher in cancer KCs compared to the normal KCs in cSCC-BCC samples observed in scRNA-seq (left; 176 KC Cancer vs normal KCs) and for genes higher in malignant melanocytes compared to the normal melanocytes in snRNAseq Melanoma samples (68 genes up in Melanoma Vs normal melanocytes).*

*(d) DE genes from comparing KC cells and Melanocytes. The rows represent pseudo-bulked gene expression values per scRNAseq cluster.*

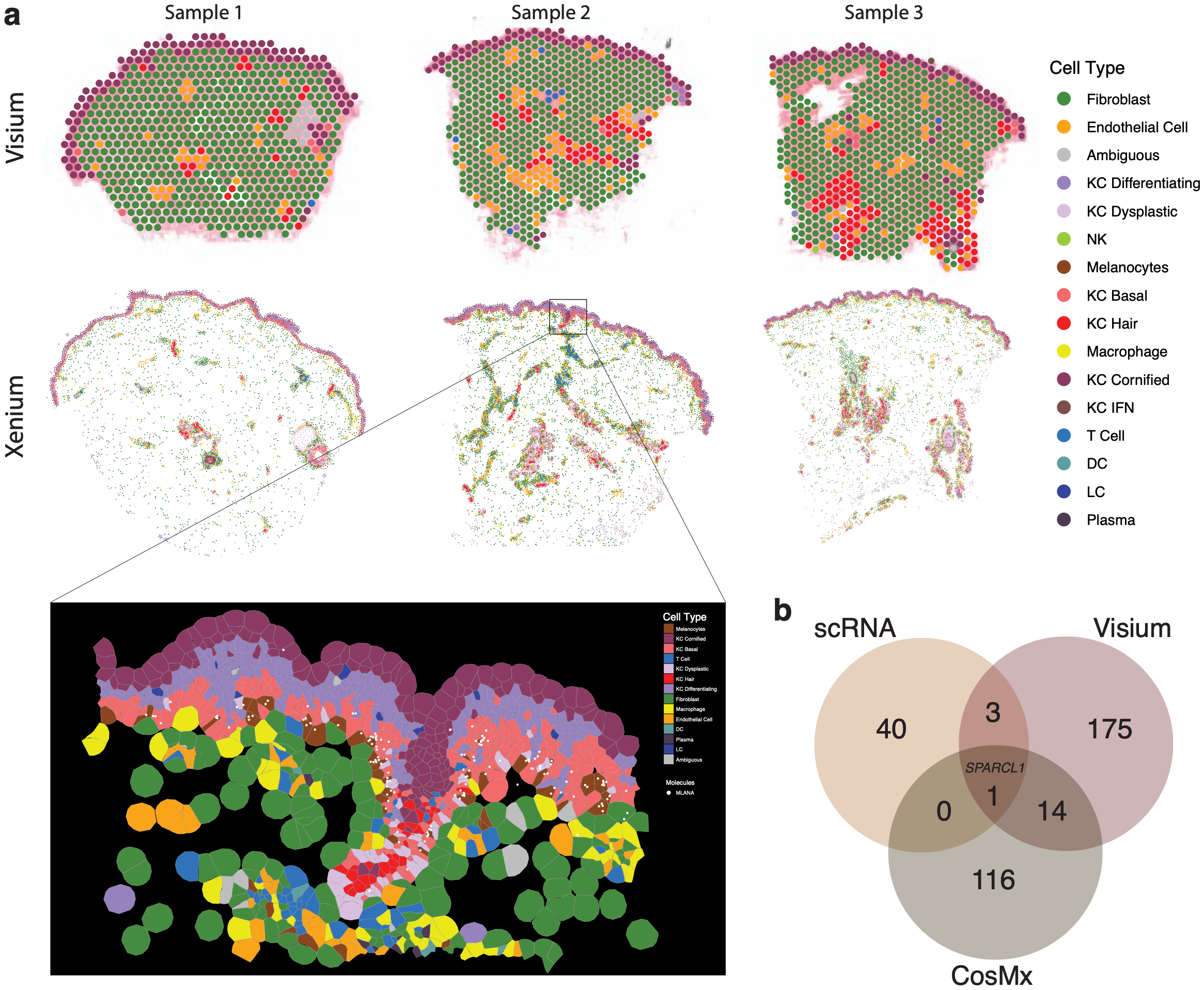

**Figure S7. Cell type annotation for the Visium and Xenium data of samples from three normal control donors.**

*(a) The donors did not have skin cancer. The top and middle panels show Visium and Xenium data of adjacent tissue sections, respectively, and a zoom-in view of a Xenium FOV is shown at the bottom panel. The legend colors indicate cell types.*

*(b) Genes significantly upregulated in Melanoma across different platforms. The three lists are derived from three comparisons: (i) Confident melanoma cells vs an equal number of normal melanocytes in scRNA-seq data (ii) Melanocytes from cancerous and normal Visium samples (iii) Melanocytes from CosMx Melanoma patients vs normal Xenium samples.*

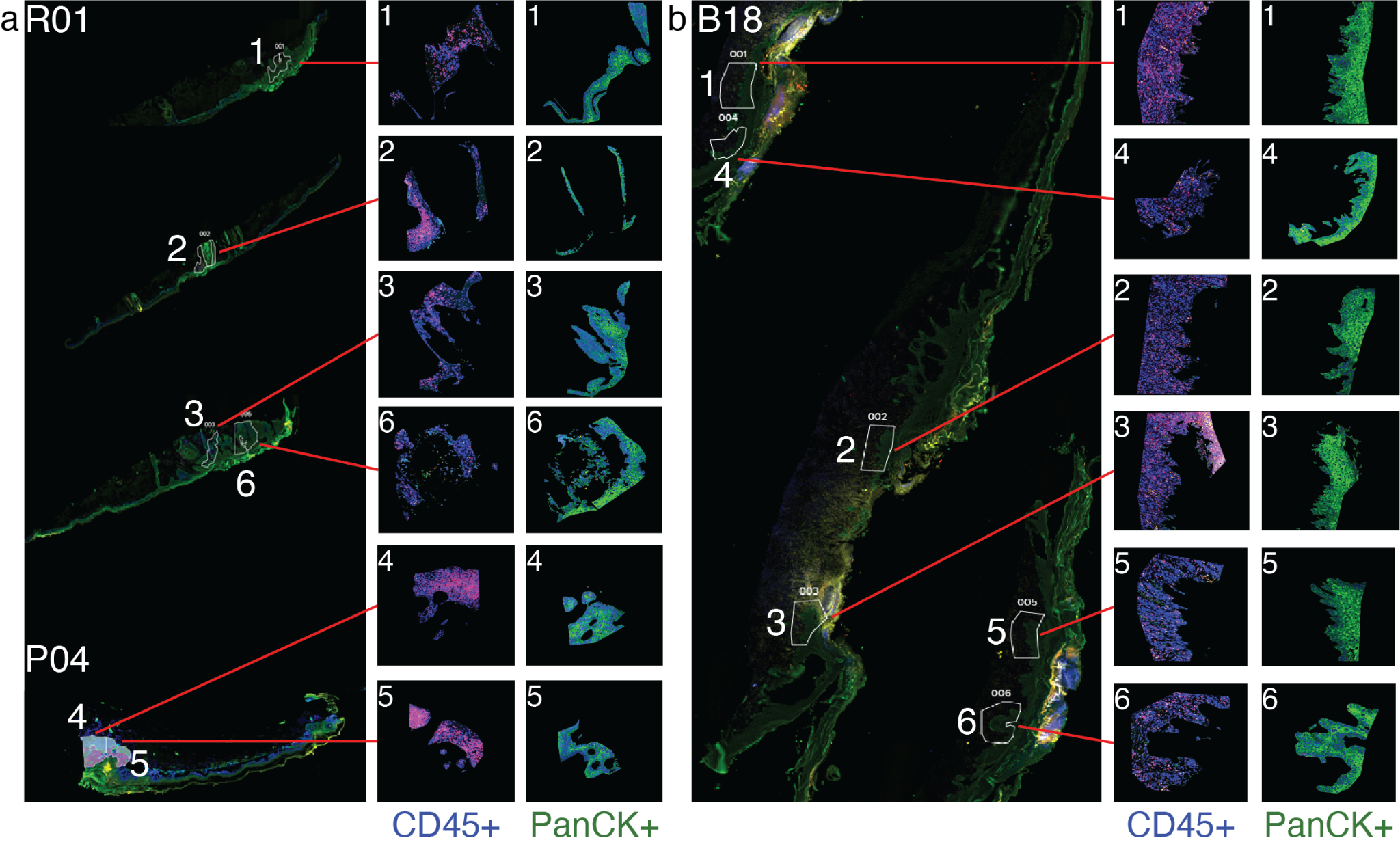

**Figure S8. Captured regions of interest for GeoMx protein quantification.**

*Morphology imaging of cSCC biopsies for GeoMx protein data, from patients R01 (a, top 3 tissues), P04 (a, bottom tissue) and B18 (b). Boxes show zoomed-in regions of interest (ROIs) selected for marker quantification, connected to their location in the full tissue image by red lines. Morphology markers show DNA (blue, DAPI), CD3 (red), CD45 (yellow) and PanCK (green). ROIs were segmented by either CD45 or PanCK expression*.

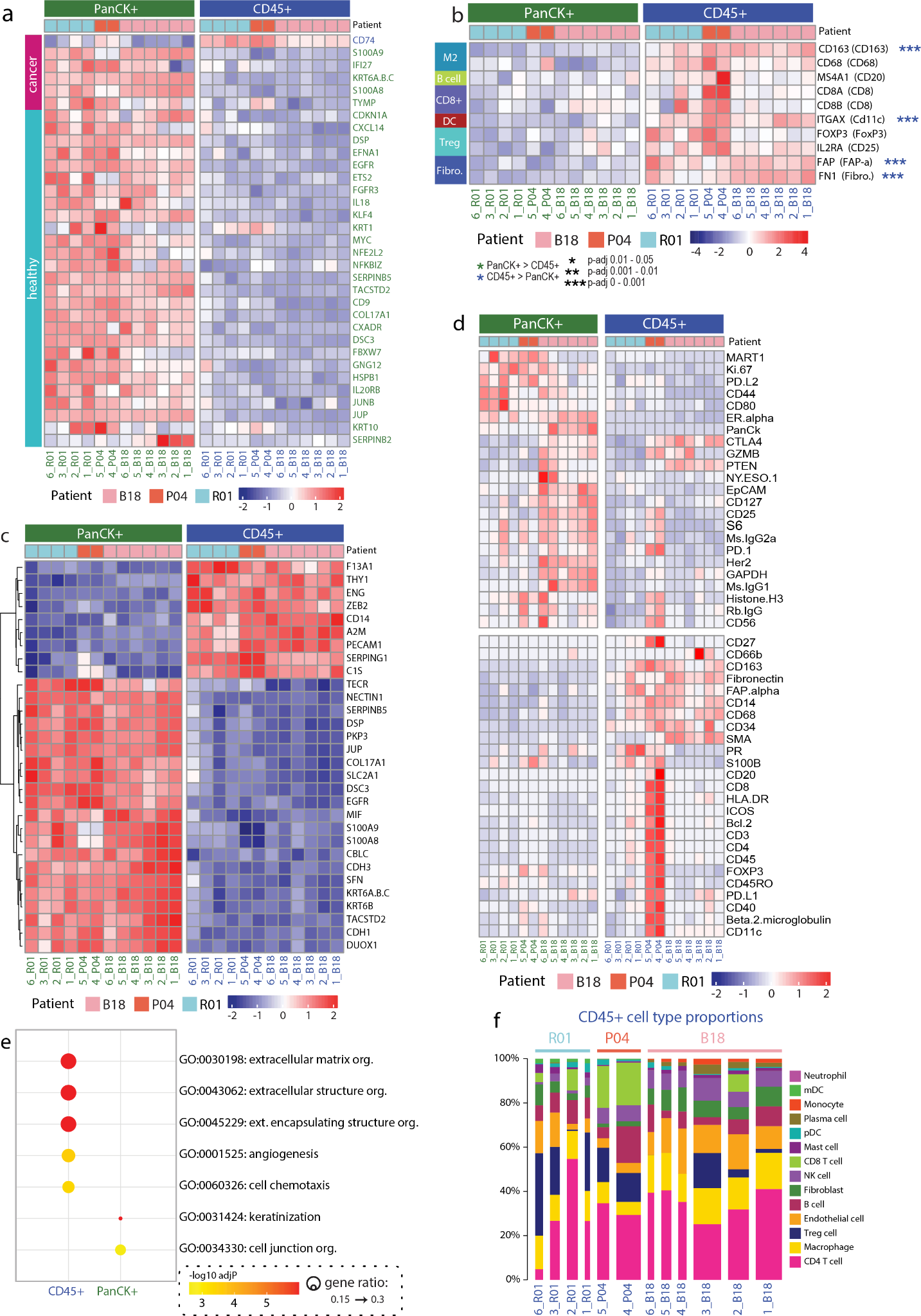

**Figure S9. Analysis of GeoMx gene markers and cell type compositions using GeoMX RNA and protein assays.**

*(a) Heatmap of 33 genes which were differentially expressed between PanCK+ and CD45+ segments within the GeoMx RNA dataset, and were also identified as members of the core cancer- or healthy-associated gene suites. Column annotations separate samples by GeoMx segment (PanCK+ and CD45+ segments) and patient, while row annotation indicates whether genes were present in the cancer- or healthy-associated gene suites from* ***Fig S4f****. Gene label colour indicates whether the gene was upregulated in the CD45+ (blue) or PanCK+ (green) segment in the GeoMx analysis.*

*(b) Validation of the presence of key cell types from our scRNASeq atlas, using RNA markers captured by GeoMx; this plot is complementary to* ***Fig S10n*** *which shows the same comparison for equivalent protein markers. Selected markers are CD163 and CD68 (M2 markers), MS4A1 (B cell marker), CD8A and CD8B (CD8+ T cell markers), ITGAX (dendritic cell marker), FOXP3 and FOXP3 (Treg markers), and FAP and FN1 (fibroblast markers). Corresponding protein markers for each gene are given in brackets. Top annotation bars separate samples by segment (PanCK+ and CD45+ segments) and patient. Marker association with particular cell types was taken from NanoString product information. Asterisks indicate markers that were differentially expressed between CD45+ and PanCK+ segments; markers that were not found to be differentially expressed are included as their expression provides evidence for the presence of these cell types.*

*(c) Top 30 differentially expressed genes between PanCK+ and CD45+ segments within the GeoMx RNA dataset. A total of 267 differentially expressed genes were identified between segments. Differentially expressed genes were sorted by FDR and the top 30 genes were selected regardless of whether they were enriched in the PanCK+ or CD45+ segments.*

*(d) Expression of all the 48 proteins (Immuno-oncology panel) in GeoMX proteomics data. The heatmap shows normalised data across FOVs.*

*(e) Top 5 GO enrichment results per segment from an analysis of the top 100 differentially expressed genes associated with the CD45+ and PanCK+ segments of the GeoMx RNA data. Differentially expressed genes were sorted by absolute fold-change values and the top 100 genes were selected from each of the CD45+ and PanCK+ segments.*

*(f) Inferred cell type proportions of ROIs from patients B18, R01 and P04 in the CD45+ segment of the RNA GeoMx data. RNA expression information was used to perform cell type deconvolution for the segmented ROIs. Bar width is proportional to the number of nuclei per ROI. Top annotations indicate the sample of origin for each ROI.*

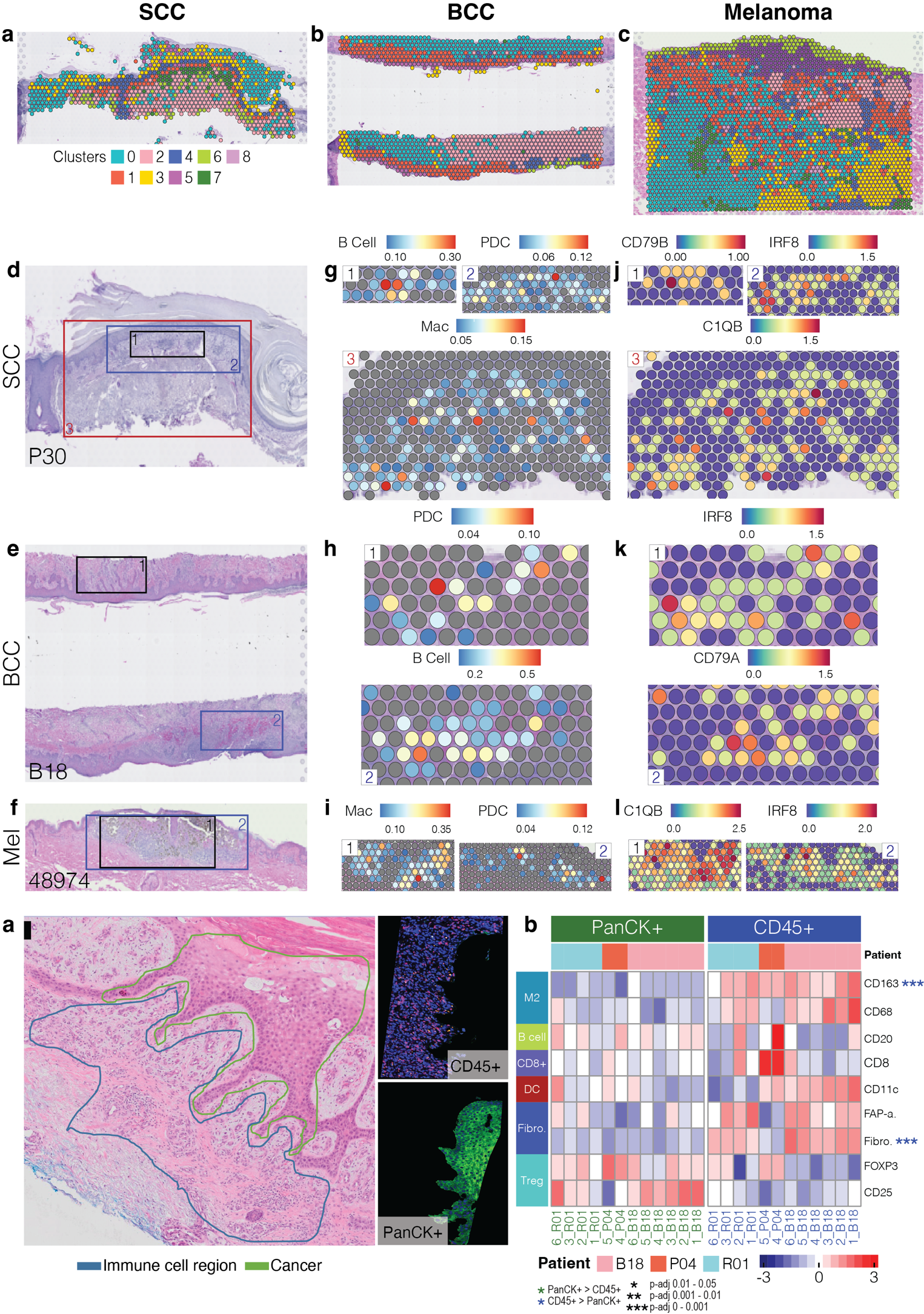

**Figure S10. Spatial mapping of cancer cell types within cSCC skin cancer as measured by GeoMx spatial transcriptomics.**

*(a) Left, H&E-stained image of cSCC tissue from patient B18, with pathologist’s annotation indicating cancer (green) and immune cell (blue) tissue regions. This annotation was used in combination with morphology marker staining to define regions of interest (ROIs) for GeoMx spatial proteomics. Right, selected ROIs corresponding to the highlighted regions (ROI 002, patient B18), segmented based on CD45 (top) and PanCK (bottom) markers.*

*(b) Validation of the presence of key cell types from our scRNASeq atlas, using protein markers captured by GeoMx. Selected markers are CD163 and CD68 (M2 markers), CD20 (B cell marker), CD8 (CD8+ T cell marker), CD11c (dendritic cell marker), FOXP3 and CD25 (Treg markers), FAP alpha and Fibronectin (fibroblast markers). Top annotation bars separate samples by segment (PanCK+ and CD45+ segments) and patient. Marker association with particular cell types was taken from NanoString product information. FAP-a. = FAP-alpha, Fibro. = Fibronectin. Asterisks indicate markers that were differentially expressed between CD45+ and PanCK+ segments; markers that were not found to be differentially expressed are included as their expression provides evidence for the presence of these cell types.*

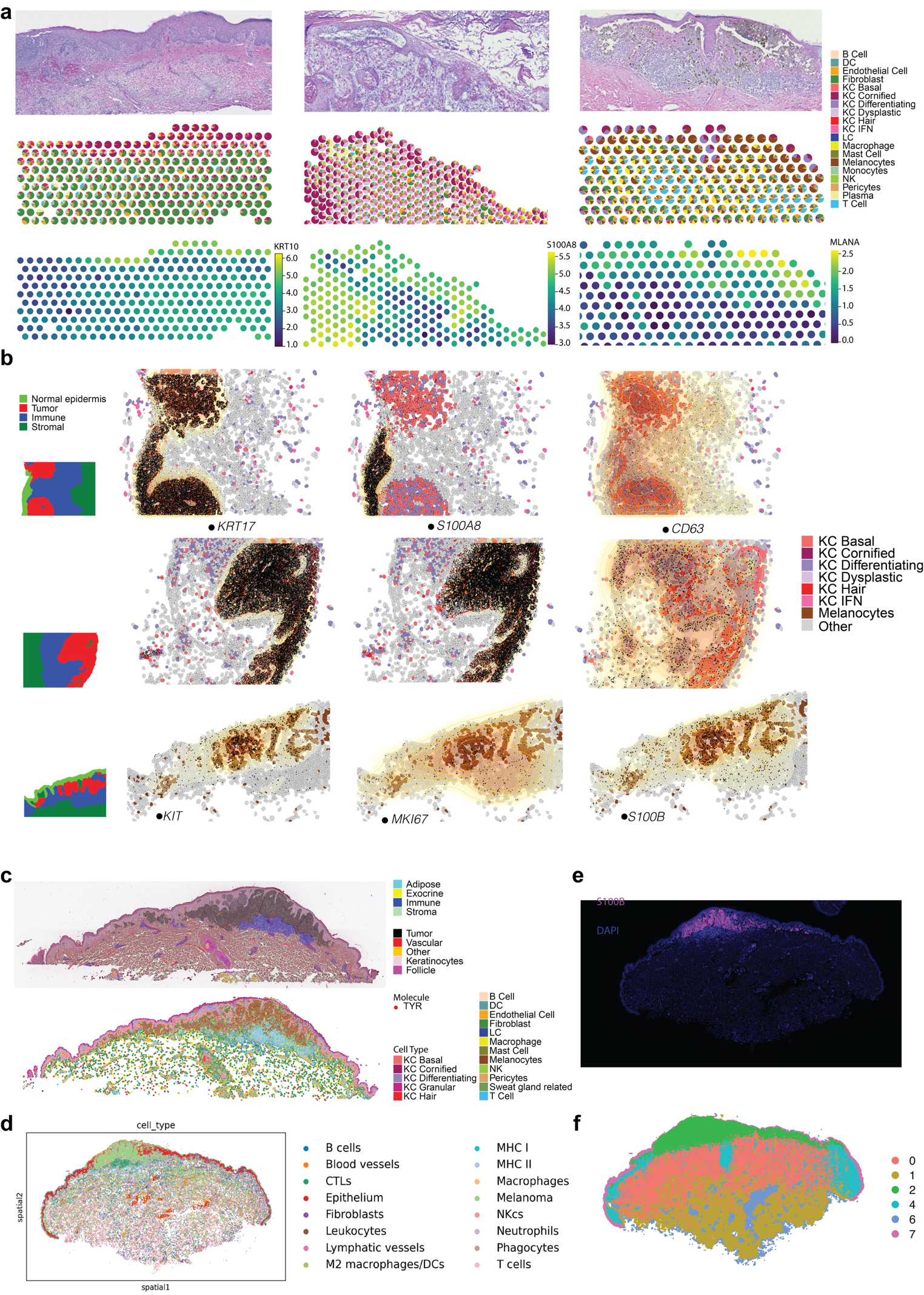

**Figure S11.** **Integrative analysis of spatial multiomics atlas for melanoma - Cross-modality spatial mapping across three cancer types.**

*(a) Visium mapping for BCC (left), cSCC (middle), and Melanoma (right), with Hematoxylin and Eosin (H&E) staining (top), cell type annotation (middle), and marker expression levels (bottom).*

*(b) Spatial localisation of marker gene expression in CosMx data from BCC (top), cSCC (middle), and melanoma (bottom). Cell type annotations and marker expression are shown together (bottom), with yellow regions highlighting contours of transcript density. The transcripts are represented as black dots. Pathological annotations are provided in the leftmost thumbnails.*

*(c) Pathological annotation of the Xenium melanoma sample (48974-2B) on H&E image (top), and corresponding cell type annotation and TYR marker gene expression shown for the same region (bottom).*

*(d) Cell type annotation for CODEX data on melanoma tissue*

*(e) Immunofluorescent staining of S100A8 protein marker corresponds to the tissue region containing melanoma cells in Panel e.*

*(f) Cell type annotation for spatial Glycomics data on an adjacent tissue section from the same block as shown in Panels d-e.*

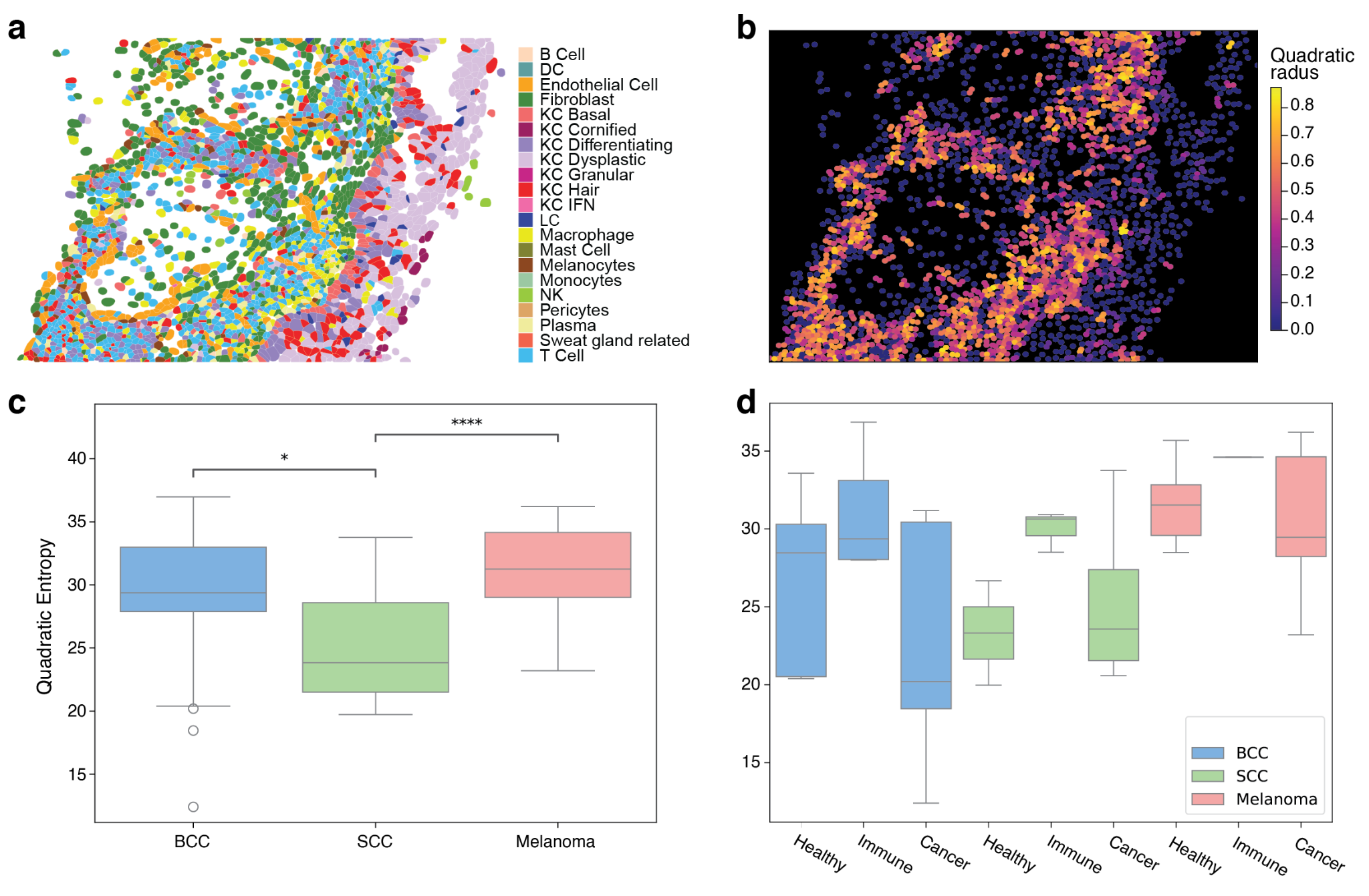

**Figure S12: Spatial cellular heterogeneity across cancer types using CosMx.**

*(a) Spatial cell type annotations of cSCC patient B18 FOV 15, the FOV with the highest overall heterogeneity score for this sample.*

*(b) Spatial cell heterogeneity plot for the FOV shown in Panel a. Cells are coloured by heterogeneity score (measured by cell neighbourhood analysis), highlighting regions of the FOV containing diverse cell types in close proximity.*

*(c) Boxplot of heterogeneity scores across three cancer types, indicating the lowest overall heterogeneity score for cSCC. Asterisks indicate significant p-values from paired Wilcoxon test.*

*(d) Boxplot of heterogeneity scores across tissue regions of each cancer type. Each FOV was designated as being either healthy, immune or cancer by a pathologist. Wilcoxon ranked sum test was applied.*

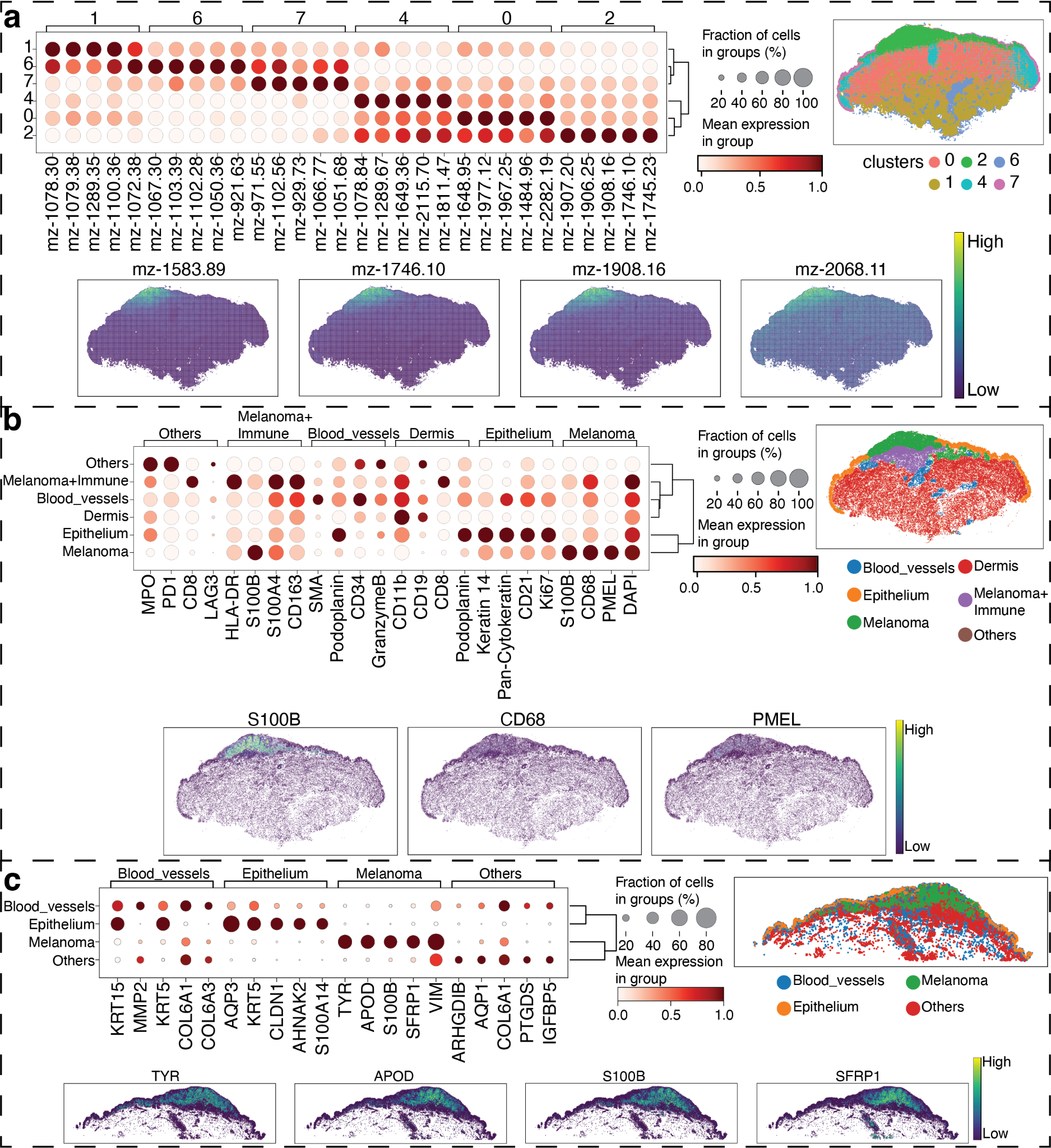
 **Figure S13. Integrative analysis of spatial transcriptomics, proteomics, and glycomics data for melanoma community.**

*(a) Joint pathway analysis using DE genes/proteins of melanocyte communities in Xenium and CODEX data, and glycans of melanocytes in MALDI data, to map to KEGG metabolic pathways. The X-axis shows enriched genes/proteins in Xenium and CODEX data, while the Y-axis shows enriched glycans.*

*(b-c) Cell type proportions of the community identified in CODEX data (b) and Xenium data (c). The melanoma community in both datasets is enriched with melanocytes.*

*(d) Clustering of the melanoma spatial glycomics sample, with the identification of the melanoma-enriched cluster 2. A heatmap of top markers for each cluster is on the left, and a tissue plot with cluster label is shown on the right. The expression for four representative metabolites is shown at the bottom on the panel.*

*(e) Annotation for CODEX data from the adjacent tissue section, showing the melanoma cluster, with a heatmap of protein markers for each cell type, the tissue plot showing all key cell types, and the the expression of the three proteins at the bottom.*

*(f) Similar to d) and e) the annotation of the melanoma adjacent tissue section for the Xenium data is shown.*

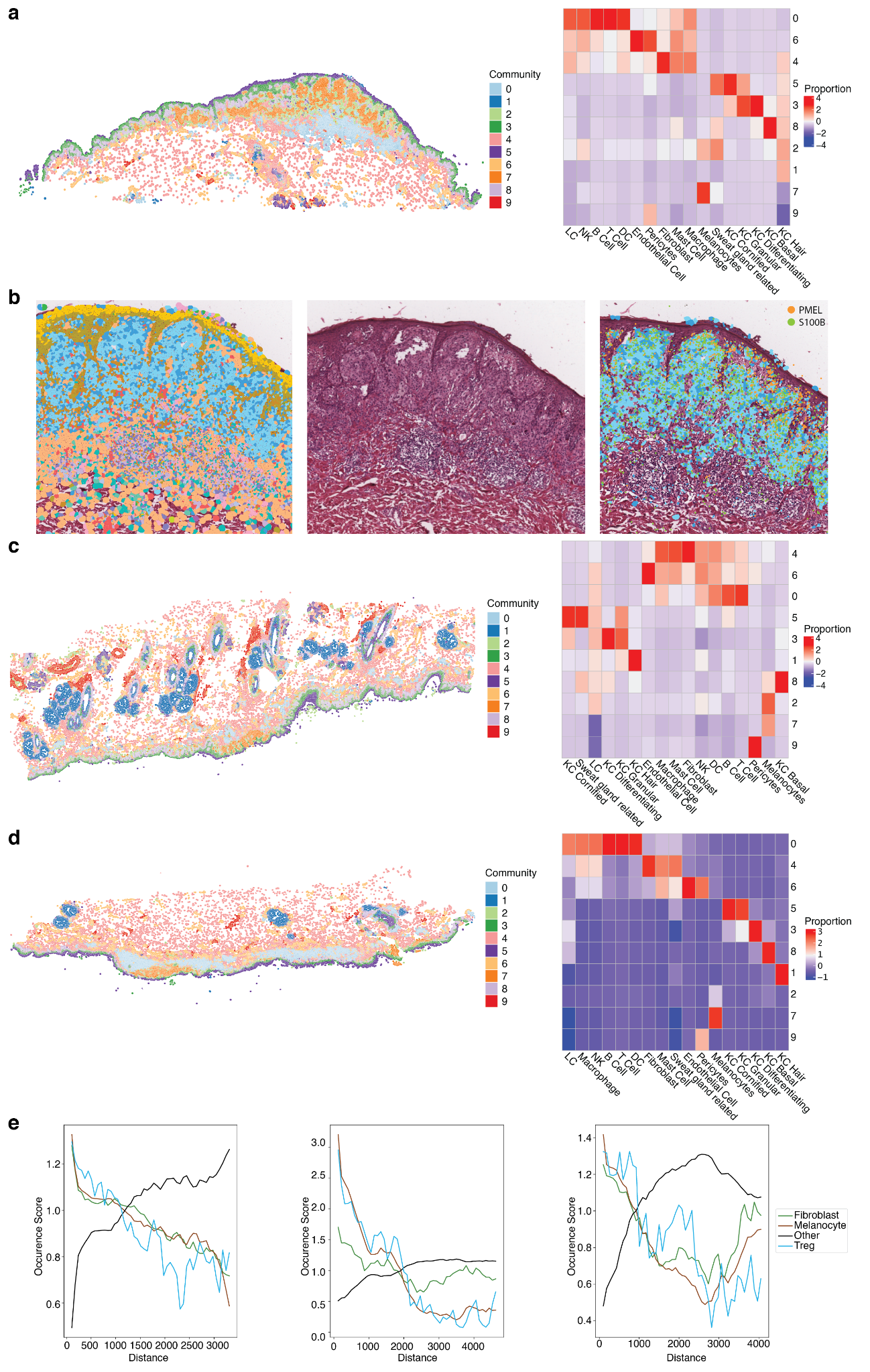

**Figure S14. A spatial community mapping of three melanoma samples for Xenium data.**

*(a) Community mapping for one sample based on cell type composition. The heatmap shows cell type composition per community.*

*(b) Expression of marker genes at single cell resolution, showing cell type (left), H&E image, and expression.*

*(c) Communities and cell type composition for two other melanoma samples.*

*(d) Cell-cell interaction analysis of the melanoma community. The line shows co-occurrence score with melanocytes, where a high score indicates the cells are closely distributed with melanocytes across the whole tissue. The scores change when the distance to search for neighbouring cells increases (i.e., less specific to a spatial domain).*

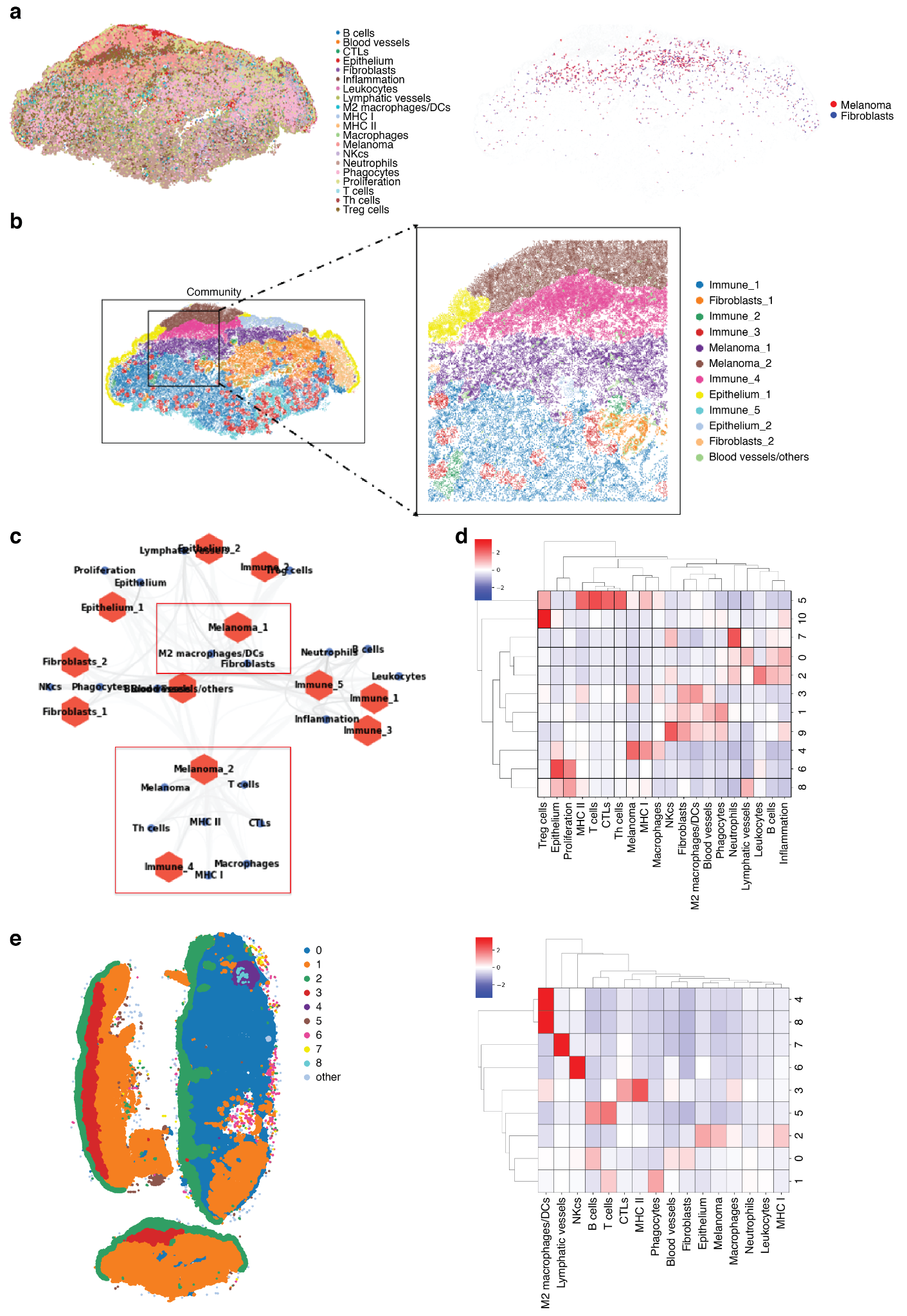

**Figure S15. A spatial community mapping of three melanoma samples for CODEX data.**

*(a) A detailed cell type annotation and an example of colocalization between two cell types (Melanoma and Fibroblasts) are shown.*

*(b) Community mapping with a zoom-in view of communities, highlighting two melanoma communities (shown. as melanoma-1 and melanoma-2).*

*(c) Community network analysis, with most similar communities connected in a nearest neighbourhood*

*(d) Examples of cell-type composition for each community of all three melanoma samples*

*(e) Spatial distribution of communities, defined from merged analysis of all three samples.*

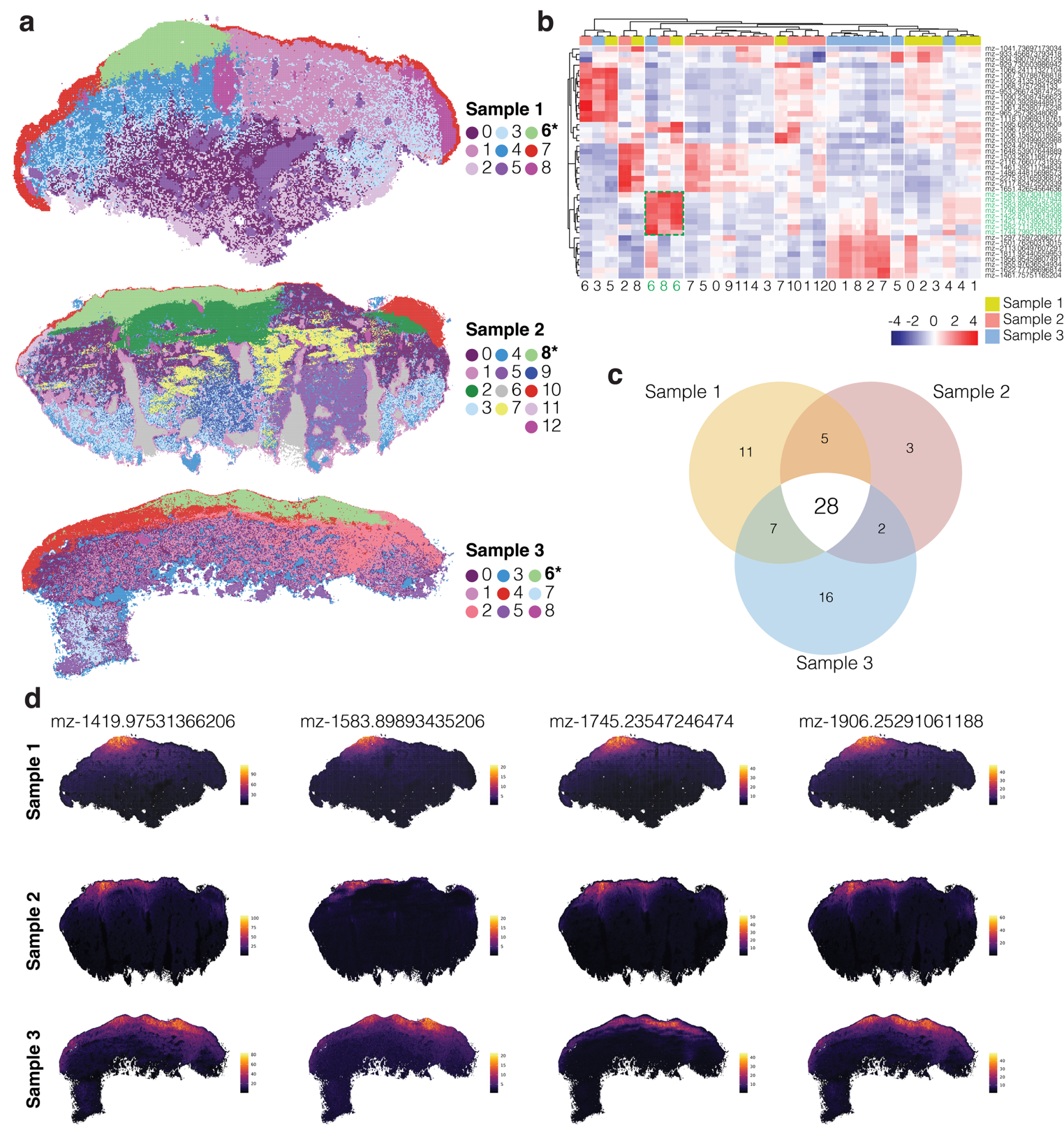

**Figure S16. Analysis of spatial glycomics data.**

*(a) Melanoma community analysis using spatial glycomics data. The colors show clustering results. Identification of a melanoma cluster (cluster 6 in samples 1 and 3, cluster 8 in sample 2; indicated with an asterisk) is based on histology and relative location compared to clusters from CODEX data.*

*(b) Heatmap showing the expression of top metabolite markers (named by the mass m/z values) for each cluster identified in A. Top annotation bar shows sample of origin. The green box and associated green labels highlight the metabolites enriched across the three melanoma clusters.*

*(c) Venn diagram highlighting the similarities and differences between markers identified for the melanoma cluster in each sample.*

*(d) Expression of the top four metabolite markers shared between samples. The top metabolites are the most differentially expressed compared to all other clusters.*

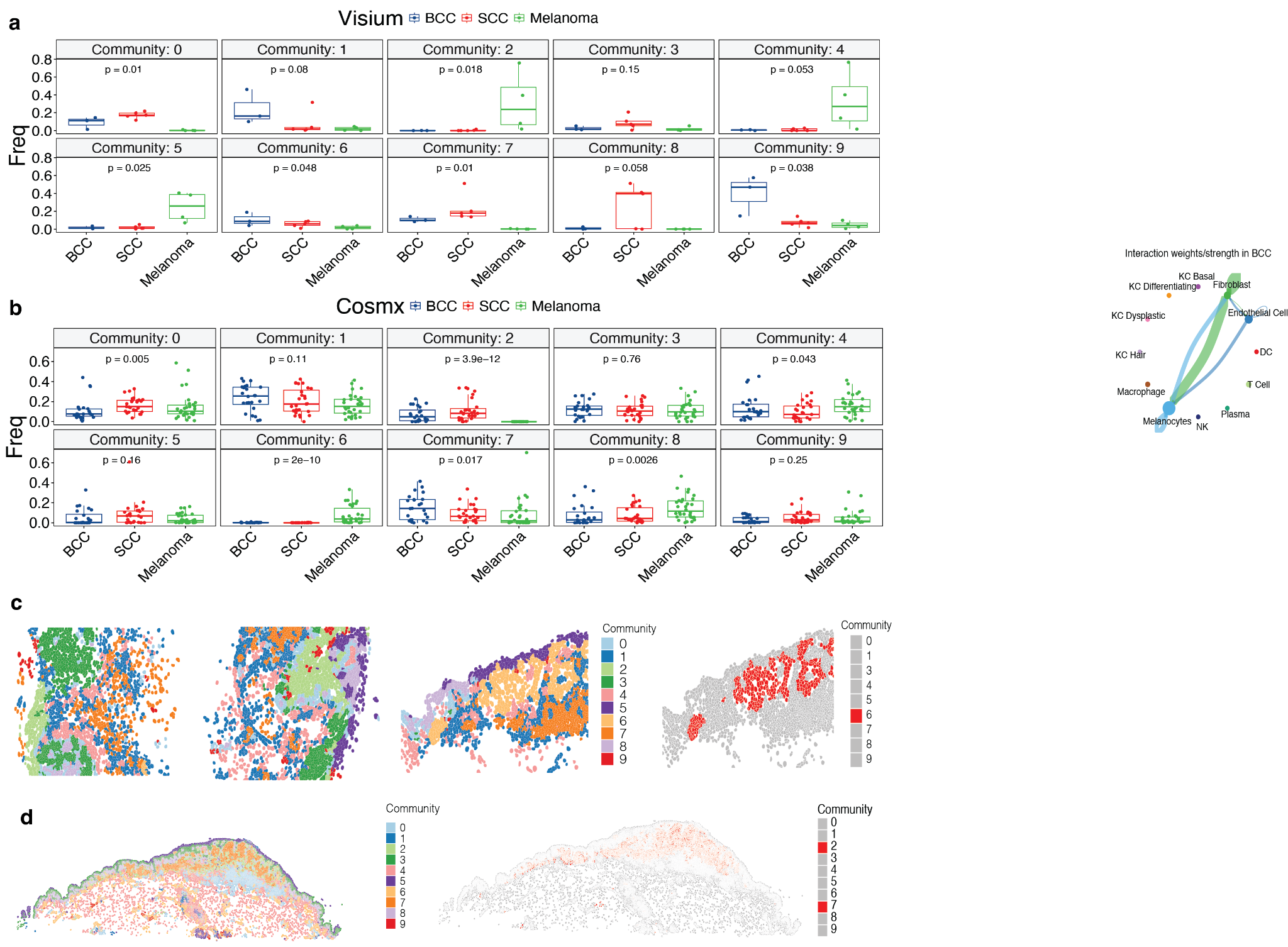

***Figure S17.*** **Integrative analysis across different individuals and technologies to build a robust spatial community atlas.**

*(a-b) Boxplots showing the proportion of each community deriving from BCC, cSCC and melanoma in Visium (a) and CosMx (b) data. Wilcoxon ranked sum test was applied.*

*(c) Spatial distribution of ten communities in CosMx data in representative FOVs from BCC (patient B18, FOV 13, left), cSCC (patient B18, FOV 14, second left) and melanoma (patient 48974-2B, FOV 12, second right). Pathological annotations for these FOVs are given in* ***Fig S11b****. Spatial location of cells from CosMX_6, a member of the melanoma meta-community (Fig 5), is shown next to the original distribution of all communities (right). The community 6 shown again here (also shown in Fig 5b) for visual comparisons.*

*(d) Spatial distribution of ten communities in Xenium melanoma sample (right) relative to the location of cells from Community Xenium_2 from the melanoma meta-community (left). The communities 2 and 7 shown again here (also shown in Fig 5b) for visual comparisons.*

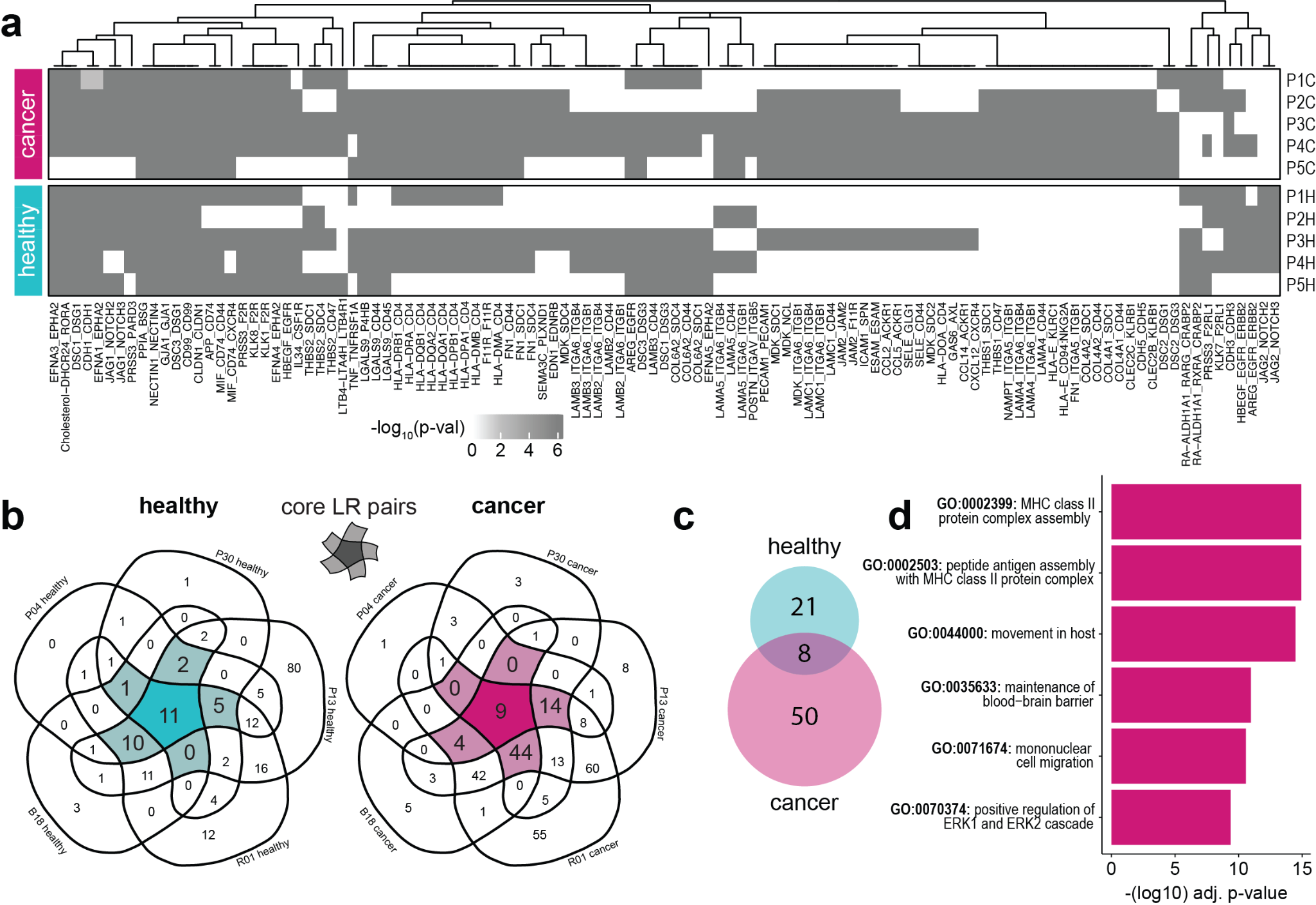

**Figure S18. Cell-cell interaction analysis of scRNAseq data for cSCC.**

*(a) Heatmap indicating significant LR pairs across samples. 312 significant pairs were detected from the entire suite of expressed genes in each sample. For visualisation purposes, only those LR pairs predicted in at least 4 of 10 samples are shown, with their corresponding -log10(pval) values. Non-significant LR pairs in each sample are white. LR communication was calculated between pairs of Level 2 cell types; duplicate LR pairs in each sample (i.e. predicted for multiple cell types within one sample) were collapsed, with only the highest -log10(pval) score shown.*

*(b) Patient-specific LR predictions in healthy (left) or cancer (right) cells across patients. The core suites of LR pairs that are shared across four or more of the five patients are highlighted in the centre of each Venn diagram.*

*(c) Overlap between core LR suites in cancer and healthy patients.*

*(d) The top six non-redundant GO terms that are enriched in the cancer-specific suite of LR genes, ranked by -log10 adjusted P-value. GO analysis was performed against a background gene universe of all LR pairs that are present in the CellChat database and are also expressed in our dataset.*

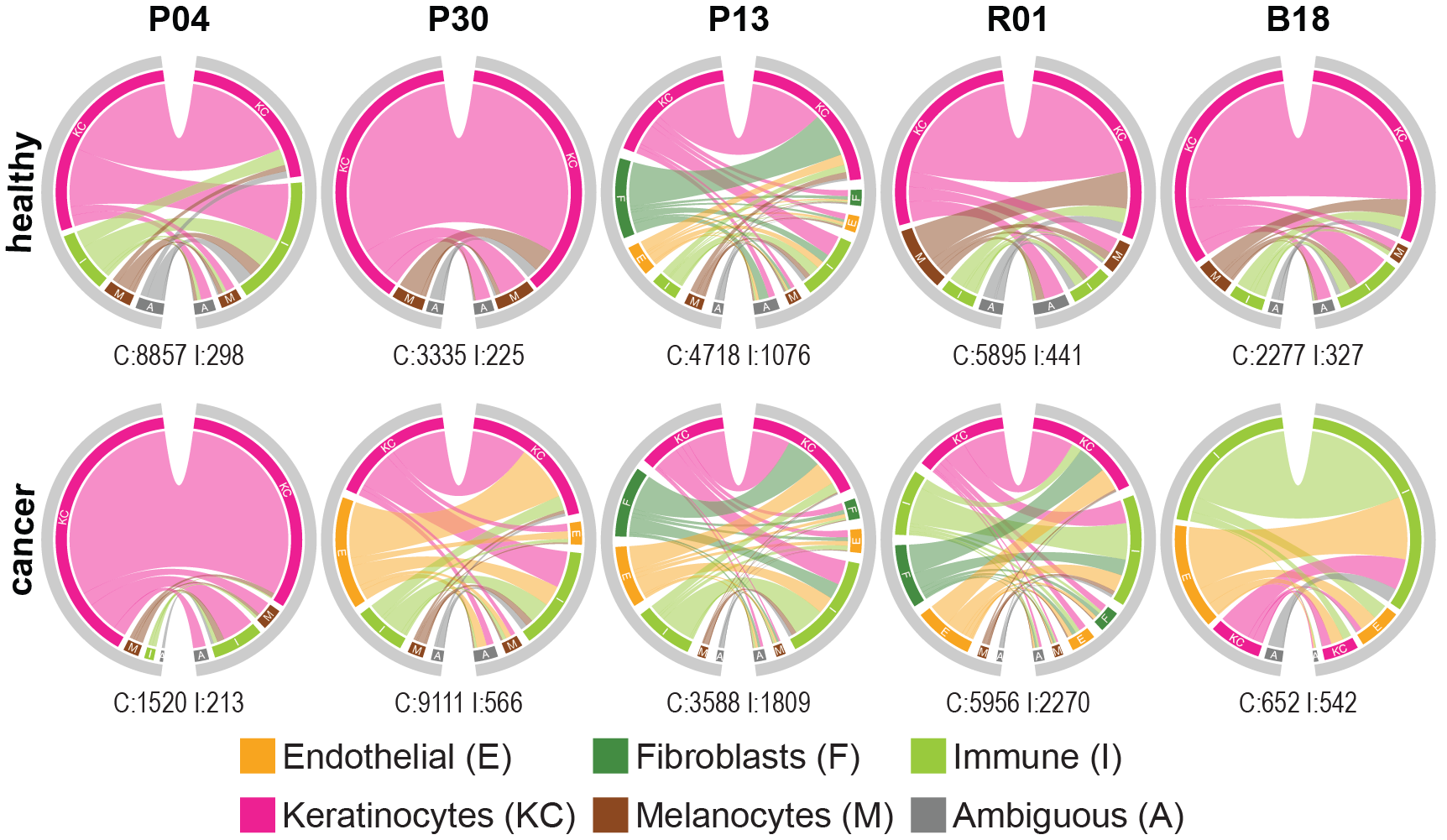

**Figure S19. CCI analysis in cSCC samples using scRNA-Seq.**

*Changes in cell type participation in cellular crosstalk between ligands and receptors in healthy and cancer samples from different patients. Circle diagrams show cells expressing ligands on the left, and receptors on the right. Connecting line colours indicate sender cell identity, and line widths indicate the number of unique LR connections predicted between pairs of cell types. Colours around the edge of the ring indicate sender and receiver cells’ identities. Numbers below each plot indicate the number of cells (C) and significant interactions (I) for each sample. Interactions were predicted for pairs of Level 2 cell types but summarised to broader cell type categories for visualisation.*

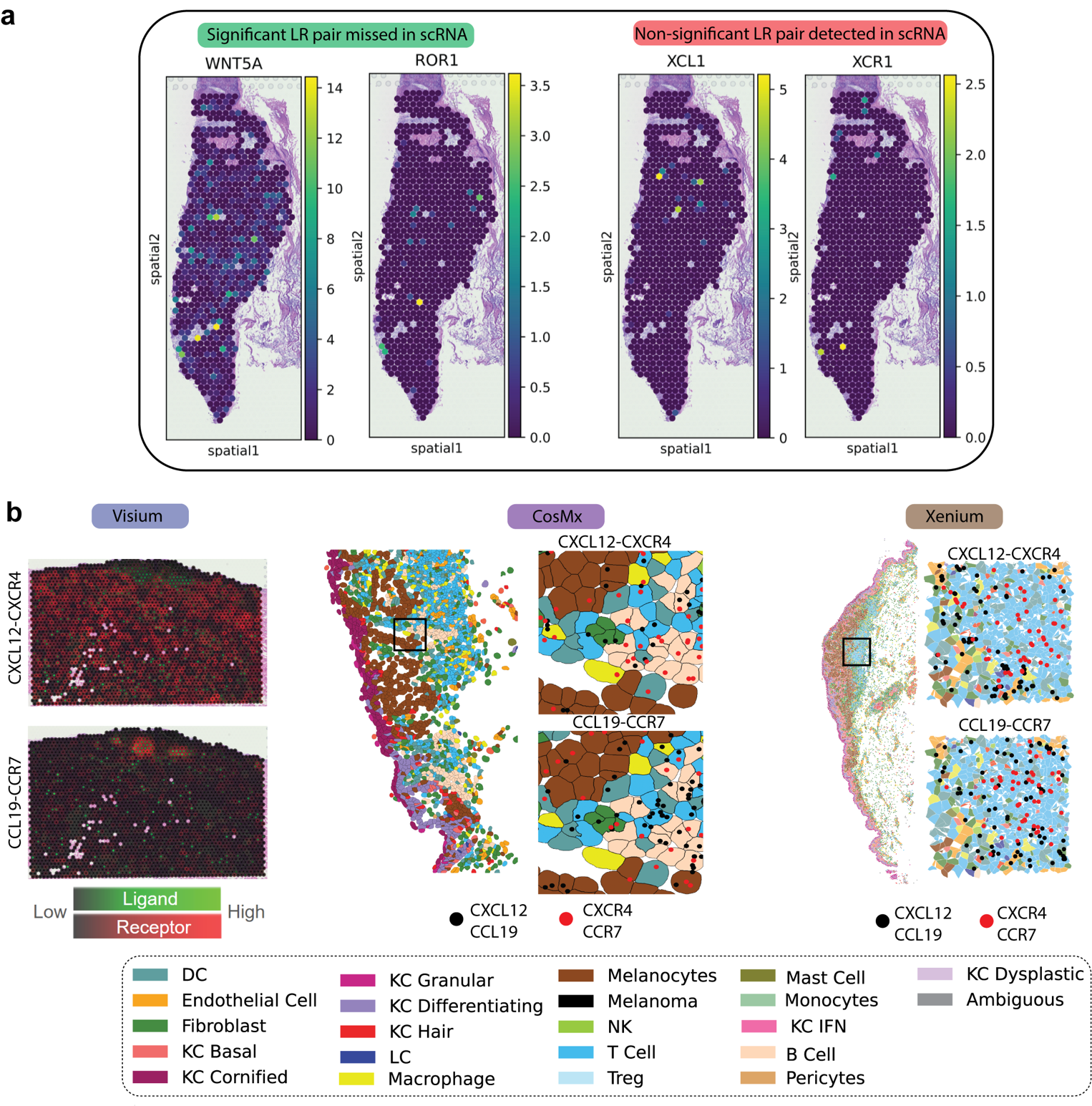

**Figure S20. Comparing LR detection between scRNAseq data without and different spatial transcriptomics platforms with spatial context.**

*(a) Examples of cases mis-detected by scRNAseq (WNT5A-ROR1) or detected by scRNAseq but not colocalize in spatial data, suggesting potential false positive.*

*(b) Consistent detection of LR pairs by Visium, CosMX and Xenium data*

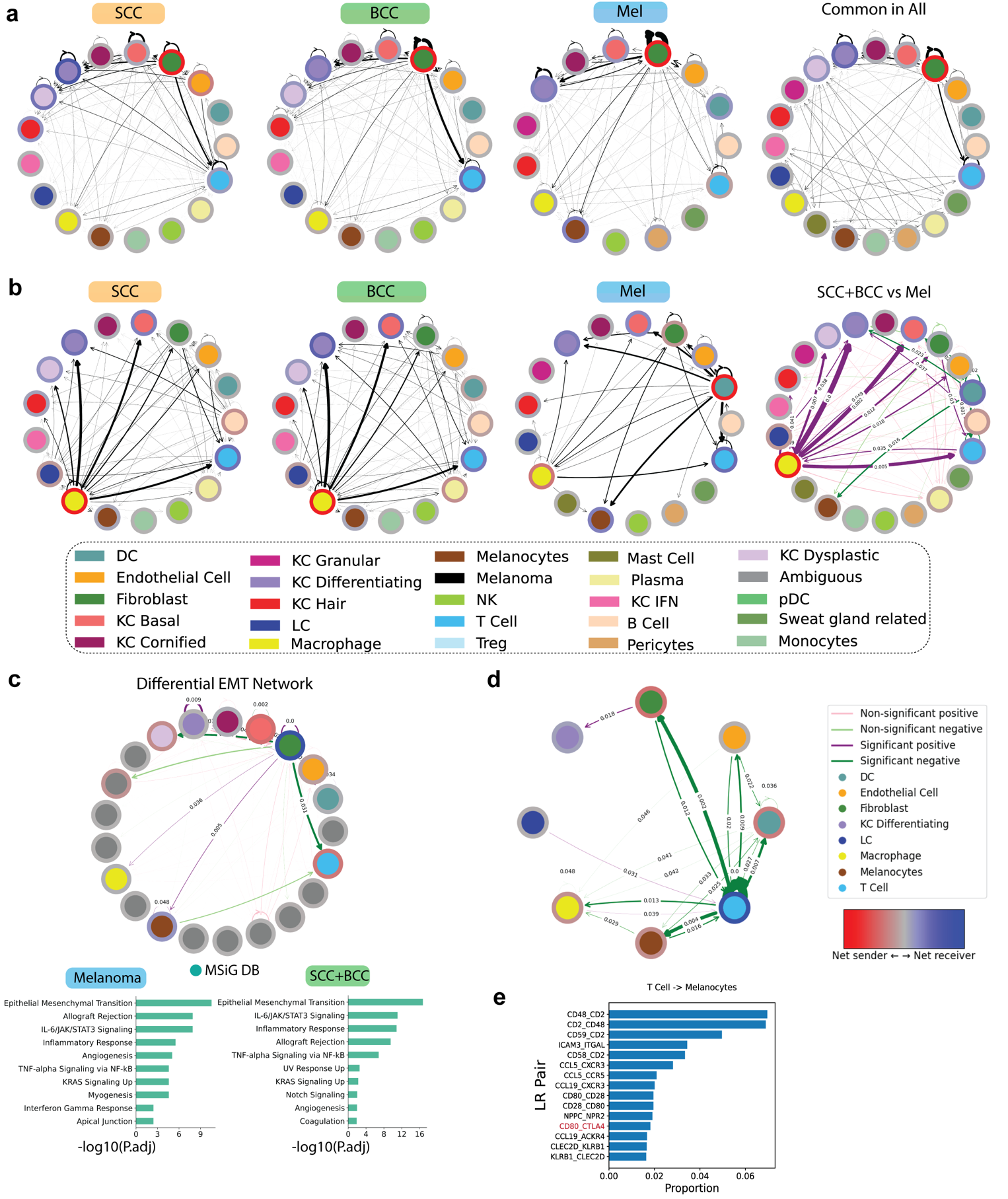

**Figure S21. Comparing interactions across the three cancer types.**

*(a) MMCCI integrated interactions for each cancer type and across multiple samples and spatial transcriptomics platforms. From left to right, circos interaction plots for BCC, cSCC and Melanoma. The last plot on the right is the integrated interactions for all cancer types for all samples.*

*(b) Comparing differential interactions at cell type level for BCC vs cSCC and for the integrated BCC-cSCC vs Melanoma using all L-R pairs. For cSCC vs BCC comparison, purple arrows show more interactions in BCC and green arrows indicating more in cSCC. For cSCC-BCC vs Melanoma, the purple arrows show more in melanoma and green arrows show more in cSCC-BCC.*

*(c) Differential interactions at cell type level using MIF-CD74 pair as the top different L-R pair shown in panel b. The difference indicates which cell type pairs are more actively interacting by considering the MIF-CD74 pair.*

*(d) Differential CCI network plots for LR pairs in the EMT pathway found from MSigDb pathway analysis. The pathway-centric interaction analysis shows specific cell type pairs where the EMT pathway was most significantly different. Pathway enrichment analysis from MSigDb on upregulated LR pairs common across CosMx and Visium 1) Melanoma and 2) cSCC+BCC.*

*(e) Interactions between cell-types in Melanoma compared to cSCC only.*

*(f) Top LR pairs interacting between T cells and melanocytes.*

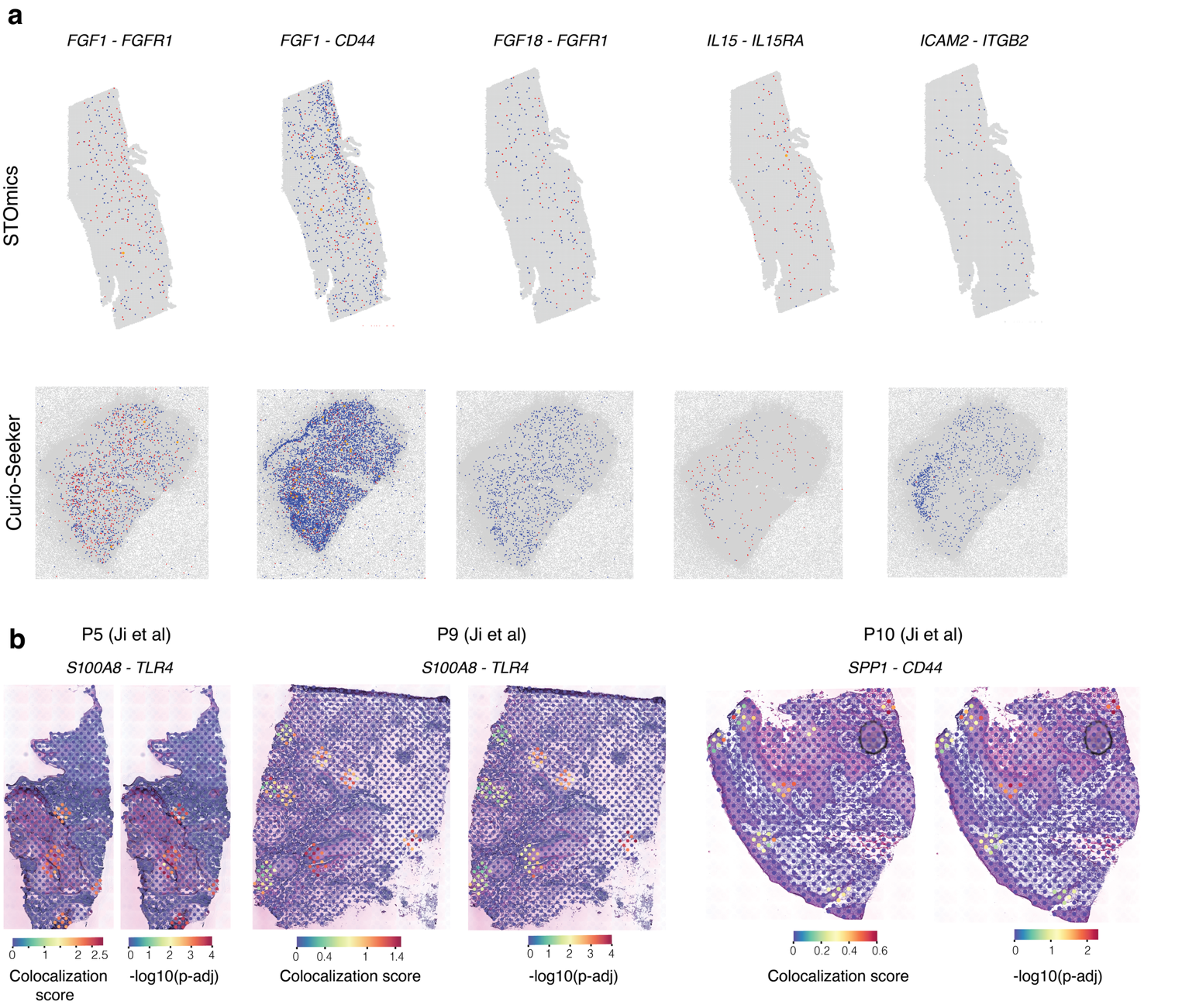

**Figure S22.** **Validation of cellular interactions using external datasets.**

*Colocalization of LR pairs replicated across public datasets with a significant p-value for:*

*(a) Melanoma samples from STOmics and Curio-Seeker. The red dots (cells) represent those expressing the ligand, blue cells represent those expressing receptor, and orange for those expressing both. All LR pairs with a significant colocalization are shown.*

*(b) SCC 10X Visium samples from the study by Ji et. al. The colocalization scores and p-values are shown at spot level for each tissue.*

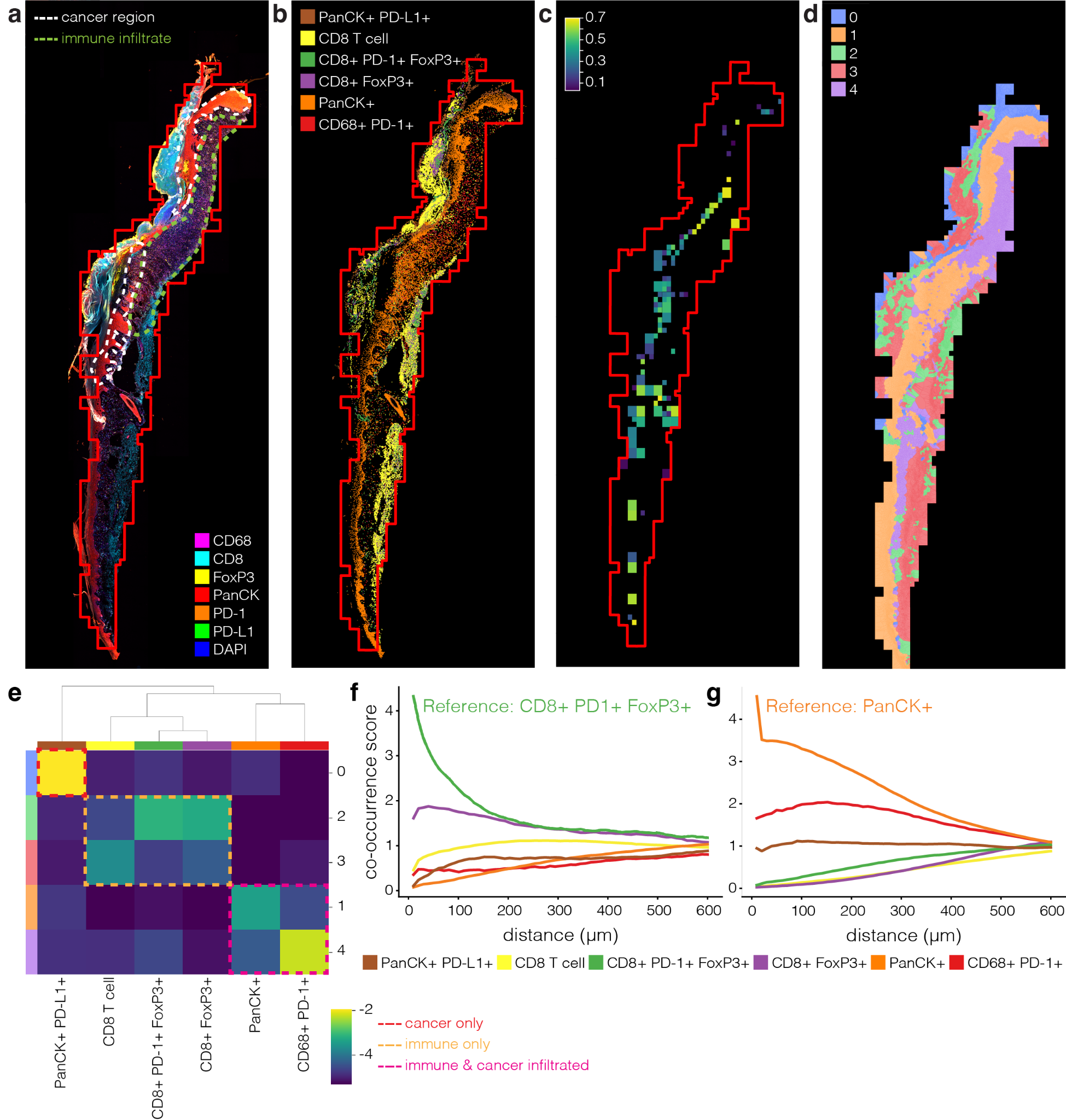

**Figure S23. Spatial cell type and community mapping at the protein level by targeted multispectral imaging (Polaris).**

*(a) Polaris protein imaging data from cSCC patient B18 with pathological annotation of cancer and immune regions, based on tissue morphology. A panel of six antibodies was used to profile protein markers and major cell types of interest including CD8 (T cell marker), FoxP3 (T-regulatory cell marker), CD68 (monocyte/macrophage marker), PanCK (epithelial cell marker), PD-1 and PD-L1 (immune checkpoint inhibitor, with the former expressed on immune cells and the latter on tumour cells); DAPI was also included as a nuclear stain. White and green dashed lines outline the cancer and immune regions of the tissue, as annotated by a pathologist. The red box indicates the tissue border as determined later during STRISH analysis, shown here to aid comparison with subsequent panels.*

*(b) Cell type classification based on clustering and gating of single-cell resolution protein expression signals.*

*(c) Mapping of cancer-immune cell co-localisation using STRISH, identifying image tiles containing double-positive CD8+ PD-1+ immune cells and PanCK+ PD-L1+ cancer cells. The identified colocalisation zone correlates with the cancer infiltrate region identified in Panel a.*

*(d) Voronoi diagram mapping the spatial distribution of cell communities, defined by further clustering of cell type classifications.*

*(e) Cell type composition of the five cell communities defined in Panel d, revealing higher-level groups of communities representing cancer only (red box), immune cells only (orange box), and cancer-immune infiltrate (pink box). Heatmap colours represent the neighbourhood enrichment score for each community-cell type combination.*

*(f-g) Cell type co-occurrence measurements of CD8+ PD-1+ FoxP3+ immune cells (f) and PanCK+ cancer cells (g) with either the same or other cell types in the dataset. Each line plots the co-occurrence score between the reference cell type and the test cell type calculated over increasing spatial distances.*

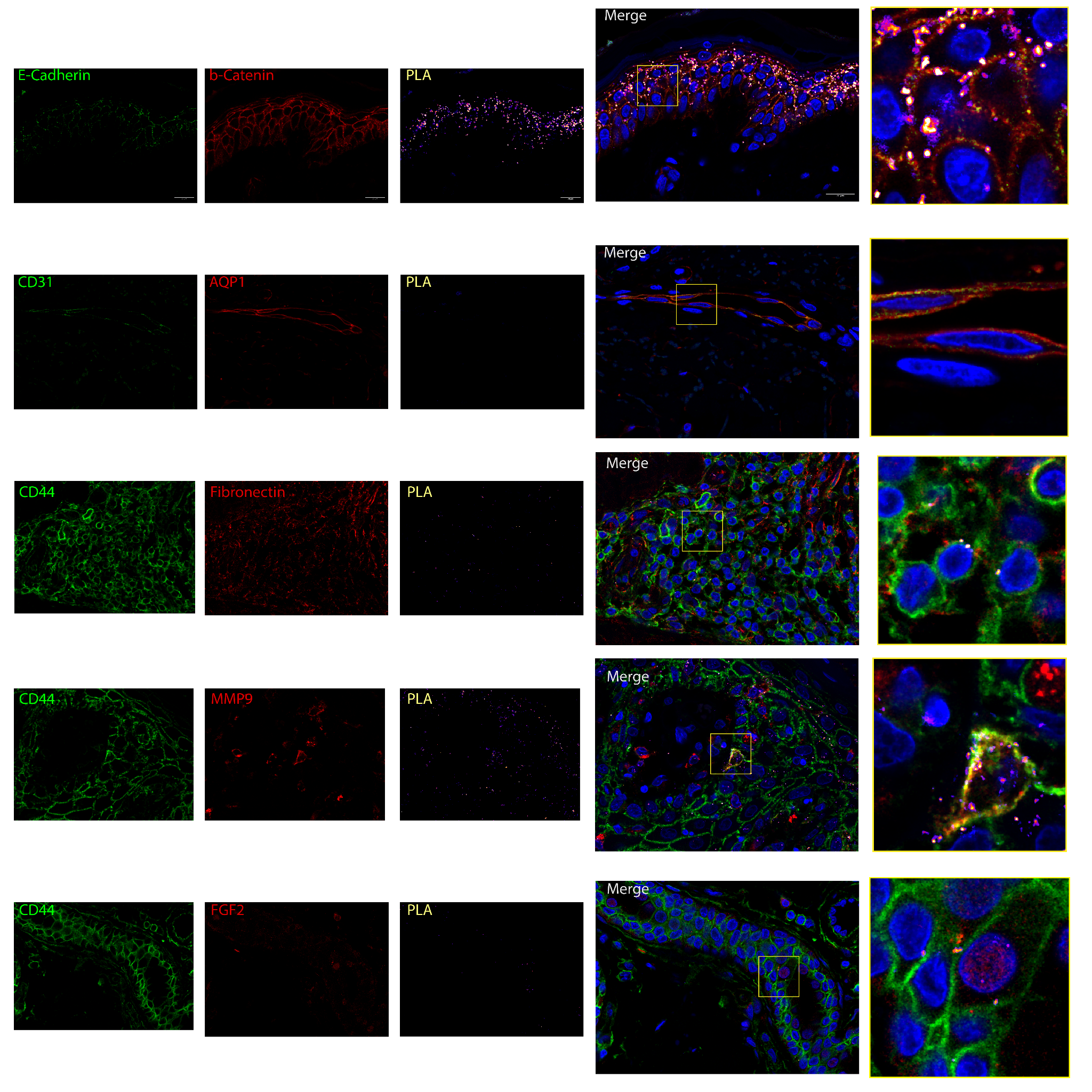

**Figure S24. Experimental validation of ligand-receptor interaction using proximal ligation assay for melanoma samples.** *In each row, fluorescence signal for a single antibody and PLA is shown, followed by a merge image and a Zoom-in window highlighting the interaction on the cell membrane. A positive PLA signal is visible within two proteins in a proximity less than 20 nm. From top row to bottom row, a positive control E-Cadherin and b-Catenin, a negative control for CD31 and AQP1, and the interaction results for the three pairs CD44-Fibronectin, CD44-MMP9, and CD44-FGF2 ligand-receptor pairs.*

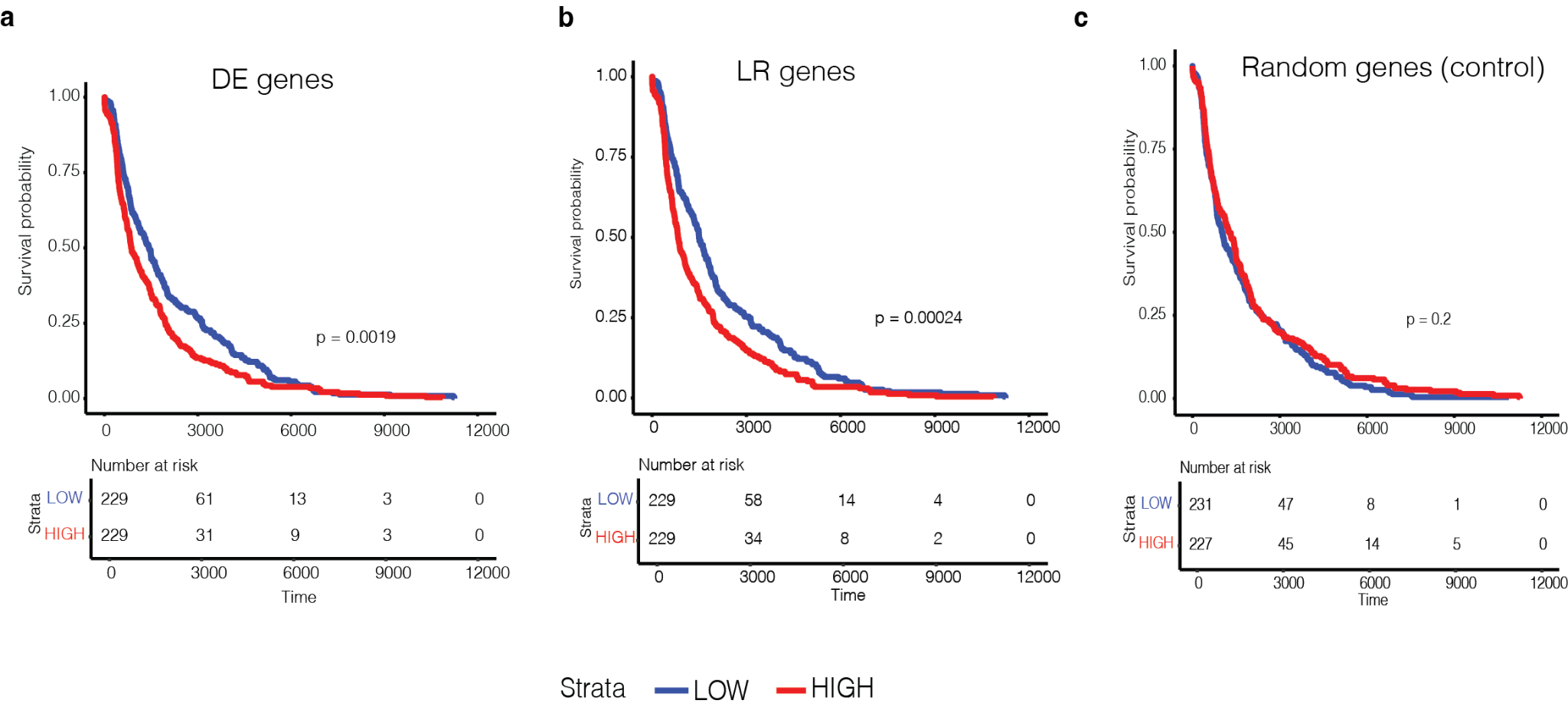

**Figure S25. Prognostic value of melanoma specific genes identified with scRNA-seq and spatial transcriptomics.**

*The plots indicate survival curves of patients from the TCGA SKCM dataset for groups stratified as high and low expressing patients for:*

*(a) genes upregulated in malignant melanocytes compared to normal melanocytes identified by DE analysis with edgeR*

*(b) LR genes upregulated in Melanoma compared to SCC and BCC*

*(c) a random set of genes excluding the genes used in a and b. The melanoma specific genes identified in our study are associated with poor prognosis where the high expressing patients indicate significantly poor survival while the random genes show no significant stratification.*

### **List of Supplementary Tables**

**Table S1:** Summary of samples used in this study

**Table S2: Table of genes upregulated in Cancer KCs vs Other KCs in the scRNA-seq dataset in the pseudobulk replicate 3. 'Heatmap_top50_common' represents the top 50 genes upregulated in all three pseudobulk replicates in the DE analysis with edgeR**

**Table S3: Table of genes upregulated in Melanoma vs Melanocytes in the scRNA-seq dataset in the pseudobulk replicate 3. 'Heatmap_top50_common' represents the top 50 genes upregulated in all three pseudobulk replicates in the DE analysis with edgeR**

**Table S4: Table of genes upregulated in Melanocytes from Melanoma patients sequenced on CosMx platform compared to the melanocytes in normal Xenium samples**

**Table S5: Table of differentially expressed L-R pairs comparing each cancer type vs the others using pseudobulked L-R scores with 3 pools per sample.**

**Table S6:** Pre-oligonucleotide-conjugated-antibodies and complementary reporters

**Table S7:** Table of Primary antibodies used in PLA assay

**Table S1. 24 patient samples to generate the multiomics data resource, suitable for addressing multiple questions.**

| **Patient#** | **Cancer type/s** | **Age** | **Sex** | **Lesions** | **scRNAseq (cSCC) snRNAseq (Mel)** | | **Visium** | | | **CosMx** | | **Polaris** | | **RNAScope** | | | **GeoMx (RNA)** | | **GeoMx (protein)** | | **Codex** | | | **Xenium** | |
| --- | --- | --- | --- | --- | --- | --- | --- | --- | --- | --- | --- | --- | --- | --- | --- | --- | --- | --- | --- | --- | --- | --- | --- | --- | --- |
|  |  |  |  |  | **cSCC** | **Mel** | | **cSCC** | **BCC** | | **Mel** | **cSCC** | **BCC** | | **Mel** | **cSCC** | | **cSCC** | **BCC** | **cSCC** | | **cSCC** | **Mel** | | **Mel** |
| **B18** | cSCC + BCC^†^ | 76 | F | 1x leg BCC  1x face IEC  1x leg IEC | ✔️*^†^ |  | | ✔️ | ✔️^ | |  | ✔️ | ✔️ | |  | ✔️ | | ✔️ | ✔️^§^ | ✔️ | | ✔️ |  | |  |
| **E15** | cSCC + BCC | 54 | F | 1x leg BCC  2x leg IEC  1x leg cSCC |  |  | | ✔️ | ✔️ | |  |  |  | |  |  | | ✔️^§^ | ✔️^§^ |  | |  |  | |  |
| **F21** | cSCC + BCC | 76 | M | 1x face BCC  1x leg cSCC |  |  | | ✔️ | ✔️ | |  |  |  | |  |  | | ✔️^§^ | ✔️^§^ |  | |  |  | |  |
| **P30** | cSCC | 65 | M | 2x arm IEC | ✔️* |  | | ✔️ |  | |  | ✔️ |  | |  |  | |  |  |  | |  |  | |  |
| **P13** | cSCC | 75 | M | Leg | ✔️* |  | | ✔️ |  | |  | ✔️ |  | |  |  | |  |  |  | |  |  | |  |
| **P04** | cSCC | 72 | M | Arm | ✔️* |  | |  |  | |  |  |  | |  | ✔️ | |  |  | ✔️ | | ✔️ |  | |  |
| **R01** | cSCC | 73 | M | Neck | ✔️* |  | |  |  | |  |  |  | |  | ✔️ | |  |  | ✔️ | | ✔️ |  | |  |
| **D12** | BCC | 82 | M | Face |  |  | |  |  | |  |  | ✔️ | |  |  | |  |  |  | |  |  | |  |
| **6747-085P** | Mel | 89 | M | Chest |  |  | |  |  | | ✔️ |  |  | | ✔️ |  | |  |  |  | |  |  | |  |
| **21031-08TB** | Mel | 89 | M | Face |  |  | |  |  | | ✔️ |  |  | | ✔️ |  | |  |  |  | |  |  | |  |
| **48974-2B** | Mel | 79 | F | Back |  |  | |  |  | | ✔️ |  |  | | ✔️ |  | |  |  |  | |  |  | |  |
| **66487-1A** | Mel | 53 | F | Arm |  |  | |  |  | | ✔️ |  |  | | ✔️ |  | |  |  |  | |  |  | |  |
| **6475-07FC** | Mel | 53 | F | Abdomen |  |  | |  |  | |  |  |  | |  |  | |  |  |  | |  | ✔️ | | ✔️ |
| **9474-06BR** | Mel | 55 | M | Back |  |  | |  |  | |  |  |  | |  |  | |  |  |  | |  | ✔️ | | ✔️ |
| **23346-10SP** | Mel | 70 | M | Back |  |  | |  |  | |  |  |  | |  |  | |  |  |  | |  | ✔️ | | ✔️ |
| **53023-07BR** | Mel | 81 | M | Face |  |  | |  |  | |  |  |  | |  |  | |  |  |  | |  |  | | ✔️ |
| **98594-09PY** | Mel | 65 | F | Back |  |  | |  |  | |  |  |  | |  |  | |  |  |  | |  |  | | ✔️ |
| **30037-07BR** | Mel | 76 | M | Scalp |  |  | |  |  | |  |  |  | |  |  | |  |  |  | |  |  | | ✔️ |
| **healthy skin 1** | Healthy ctrl | 27 | M | Forearm |  |  | |  | ✔️ | |  |  |  | |  |  | |  |  |  | |  |  | | ✔️ |
| **healthy skin 2** | Healthy ctrl | 41 | M | Forearm |  |  | |  | ✔️ | |  |  |  | |  |  | |  |  |  | |  |  | | ✔️ |
| **healthy skin 3** | Healthy ctrl | 45 | M | Forearm |  |  | |  | ✔️ | |  |  |  | |  |  | |  |  |  | |  |  | | ✔️ |
| **MPS13** | Mel  Malig. | 28 | F | Right upper arm |  | ✔️ | |  |  | |  |  |  | |  |  | |  |  |  | |  |  | |  |
| **MPS42** | Mel  intermediate | 35 | M | Back |  | ✔️ | |  |  | |  |  |  | |  |  | |  |  |  | |  |  | |  |
| **MPS43** | Mel  benign | 37 | F | Left upper back |  | ✔️ | |  |  | |  |  |  | |  |  | |  |  |  | |  |  | |  |
|  | # B18 = P5, P30 = P2, P13 = P3, P04 = P1, R01 = P4 in Supplementary Tables  * Healthy and cancer biopsies  ^†^ Cancer biopsy from patient B18 contained pooled cSCC and BCC cells  ^ 3x biological replicates  ^§^ Tran et. al. 2022 | | | | | | | | | | | | | | | | | | | | | | | | |

### **Supplementary Methods**

**Integrating 12 Spatial and Single Cell Technologies to Characterise Tumour Neighbourhoods and Cellular Interactions in three Skin Cancer Types**

P. Prakrithi^1,2#^, Laura F. Grice^1,2,3#^, Feng Zhang^1,2,4#^, Levi Hockey^1,2^, Samuel X. Tan^5^, Xiao Tan^1,2^, Zherui Xiong^1,2^, Onkar Mulay^1,2^, Andrew Causer^1,2^, Andrew Newman^1,2^, Duy Pham^1^, Guiyan Ni^1^, Kelvin Tuong^6^, Xinnan Jin^1,2^, Eunju Kim^2,^, Minh Tran^1^, Hani Vu^1,2^, Nicholas M. Muller^5^, Emily E. Killingbeck^7^, Mark T. Gregory^7^, Siok Min Teoh^1^,Tuan Vo^1^, Min Zhang^8^, Maria Teresa Landi^9^, Kevin M. Brown^9^, Mark M. Iles^10^, Zachary Reitz^7^, Katharina Devitt^5^, Liuliu Pan^7^, Arutha Kulasinghe^5^, Yung-Ching Kao^5^, Michael Leon^7^, Sarah R. Murphy^7^, Hiromi Sato^7^, Jazmina Gonzalez Cruz^5^, Snehlata Kumari^5^, Hung N. Luu^11^, Sarah E. Warren^7^, Chris McMillan^12,13^, Joakim Henricson^14,15^, Chris Anderson^14,15^, David Muller^12,13^, Arun Everest-Dass^16^, Blake O’Brien^17^, Huanwei Wang^18^, Mathias Seviiri^18^, Matthew H. Law^18, 19, 20^, H. Peter Soyer^5^, Ian Frazer^5^, Youngmi Kim^7,21^, Mitchell S. Stark^5^, Kiarash Khosrotehrani^5^, Quan Nguyen^1,2^

**1. Single-cell RNA sequencing, data pre-processing and annotation**

***Sample preparation***

10x Genomics Chromium single-cell library preparation and sequencing was performed according to the manufacturer’s instructions, using the Single Cell 3' Library, Gel Bead and Multiplex Kit (version 2, PN-120233; 10x Genomics). Cell numbers were optimised to capture approximately 3,000 cells per reaction. Single-cell transcriptome libraries were sequenced on an Illumina NextSeq500 machine, using a 150-cycle High Output reagent kit (NextSeq500/550 version 2, FC-404-2002; Illumina) with the following reads and indices: read 2 (98 bp), i7 index (8 bp), read 1 (cell barcodes and UMI; 26 bp).

**scRNAseq** ***data processing (cSCC-BCC)***

The BCL sequencing output file was converted to a FASTQ file using bcl2fastq/2.17. Read mapping was performed using CellRanger/3.0.2 against the *Homo sapiens* GRCh38p10 reference genome. Median absolute deviation (MAD) filtering was performed using scater v1.14.6 to remove outlier cells with low library size and/or gene counts (>3 MADs below the median value), or high mitochondrial or ribosomal gene percentage (>3 MADs above the median value). Doublet scoring was performed using scds v1.2.0 and cells were removed if they were predicted as doublets by at least two of the three included prediction methods (bcds, cxds, or hybrid), and expressed more than 3,000 genes. After QC filtering, the 48,226 remaining cells in P30 were randomly subsampled to 3,335 cells due to its much larger depth compared to other samples. This number of cells was selected as the other four healthy samples contained an average of 3,335.25 cells each. Samples were processed individually using Seurat version 5.0 before being integrated using Seurat canonical correlation analysis (CCA). Following integration, dimensionality reduction (using the first 20 principal components), clustering and sub-clustering, and marker prediction were all performed in Seurat v5.0. Different cluster resolutions were tested and assessed for cluster stability using Clustree v0.4.2.

***Cell* type *annotation BCC-cSCC***

We performed cell type annotation at three granular levels, which we call “Level 1” , “Level 2” and “Level 3” annotation. For the broader Level 1 annotation, the integrated set of 11 samples was clustered using Seurat (resolution 0.4, 20 principal components (PCs)) ^1^. Seurat’s FindAllMarkers function was used to find the top differentially expressed genes for each cluster. Cluster annotation was performed based on manual comparison of the top gene markers with the literature. To determine more detailed annotations, each major cell type (endothelial cells, fibroblasts, immune cells, keratinocytes and melanocytes) was extracted and re-clustered in turn for our Level 2 classification. Optimal cluster resolutions were selected using clustree. Cell subtypes were identified by comparing marker genes, detected using Seurat’s FindAllMarkers function, with the literature.

**Annotation of aneuploid cells using inferred CNV profiles**

We applied two top-performing tools, CopyKAT v1.1.0 and InferCNV v1.19.1, to add complementary information to prioritise malignant cells. In both cases, the raw count matrices were used as the input, using the default parameters and the reference free option (no normal cell IDs were provided). CopyKAT predicted ploidy status for each cell barcode, including Aneuploid, Diploid and Not Defined (for uncertain cells, often associated with low quality data). To define a set of ‘confident aneuploid’ cells, we integrated results from CopyKat and InferCNV. Although InferCNV does not predict aneuploid cells and is mainly used to infer CNV profiles to further stratify the known malignant cells into subclones, we made use of its quality filtering analysis results. InferCNV excludes cells of low quality and the ones with CNV profiles that are likely to be normal and retains only the cells likely to be aneuploid. Finally, we called the ‘Aneuploid’ cells that passed the selection criteria of both tools. The inferred CNV results were then combined with DE gene module score analysis to finally classify keratinocytes or melanocytes as cancer cells.

**Single Nucleus RNA Sequencing (snRNAseq) of melanoma samples**

snRNAseq was performed using nuclei dissociated from two 25-μm FFPE melanoma tissue sections. Nuclei dissociation was carried out using the gentleMACS Octo Dissociator (Miltenyi Biotec) with the default program 37C_FFPE_1. The dissociated nuclei were purified by flow cytometry based on Ethidium Homodimer-1 staining to exclude keratin debris. Library preparation was performed following the Chromium Fixed RNA Profiling user guide (10x Genomics). Single-nucleus emulsions (GEMs) were generated using the Chromium X system (10x Genomics). Libraries were constructed from the captured nuclei by probe ligation, probe extension, amplification, and indexing. Library was sequenced on a NextSeq 2000 platform (Illumina) with a dual-indexed sequencing run following read configuration: Read1, 28 cycles; i7 index, 10 cycles; i5 index, 10 cycles; and Read2, 90 cycles.

**snRNAseq data processing (melanoma samples)**

The reads were mapped to the reference Human Transcriptome Probeset V1.0.1 GRCh38 (2020-A) using CellRanger (V7.0), implementing the cellranger multi pipleine. The filtered cell count matrix from CellRanger output was used for downstream pre-processing, integration and clustering analysis. Doublets and cells with more than 7000 genes or over 20,000 reads were removed. Harmony was used to integrate the datasets from the three patients (n_neighbors**=**30, n_pcs**=**75). Leiden clustering was then applied with the default resolution 1, resulting in 33 clusters. Similar to the annotation pipeline applied for the cSCC-BCC data, the melanoma clusters were annotated at three levels, based on gene markers most differentially expressed from the Wilcoxon test comparing each cluster to all the other clusters. A list of known skin gene markers were also used to check for specificity of marker genes to corresponding clusters. All melanocytes defined from the level 1 clustering were further subclustered, forming seven melanocyte subclusters. The immune cells were also subclustered into eight level 2 clusters, and a total of 19 clusters at the level 3 clustering.

To define melanoma, only melanocytes from the patient with definite invasive melanoma (MPS13) were used to find highly-likely cancer cells. Two selection criteria include that the melanocytes had to have a high “melanoma score” and that the cell had an aneuploid genotype as predicted by inferCNV and CopyCAT. To compute a module score we selected a list of genes upregulated in the melanoma sample compared to the benign sample using both edgeR pseudobulking and *scanpy* non-parametric test. A density plot was produced to inform the cutoff selection. As the majority of the cells were with a score >80th percentile cut-off. To increase confri inferred ‘Aneuploid’ by the CNV analysis and with a high module score by both the aforementioned methods are labelled as malignant melanocytes (red) as shown in the UMAP.

***scRNAseq KC cancer vs TSK cell comparison***

cSCC scRNA-seq data from Ji et al. (2020) were obtained from the GEO database (accession GSE144236) and subset to cells annotated as epithelial cells (Level1_celltype). Data were re-processed in Seurat following the authors’ tutorials. UMAPs generated from either merged data or Seurat-integrated data showed better separation of published cell type annotations in the merged object (data not shown), so this object was used for downstream analyses. The top 100 TSK marker genes were taken from Table S3 of Ji et al. (2020). For our dataset, top 100 markers per keratinocyte cell type were identified using Seurat’s **FindAllMarkers** function (parameters: only.pos = TRUE, min.pct = 0.25, logfc.threshold = 0.25, adjusted p < 0.05). Gene lists from both studies were used to calculate module scores with Seurat’s **AddModuleScore** function; top 100 markers were used for boxplots, and the top 20 for dot plots and UMAP visualisation. Overlap between the top 100 markers of each list was assessed by hypergeometric test, using the set of genes expressed in both datasets as the background gene set.

***scRNASeq* cancer *vs healthy comparison***

Global transcriptional differences between healthy and cancer cells, either across the whole dataset or for each patient individually, were determined using Seurat’s FindAllMarkers function (with parameters only.pos = TRUE, min.pct = 0.25, logfc.threshold = 0.25 and with adjusted p-value threshold ≤ 0.05). Comparisons between patients were performed using the Venn diagram tool (<http://bioinformatics.psb.ugent.be/webtools/Venn/>). Differentially expressed genes detected in ≥4 patients for either cancer or healthy samples were defined as “core gene suites''. Each gene suite was tested for GO enrichment using Cluster Profiler ^81^. Specifically, the ontology enrichment was performed using the enrichGO function (parameters OrgDb = "org.Hs.eg.db", ont = "BP", readable = TRUE, pvalueCutoff = 0.01, pAdjustMethod = "BH" against a gene universe of all genes that were expressed in our dataset), results were filtered using gsfilter (max = 200) and simplified using the simplify function (cutoff = 0.5). Where multiple GO terms matched identical sets of genes of interest, they were filtered to keep the GO term with the lowest adjusted p-value. The top six remaining enriched GO terms were categorised (**Fig 2d**) according to their relevant ancestor GO term that sat one level below Biological Process (GO:0008150) in QuickGO (<https://www.ebi.ac.uk/QuickGO/>); where two ancestor terms were possible, the most specific term was selected manually. Scaled expression of the 39 core cancer genes was visualised in a heatmap using Complex Heatmap ^82^. Differential cell type abundance was calculated using a modified EdgeR pipeline ^83^, with cell type counts for each patient used as input and with design matrix “~patient + disease”.

**2. Visium spatial transcriptomics and data pre-processing**

***Sample* preparation**

Visium spatial transcriptomics data was generated for twelve samples, five cSCC (patients B18, E15, F21, P30 and P13), three BCC (B18, E15 and F21, i.e. cSCC lesions were also taken from all three patients), and four melanoma samples (6747-085P, 21031-08TB, 48974-2B and 66487-1A). FFPE samples (cSCC P30 and P13, BCC B18, and Melanoma 66487, 48974, 21031, and 6747) and fresh-frozen samples (cSCC E15, F21 and B18, and BCC E15 and F21) were used for data generation.

For fresh-frozen samples, tissue optimisation was performed as per the Visium Spatial Tissue Optimization User Guide (CG000238 Rev A, 10x Genomics), and a tissue permeabilisation time of 18-25 min was selected. For data generation, tissue was cryosectioned at 10 µm thickness and transferred to chilled Visium Spatial Gene Expression Slides (2000233, 10x Genomics, USA), and allowed to adhere by warming the back of the slide. Tissue sections were dried for 1 min at 37°C, fixed in chilled 100% methanol for 30 min, and stained with hematoxylin for 5 min and then eosin for 2 minutes, as per the Visium Spatial Gene Expression User Guide (CG000239 Rev A, 10x Genomics). Brightfield histology images were captured using a 10x objective on an Axio Z1 slide scanner (Zeiss). Brightfield images were exported as high-resolution tiff files using Zen software. This H&E staining and imaging protocol was used to stain all skin sections for histopathological annotation in this study. Libraries were sequenced on a NextSeq500 (Illumina) machine using a High Output 150 cycle kit (Illumina) at the University of Queensland Sequencing Facility, with the read structure as: Read1 - 28bp, Index1 - 10bp, Index2 - 10bp, Read2 - 120bp. The sequencing data was converted to FASTQ format as described above for scRNASeq data and the data was processed as described below. For FFPE samples, the FFPE blocks were sectioned at 5μm by rotary microtome and the sections were processed for spatial sequencing library preparation following the Visium Spatial Gene Expression for FFPE User Guide (CG000407, CG000408, CG000409 - 10x Genomics). The imaging and sequencing were similar to the protocol for the fresh-frozen samples as described above.

**Data *processing***

Reads were trimmed by cutadapt/1.8.3 to remove sequences from poly-A tails and template-switching-oligos. SpaceRanger V1.0 was used to map FASTQ reads to the human reference genome (version GRCh38-3.0.0). After read mapping, spots were removed if they expressed mitochondrial genes more than 20 percent. Thirteen mitochondrial and 103 ribosomal genes were removed from the samples that were originally fresh-frozen; these genes were not captured using the probe-based FFPE library preparation protocol. In total, we captured 14,990 transcriptional capture spots and 36,485 unique genes across all samples. Replicate samples from each of the three cancer types were integrated using Seurat CCI ^1^ with 3000 variable genes, and further processed using SCTransform ^1^. Uniform Manifold Approximation and Projection (UMAP) plots were generated based on the first 20 PCs of cSCC and BCC, and the first 30 PCs for melanoma. Clustering was performed on each integrated cancer dataset separately, using resolutions of 0.5 for cSCC, 0.3 for BCC and melanoma.

***Cell type annotation using spot deconvolution***

We performed Visium spot deconvolution using the Robust Cell Type Decomposition (RCTD) method ^4^, using raw gene counts and with the Level 1 cell type annotations from our single-cell cSCC atlas used as a reference. RCTD was run in doublet mode “full”, and RCTD scores were weighted to a maximum value of 1, to indicate the proportion of each spot predicted to be taken up by a given cell type. Neighbourhood cell type composition was determined by calculating the average RCTD score for each cell type in each cluster.

***CCI analysis***

Visium CCI analysis was performed for each Visium dataset separately using stLearn ^5^. For each of the twelve samples in turn, raw Visium data was input into stLearn, and spots expressing ≥20% mitochondrial reads, and genes expressed in <3 spots were filtered. Data were normalised using stLearn’s normalize_total function. CCI analysis was performed at the spot level for each sample separately, using stLearn’s st.tl.cci.run function with parameters min_spots = 20 and n_pairs = 10,000.

**3. GeoMx data generation and pre-processing**

***Sample processing***

Formalin-fixed skin samples from three cSCC patients (B18, R01 and P04) were profiled by NanoString GeoMx DSP RNA and protein assays. For each assay, 24 regions of interest (ROIs) of variable size (approximately 300-1,500 cells/ROI) were selected based on tissue morphology from adjacent H&E images, representing 12 pairs of adjacent cancer and immune tissue regions (two pairs from patient P04, six from B18 and four from R01) **(Fig S4)**.

Sample preparation for both assays followed standard procedure. Briefly, freshly-cut FFPE sections of 5 µm thickness were placed onto a glass slide. After baking for 1 h at 60°C, slides were processed on a Leica automation platform, with three major processing steps: 1) slide baking and dewax, 2) antigen retrieval for 20 min at 100°C, and 3) 1ug/ml Proteinase K treatment for 15 min. Slides were then removed from the instrument and were incubated with the relevant probe cocktail (i.e. the GeoMx Cancer Transcriptome Atlas assay (CTA) assay probe cocktail for the RNA assay, or the GeoMx protein immuno-oncology (IO) cocktail incubation for the protein assay) overnight. The following day, slides were washed and applied with morphology marker incubation before loading onto the GeoMx machine for processing.

For the RNA assay, cell segmentation markers used were CD3E (red), PTPRC (yellow), KTR5 (green) and DAPI (blue). For gene expression quantification, we used the CTA probe panel, of which the majority of genes are related to cancer biology (i.e. global immune response, microenvironment immune activities, tumour reactivity to treatment, tumour inflammation signature and cancer metastasis), but also includes 31 endogenous control housekeeping genes and one negative control spike-in. We captured RNA quantification information for 1,825 genes using this panel.

For the protein assay, equivalent cell segmentation morphology markers were used as in the RNA assay, namely CD3 (red), CD45 (yellow), PanCk (green), DAPI (blue). The IO protein panel includes a total of 48 protein markers, including three positive controls (Histone H3, GAPDH, and S6) and three negative controls (Rb IgG, Ms IgG1, and Ms IgG2a).

***Differential expression analysis (protein and RNA data)***

Differential gene expression analysis was performed for RNA and protein data separately using the edgeR pipeline ^7^. Analysis was performed using log Counts Per Million (logCPM)-transformed expression values. Prior to analysis of RNA data, five lowly-expressed genes were filtered using the function *filterByExpr*, leaving 1,820 markers for downstream analysis. No proteins were filtered prior to analysis, leaving 48 protein markers for downstream analysis. Library size normalisation was performed using the calcNormFactors function. Differential expression analysis between CD45+ and PanCK+ regions was performed using the cpm.DGEList function with the design matrix “~patient + group” to compare between-group differences while accounting for between-patient differences. We used the *glmQLFit* quasi-likelihood negative binomial model to fit a linear model to the expression data. Genes/proteins were categorised as differentially expressed if they received a false detection rate (FDR) ≤ 0.05 and log_2_ fold change >1.0. GO enrichment analysis of differentially expressed genes from the RNA assay was performed as described above for scRNASeq comparisons. GO analysis was not performed for protein data due to the small number of captured markers available to serve as the background universe for statistical testing.

***Spatial Deconvolution (RNA data)***

We used the R package SpatialDecon^8^ to assess the abundance of different cell types across 12 CD45+ ROIs. SpatialDecon estimates cell type abundance within spatially-resolved gene expression datasets. SpatialDecon takes as input the normalised gene expression data (here, the logCPM-normalised GeoMx data). It also uses the function derive_GeoMx_background to calculate expected background counts for the data using the normalised gene expression matrix. Cell types were assigned based on an inbuilt reference expression profile of expected cell types, the SafeTME matrix included with the SpatialDecon package. This reference contains the expression profiles of 906 genes over 18 cell types, and was purposely designed to exclude genes commonly expressed by cancer cells ^8^. For this reason, we did not perform deconvolution for PanCK+ segments as the reference profiles available for GeoMx data are biassed towards immune cells, and specifically exclude the cancer-associated genes expected to be enriched in the PanCK+ fraction.

**4. CosMx data collection and pre-processing**

***Sample preparation***

Nanostring CosMx data was generated for nine samples, three cSCC (patients B18, P30 and P13), two BCC (patients B18 and D12), and four melanoma samples (patients 6747-085P, 21031-08TB, 48974-2B, 66487-1A). FFPE tissue sections were prepared as previously described ^9^. Briefly, 5 µm tissue sections on VWR Superfrost Plus Micro slides (Cat. No. 48311-703) were baked overnight at 60°C to improve tissue adherence to the slides, then prepared for *in situ* hybridisation (ISH) by deparaffinisation and heat-induced epitope retrieval (HIER) at 100°C for 15 min. For all samples except cSCC Patient P30, HIER was performed in a pressure cooker using ER2 epitope retrieval buffer (Leica Biosystems product, EDTA-based, pH 9.0). HIER for Patient P30 was performed on a Leica Biosystems Bond RX automated tissue handler using ER1 epitope retrieval buffer (Leica Biosystems product, citrate-based, pH 6.0).

Following HIER, tissue sections were digested with Proteinase K diluted in ACD Protease Plus at 40°C for 30 min (3 µg/ml Proteinase K for all samples except Patient P30, which was at 5 µg/ml). The tissue sections were washed twice with diethyl pyrocarbonate (DEPC)-treated water (DEPC H_2_O) and incubated in 1:3,000 diluted† fiducials (1:5,000 for Patient P30) (Bangs Laboratory) in 2X SSCT (2X saline sodium citrate, 0.001% Tween-20) solution for 5 min at room temperature in the dark. Excess fiducials were removed by rinsing the slides in 1X PBS, before tissue sections were fixed with 10% neutral buffered formalin (NBF) for 5 min at room temperature. Fixed samples were rinsed serially with Tris-glycine buffer (0.1M glycine, 0.1M Tris-base in DEPC H_2_O) and 1X PBS for 5 min each before blocking with 100 mM *N*-succinimidyl (acetylthio) acetate (NHS-acetate, ThermoFisher) in NHS-acetate buffer (0.1M NaP, 0.1% Tween PH 8 in DEPC H_2_O) for 15 min at room temperature. The sections were then rinsed with 2X saline sodium citrate (SSC) for 5 min and an Adhesive SecureSeal Hybridization Chamber (Grace Bio-Labs) was placed over the tissue.

NanoString ISH probes were prepared by incubation at 95°C for 2 min and immediately placed on ice, and the ISH probe mix (1 nM ISH probe, 1X Buffer R, 0.1 U/μL SUPERase•In [Thermofisher] in DEPC H_2_O) was pipetted into the hybridisation chamber. The chamber was sealed to prevent evaporation, and hybridisation was performed at 37°C overnight. The tissue sections were then washed twice in 50% formamide (VWR) in 2X SSC at 37°C for 25 min, washed twice with 2X SSC for 2 min at room temperature, and blocked with 100 mM NHS-acetate in the dark for 15 min. In preparation for loading onto the CosMx SMI instrument, a custom-made flow cell was attached to the slide.

***CosMx SMI instrument run***

RNA target readout on the CosMx SMI instrument was performed following the published protocol ^9^. Briefly, the assembled flow cell was loaded onto the instrument and washed with Reporter Wash Buffer to remove residual air bubbles. A preview scan of the entire flow cell was taken, and 15-25 FOVs were selected to match areas of interest identified by H&E staining of adjacent serial sections. For some tissues we selected closely-adjacent FOVs to capture almost the entire tissue biopsy, while for other tissues we selected FOVs covering a smaller fraction of the biopsy. RNA readout began by flowing 100 μl of Reporter Pool 1 into the flow cell and incubation for 15 min. Reporter Wash Buffer (1 mL) was flowed into the flow cell to wash away unbound reporter probes, and Imaging Buffer was added to the flow cell for imaging. Nine Z-stack images (0.8 μm step size) were acquired for each FOV, and photocleavable linkers on the fluorophores of the reporter probes were released by UV illumination and washed away with Strip Wash buffer. The fluidic and imaging procedure was repeated for the 16 reporter pools, and the 16 rounds of reporter hybridisation-imaging were repeated multiple times to increase RNA detection sensitivity.

After RNA readout, tissue samples were incubated with a 4-fluorophore-conjugated antibody cocktail against CD298, S100b/PMEL17 (PanCK for Patient P30), CD45, and CD3 proteins and DAPI stain in the CosMx SMI instrument for 1 h. After unbound antibodies and the DAPI stain were rinsed with Reporter Wash Buffer and Imaging Buffer was added to the flow cell, nine Z-stack images for five channels (four antibodies and DAPI) were captured.

***Primary data processing***

CosMx data was processed as described previously ^9^. Briefly, the image processing comprises three main steps: registration, features detection, and localisation. 3D rigid image registration was performed with the use of fiducial markers embedded within the tissue sample. Each subsequent image stack was registered using phase correlation and individual channels were aligned through a pre-calibrated affine transformation.

Diffraction-limited features were identified that represented the fluorescence response from a single molecule. A 2D Laplacian of Gaussian (LoG) filter was applied to remove background and enhance the encoded reporter signatures. Potential reporter locations were identified as local maxima using a 3D nearest neighbour search. Sub-pixel localisation for each feature was obtained by fitting a 2D polynomial to the maxima in the X, Y, and Z axes.

***Secondary analysis and decoding***

The XYZ locations of all individual reporter-binding events were used to determine the presence of individual transcripts. Briefly, each unique location with at least one reporter-binding event is considered a ‘seed’, and all neighbouring locations to each seed with at least one reporter-binding event were determined. All possible four reporter combinations of unique reporter probes in a seed’s neighbourhood, such that at least one of the four reporter-binding events was present at the seed location, were then matched with gene-specific barcodes to detect the presence of a gene in a seed’s neighbourhood.

***Cell segmentation algorithm***

NanoString’s cell segmentation pipeline is a combination of image preprocessing and machine learning techniques ^9^. Briefly, the pipeline takes tissue images stained with both nuclear and membrane markers to perform rescaling, normalisation, image deconvolution, and boundary enhancement. Image subtraction was performed between the nuclear and the membrane channels to enhance the contrast between adjacent cells while reducing auto-fluorescence signal. The preprocessed images were fed into pre-trained Cellpose neural network models ^10^ for both nuclear and cytoplasm modes of segmentation. Results from two segmentation tasks were combined to select the best results from each mode by analysing intersection and union between all segmented cells.

***Single-cell quality control and filtering***

Single-cell expression profiles were derived by counting the transcripts of each gene that fell within the area assigned to a cell by the segmentation algorithm. Cells with fewer than 20 total transcripts or that have abnormally large cell area, which likely represent doublets, were omitted from the analysis. A normalised expression profile was defined for each cell by dividing its raw counts vector by its total counts. A separate UMAP projection was computed for each cancer type.

***Cell type annotation***

We used our scRNASeq data to construct a reference profile matrix of cell types for CosMx annotation. This profile listed the average expression of each of the 960 genes in the CosMx panel for each of the cell types in our scRNASeq Level 1 cSCC annotations. This reference matrix was then used to initialise an Expectation-Maximisation (EM) algorithm that alternated between assigning cell types and updating the cluster profiles. The EM algorithm was built around the following likelihood model:

$$Y_{ij} \sim NegBinom(mu = b_{j}+s_{j,k(j)}x_{ik(j)}, size = 10)$$

Where ­$i$ indexes genes, $j$ indexes cells, $k$ indexes cell types, $Y_{ij}$ is the raw counts of gene $i$ in cell $j$, $b_{j}$ is the expected per-gene background counts in cell $j$, $X_{ik(j)}$ is the expected expression level of gene $i$ in cell type $k(j)$, and $s_{j}$ is a scaling factor corresponding to the total counts in cell $j$. b, X and s were pre-defined where b is calculated from a direct measurement of the background observation rate using included CosMx control target observations, X represents the cell type expression profile measurements derived from scRNASeq data, and $s_{j}$ is calculated as a cell's total counts divided by the total counts in the profile under consideration.

In the M-step, cluster assignments $k(j)$ are updated by assigning each cell $j$ to the cluster $k$ under which it achieves highest likelihood. In the E-step, each column $k$ of the matrix $X$ is estimated by taking the maximum likelihood estimate calculated by applying the above likelihood model to all cells assigned to cell type $k$. To preserve the influence of the original reference profiles, the cell type expression profiles estimated in the E-step are averaged with the original reference profiles before proceeding to the M-step. The E-step and M-step are iterated until convergence.

**5. Cell type co-localisation and CCI with Polaris spatial proteomics**

***Sample processing***

Polaris data was generated for three cSCC samples (patients B18, P04 and R01). Multispectral analysis of FFPE tissue utilised the MOTiF PD-1/PD-L1 kit (Akoya Biosciences, Cat. No. OP-000001). Staining with Leica BOND RX (Leica Biosystems) and imaging with Vectra Polaris (Akoya Biosciences) was performed at The Walter and Eliza Hall Institute (WEHI) Histology core facility as per kit manufacturer instructions. Briefly, tissue was stained through cycles of incubation with primary antibody, anti-IgG polymer HRP and covalent labelling with Opal TSA fluorophores, followed by HIER to remove bound antibodies prior to subsequent antibody cycles. Target markers included CD8 (Opal 480), PD-L1 (Opal 520), PD-1 (Opal 620), FoxP3 (Opal 570), CD68 (Opal 780, PanCK (Opal 690) and spectral DAPI DNA stain. Whole slide multispectral scanning was performed on the Vectra Polaris instrument using automatically adjusted exposure settings. Image tiles were spectrally unmixed in InForm (Akoya Biosciences), then restitched in QuPath software^11^.

***Data processing***

We applied StarDist cell detection ^12^ to segment cell boundaries and measure protein marker signals within these boundaries, transforming the imaging data into non-imaging, single-cell protein expression counts (i.e. the mean signal intensity for each protein). To remove artificially high background from fluorescent intensities (outliers), the maximum value of each marker was capped at the 95th percentile. Leiden cell type clustering was implemented in Scanpy ^13^ using a standard single-cell analysis pipeline. Resulting clusters were assessed to determine cell type identity. Cells showing very low expression of all six protein markers were classified as “unidentified” and removed from downstream analysis.

Using the cell clustering information, we sought to identify the subregions where cancer and immune cells co-localised and possibly interacted. Spatial information at the subcellular level can facilitate a lower level of spatial analysis, which uses the cell geometry and density to assist the interaction analysis. To unbiasedly detect zones of cancer and immune cell co-localisation, and potential interaction, we applied a scanning window strategy called STRISH ^14^ to identify regions containing double-positive CD8+ PD-1+ immune cells in the immediate vicinity of double-positive PanCK+ PD-L1+ cancerous epithelial cells. STRISH divides the image into tiles and applies a scanning window strategy to identify zones of co-localisation. Specifically, STRISH starts by dividing the image into four non-overlapping tiles, each one-fourth of the size of the original slide scan, and gradually splits these large tiles into smaller windows, until a given cell count threshold is met (here, <100 cells per window). Results are then scored to measure the density of cells’ expressed markers (e.g. specific ligands and receptors), normalised across the windows with a positive score, and used to plot a heatmap showing the co-localisation scores of the two markers or cell types being tested.

We also calculated co-occurrence scores between cell types using a conditional probability strategy. The samples were scanned and stitched into large whole-slide images (>2000 µm in length) while preserving the original form of the biopsy. Cell type co-occurrence was calculated between pairs of cell types across increasing distance intervals (0 to 600 µm). Note that because the skin biopsies are wide but narrow, the cell type co-occurrence scores are expected to diverge after converging. The function to calculate the conditional probability of cell type co-occurrence at different distance intervals was adapted from the co-occurrence analysis method implemented in the Squidpy package ^15^.

**6. RNAscope for multiplexed RNA in-situ hybridisation and data pre-processing**

***Sample processing***

RNA *in situ* hybridisation was performed using RNAscope for three cSCC and three BCC samples from the same patients (patients B18, E15 and F21). Two consecutive sections of 10 µm thickness were taken from OCT-embedded tissue blocks for the assay and for a negative control, respectively. Slides were stained with a mixture of five RNA probes, namely THY1 (ADV430611T2), IL34 (ADV313011T3), CSF1R (ADV310811T4), CD207 (ADV809521T7), and ITGAM (ADV555091T8) designed by ACD (Cat. No. 324110). The negative control slide was stained with the RNAscope HiPlex 12 Negative Control Probe included in the kit. After RNA hybridisation, specific signal was amplified with high efficiency using RNAscope HiPlex Amp reagents. Tissue images were captured by an Axio Z1 slide scanner (Zeiss) after adjusting fluorescent intensity. The imaging was performed in two rounds with identical microscopy parameters for each round. The first round captured nuclei (DAPI), THY1 (Cy3), IL34 (Cy5) and CSF1R (Cy7), while the second captured nuclei, CD208 (Cy5) and ITGAM (Cy7). Images from the two rounds were processed, stitched, and post-processed by ZEN software.

***Data processing***

We sought to map interactions between the LR pair IL34_CSF1R using RNAScope data. First, similar to the Polaris data processing methods described above, we performed cell segmentation across the images using starDist ^12^. Next, we converted imaging data to a quantitative RNA expression matrix by determining the mean intensity of RNA fluorescent signal within each cell boundary. We performed data pre-processing to remove outlier signals and false cell detection events. Subsequently, we applied STRISH to detect cell co-localisation, by splitting each scanned image into multiple non-overlapping windows and quantified the IL34 and CSF1R co-localisation within every window. Each window expressing both CSF1R and IL34 were statistically tested against the random combination of two pairs of markers available for the same window and across all IL34_CSF1R-positive windows. Permutation tests were performed testing different random LR pairs besides the target CSF1R and IL34 to determine the most significant ones.

***7. CODEX (Phenocycler-Fusion, Akoya Biosciences)***

Single-cell spatial phenotyping of the Melanoma FFPE slides was performed in collaboration with Akoya Biosciences (The Spatial Biology Company, Marlborough, MA, USA) on the Phenocycler^®^-Fusion (PCF) platform. The tissue slides were stained with a 33-plex antibody panel, including markers for immune checkpoints, immune cell lineages, activation states, tissue structure, and metabolism, in a single step. Then, on the Phenocycler^®^ -Fusion, combinations of three antibodies were visualized by utilising iterative fluorescent-reporter addition and imaging cycles. Commercially available oligo-conjugated antibodies were obtained from Akoya Biosciences (The Spatial Biology Company, Marlborough, MA, USA). The details of the antibodies are shown in the **Supplementary Table S2**.

PCF (Massachusetts, US) immunostaining was performed according to the PCF user manual and previously published method (Goltsev et al. 2018, Black et al. 2021). First, 5 µm tissue sections were baked for 12 hours at 55℃ on a slide warmer (Premiere XH-2004) and dewaxed through two rounds of 5-minute incubation in Histochoice Clearing Agent (VWR, PN# H103-4L). Subsequently, the FFPE sections were dehydrated by sequentially incubating in 100% ethanol (Sigma Aldrich, PN# 79317-16GA-PB), 90% ethanol, 70% ethanol, 50% ethanol and 30% ethanol each twice for 5 minutes. Slides were then rinsed three times in distilled water for 5 minutes to ensure no carryover of ethanol before target retrieval. Next, heat-mediated epitope retrieval was performed by incubating slides in a Coplin jar containing Tris EDTA solution (pH 9; Dako, #S2367) in a pressure cooker for 20 minutes. After 20 minutes, the Coplin jar was removed from the pressure cooker and left to equilibrate at room temperature for at least 30 minutes. Slides were then rinsed through two rounds of distilled water for 2 minutes and stored in Hydration Buffer from the PhenoCycler Staining Kit (Akoya Biosciences, MA, #7000008) until ready for staining.

After the pretreatment, the FFPE sections were stained with Autostainer (Parhelia). Samples were first allowed to equilibrate to room temperature in Staining Buffer from the PhenoCycler Staining Kit for 20-30 minutes, followed by incubation in a pre-blocking solution made of N, J, and S blockers in the Staining Buffer from the PhenoCycler Staining Kit (Akoya Biosciences, MA, #7000008). Then the FFPE sections were stained with an antibody cocktail with optimal dilutions of each antibody and N, J, S, and G blockers for 3 hours at room temperature. After washing with Staining buffer and post-fixation with 1.6 % PFA for 10 minutes, samples were fixed in methanol (Sigma Aldrich, MO, #34860) for 5 minutes. Lastly, PhenoCycler Fixative reagent was applied for fixation for 25 minutes, and slides were stored in Storage Buffer from the PhenoCycler Staining Kit at 4℃ until ready to image.

Before imaging, the antibody-stained slides were equilibrated at room temperature in 1X CODEX Buffer for 10 minutes. Next, the flow cell (Akoya Biosciences, #240204) was assembled onto the slide as described in the user manual. Then, the slides were incubated in 1X CODEX buffer again for 10 minutes to ensure secure sealing of the flow cell to the slide.

The reporter, fluorescently labeled–oligonucleotide probes (fluorescent-reporters) that corresponded to DNA-barcoded antibodies for multiple cycle immunostaining, was prepared as described in the PhenoCycler Fusion User Guide. Reporter stock solution was prepared with 10X CODEX Buffer (#7000001), Assay Reagent (#7000002) and Nuclear Stain (#7000003) from Akoya Biosciences. Individual tubes of 3 reporters/cycle were diluted in reporter stock solution. Subsequently, a 96 well plate was prepared with 1 well/cycle containing the corresponding working reporter solution for that cycle. Blank cycles containing only reporter stock solution without any added reporters were included as the first and last cycle in each run for subtraction of autofluorescence background. The PCF is an integrated ultrahigh plex cycling platform, including the PhenoCycler automated fluidics cycler and the PhenoImager Fusion imaging system which automates the entire process of reporter hybridization, imaging, and dehybridization to capture whole slide images of three markers (+DAPI for nuclear staining) at a time. First, the imaging protocol was set up using Fusion Experiment Designer software according to the manufacturer’s protocol. Then, two blank cycles (only DAPI nuclear stain without any fluorescent reporters) at the first and the last cycle of the experiment were run to evaluate the level of autofluorescence and subtract background using Fusion software. After images of all cycles had been acquired, the final QPTIFF file containing a composite image of all markers was viewed using the QuPath software (<https://qupath.github.io/>), where each channel can be turned on and off individually or collectively to reveal the spatial expression pattern of the marker(s) of interest. The imaging protocol was established with the Fusion Experiment Designer software V1.0 according to the manufacturer's instructions. Following the acquisition of all cycles, a final composite QPTIFF image file was exported from Phenocycler Fusion software V1.0.

**8. Spatial glycomics**

The tissue sections mounted on ITO were placed onto a heating plate at 65 °C for 60 min. Paraffin was removed by incubating the slides in xylene (2 x 2 min) and tissue sections were re-hydrated by different washes: 100% ethanol (1 x 2 min), 90% ethanol (1 x 2 min), 70% ethanol (1 x 2 min), 50% ethanol (1 x 2 min), water (1 x 2 min) and completely dried in the desiccator. The sialic acids were stabilised by methylamidation (PMID: 38953530). The derivatization was performed by incubating the tissue slides in derivatization solution: (1 M methylamide hydrochloride, 0.5 M methylmorphine and 50 mM PyAOP in DMSO) for 2 x 5 min at ambient temperature. After derivatization, tissue sections were washed in 100% ethanol (2 x 2 min), Carnoy solution (60% ethanol, 30% chloroform, 10% acetic acid, 1 x 5 min) and water (1 x 1 min). Slides were dried in a vacuum desiccator (~10 min) before antigen retrieval. Citraconic anhydride buffer (~10 mL, pH = 3) based antigen retrieval was carried out in a commercial vegetable steamer (Philips All-In-One Cooker, HD2237) for 20 min. Following the antigen retrieval, slides were cooled down for 5 min, rinsed in water and dried in the vacuum desiccator before enzymatic deglycosylation. PNGaseF (0.1 µg/µL in 25 mM Ammonium bicarbonate) was deposited using the HTX TM-Sprayer (HTX Technologies, USA), 15-layer passes at 25 µL/min, velocity of 1200 mm/min, crisscross pattern, 3.0 mm track spacing, nitrogen gas pressure was set to 10 psi. N-glycans were released in a 3-hour incubation at 37 °C in a humidity chamber protected from evaporation. After incubation the ITO slides were briefly dried in the vacuum desiccator and the matrix (10 mg/mL CHCA, 50% ACN and 0.1% TFA) was applied using HTX TM-Sprayer (5 layers, 0.1 mL/min, 80 °C, velocity of 1300 mm/min, crisscross pattern, 2.5 mm track spacing, nitrogen gas pressure was set to 10 psi).

MALDI MSI data were acquired using a rapifleX MALDI TOF/TOF mass spectrometer (Bruker, Daltonics) equipped with a Smartbeam 3D 10 kHz laser and operated in a positive-ion reflectron. The laser power was optimized at the start of each run and then held constant during the MALDI–MSI experiment. At each sampling position, 250 shots were used to acquire data at 20 μm pixel resolution for the *m*/*z* 920–3500 range. All of the methods used were internally calibrated using commonly present neutral glycans across the mass range. Spectra were normalized by total ion count (TIC) normalization of all data points, unless otherwise stated. Preprocessing, background reduction and visualisation of data was performed using SCiLS Lab (Bruker, Daltonics, version 2025b core).

### **Supplementary Notes - Additional Results**

P. Prakrithi^1,2#^, Laura F. Grice^1,2,3#^, Feng Zhang^1,2,4#^, Levi Hockey^1,2^, Samuel X. Tan^5^, Xiao Tan^1,2^, Zherui Xiong^1,2^, Onkar Mulay^1,2^, Andrew Causer^1,2^, Andrew Newman^1,2^, Duy Pham^1^, Guiyan Ni^1^, Kelvin Tuong^6^, Xinnan Jin^1,2^, Eunju Kim^2,^, Minh Tran^1^, Hani Vu^1,2^, Nicholas M. Muller^5^, Emily E. Killingbeck^7^, Mark T. Gregory^7^, Siok Min Teoh^1^,Tuan Vo^1^, Min Zhang^8^, Maria Teresa Landi^9^, Kevin M. Brown^9^, Mark M. Iles^10^, Zachary Reitz^7^, Katharina Devitt^5^, Liuliu Pan^7^, Arutha Kulasinghe^5^, Yung-Ching Kao^5^, Michael Leon^7^, Sarah R. Murphy^7^, Hiromi Sato^7^, Jazmina Gonzalez Cruz^5^, Snehlata Kumari^5^, Hung N. Luu^11^, Sarah E. Warren^7^, Chris McMillan^12,13^, Joakim Henricson^14,15^, Chris Anderson^14,15^, David Muller^12,13^, Arun Everest-Dass^16^, Blake O’Brien^17^, Huanwei Wang^18^, Mathias Seviiri^18^, Matthew H. Law^18, 19, 20^, H. Peter Soyer^5^, Ian Frazer^5^, Youngmi Kim^7,21^, Mitchell S. Stark^5^, Kiarash Khosrotehrani^5^, Quan Nguyen^1,2^

**Note S1: Confimation of cell types, differentially expressed genes, and cancer markers using GeoMx protein and RNA assays**

We generated GeoMx data from three cSCC patients (R01, P04 and B18) using both protein and RNA modalities, and captured data from CD45+ and PanCK+ tissue segments across a total of 12 paired regions of interest (ROIs) (**Fig S8**).

**Differential expression analysis (protein data)**

Differential protein expression analysis between CD45+ and PanCK+ segments revealed that 24 of 48 captured protein markers showed differential abundance between segments, with 10 genes enriched in CD45+ segments and 14 in PanCK+ segments. As expected, PanCK was amongst the differentially expressed proteins enriched in the PanCK+ segment, and CD45 was enriched in the CD45+ segment. Numerous other cancer-associated markers were also upregulated in the PanCK+ segments, including oestrogen receptor alpha (ER.alpha), which has been associated with cutaneous cSCC progression ^16^ and is negatively associated with survival in oral cSCC ^17^, the cell proliferation-associated protein Ki.67, and the melanocyte and melanoma marker MART-1 ^18^ (whose encoding gene, *MLANA*, is mapped in melanoma tissue in **Fig S11a**). CD44 and CD80 were also upregulated in the PanCK+ segment; these markers have both been associated with tumour-initiating stem cells ^19^. Meanwhile, immune markers including monocyte/macrophage markers CD163 and CD14 and T cell markers CD3 and CD4 were all enriched in the CD45+ fraction.

**Differential gene expression analysis (RNA data)**

We first visualised the RNA expression of key marker genes associated with rarer cell types from our Level 2 scRNA-seq cSCC atlas (**Fig S4e, Fig S9b**). We observed a statistical upregulation in the CD45+ segments of M2 marker gene *CD163* and fibronectin *(****Fig S9b****)* (plus fibroblast marker FAP, which was differentially expressed in the RNA **(Fig S9b**) and protein (***Fig S10n***) data). Markers for the other tested cell types (except Tregs) followed similar qualitative trends to the protein data, appearing to be more highly expressed in the CD45+ segments, albeit without statistical significance. The presence of these markers in both the protein and RNA datasets lends further support for the existence of these cell types.

We also performed differential gene expression analysis of CD45+ and PanCK+ ROIs for the RNA data, and uncovered a total of 267 of 1,825 genes exhibiting differential expression between conditions (logFC > 1.5 and FDR ≤ 0.05), including 107 genes enriched in CD45+ segments and 160 genes enriched in PanCK+ segments (**Fig S9c**). Amongst the top 50 PanCK-associated genes were four KC marker genes (*KRT5, KRT6A.B.C, KRT6B, and KRT14*) ^20^, and two members of the S100 gene family (*S100A8* and *S100A9*), which have been linked with cSCC tumorigenesis ^21^. The rest of the top 50 PanCK+ markers also include cSCC markers *NECTIN1, EGFR* and *COL17A1* (**Fig S9c**)*.* Thirty-three of the genes that were differentially expressed between CD45+ and PanCK+ segments (**Fig S9a**) were also members of the core healthy or cSCC gene suites identified in our scRNASeq atlas (**Fig S4f**), including six cancer-associated genes and 27 healthy-associated genes. *CD74* was upregulated in the CD45+ segment in the GeoMx data and was detected in the core cSCC gene suite. All other genes were upregulated in the PanCK+ segment (***Fig S9a***). It should be noted that both the CD45+ and PanCK+ segments in the GeoMx dataset are derived from cSCC samples, while the scRNASeq core gene suites were calculated separately for healthy and cancer biopsies, but without regard to cell type.

GO enrichment analysis of the top 100 differentially expressed genes per segment (**Fig S9e**) revealed that the PanCK fraction is enriched for GO terms associated with keratinisation (GO:0031424) and cell junction organisation (GO:0034330). Meanwhile, the CD45+ fraction is strongly associated with angiogenesis (GO:0001525), wound healing (GO:0042060), ECM-related genes (GO:0030198 and GO:0043062), including collagen genes *COL1A1, COL1A2, COL5A1* and *COL3A1*. These ROIs were also enriched for GO terms related to cell chemotaxis (GO:0060326) and immunological functions such as the humoral immune response (GO:0002920), consistent with these regions being selected on the basis of CD45 expression.

**Cell-type** **Deconvolution analysis (RNA data)**

We performed deconvolution using SpatialDecon ^8^ to estimate the cell type composition of our twelve CD45+ ROIs (**Fig S9f**). This analysis revealed that CD45+ ROIs have multiple types of both B cells (naïve and memory B cells) and CD4 T cells naïve and memory T cells), plus T regulatory cells, macrophages, and endothelial cells consistently present across all ROIs in abundance. Some samples also show presence of CD8 T cell naïve (CD8 naive and memory T cells) and NK cells. As the SpatialDecon reference profile is designed primarily for detecting immune cells, and explicitly excludes known cancer markers ^8^, we did not perform a comparable deconvolution analysis for the PanCK+ tumour segment regions (*see Methods for further details*).

**Note S2: Polaris for a targeted study of interactions via PD-1 and PD-L1**

We used Polaris high-plex protein staining to perform a focussed analysis of cell co-localisation and CCI in spatial context for three cSCC patients (R01, P04 and B18). Based on combinations of an antibody panel of six proteins (**Fig S23a**), we were able to identify major cell types, including epithelial (PanCK+), inmate immune (CD68+), and adaptive immune (CD8+, FoxP3+) cells (**Fig S23b**) from all tissue samples.

In a non-cancer state, the immune cell-bound PD-1 receptor acts as an immune checkpoint inhibitor which, when bound by its ligand PD-L1 (or PD-L2), dampens the immune response ^31,32^; this serves as an important feedback mechanism limiting autoimmunity ^33^. In cancer, PD-L1 expression is co-opted by cancer cells as an immune evasion mechanism, inducing immune suppression of activated T cells ^34^. For this reason, inhibitors of the PD-L1/PD-1 interaction have proved valuable for the treatment of metastatic skin and other cancers ^32^. We therefore sought to identify immune cells expressing PD-1 and cancer cells expressing PD-L1 sitting in close proximity to one another. While this does not provide direct evidence of CCIs, it does suggest their cross-activity.

We employed our pipeline, STRISH ^14^, to automatically and unbiasedly detect regions containing T-cells double positive for CD8 and PD-1 in the immediate vicinity of cancerous epithelial cells, which themselves are double positive for PanCK and PD-L1 (**Fig S23c**). Interestingly, for patient B18, we found that the PD-1+ immune cells and PD-L1+ epithelial cells mostly co-localised at the interface of cancer and immune infiltration, as defined by a pathologist’s annotation (**Fig S23a**). Similar patterns were observed by applying STRISH pipeline to the tissue samples from patients R01 and P04 (data not shown).

We then investigated cell type co-localisation more broadly using spatial community analysis ^35^. We performed a clustering analysis to group tissue regions with similar local densities of various cell types, which for patient B18 resulted in the identification of five cell communities distributed across the tissue (**Fig S23d**). Quantitative assessment of cell composition in each community allowed us to deduce biologically interpretable features of these communities (**Fig S23e**). In particular, Community 0 has a very high density of PD-L1+ epithelial cells, while Communities 2 and 3 are enriched for three types of CD8+ T cells (**Fig S23e**). Visual inspection of the location of Communities 2 and 3 reveals their spatial proximity (**Fig S23d**), which may be explained by the tendency of cancerous epithelial cells to reside densely around the cancer nest, and for immune cells to sit under the epidermis layer ^36^. We also observed that Communities 1 and 4 group together (**Fig S23e**), and that these communities contain a mixture of both cancer and immune cell populations. The spatial distribution of the mixed cancer/immune communities (**Fig S23d**) aligns with the spatially-specific interaction observed between PD-1 and PD-L1 along the interface between cancer and immune cells (**Fig S23c**).

We investigated the immune-only (Communities 2 and 3) and cancer-immune (Communities 1 and 4) groups in greater detail by performing a cell type co-occurrence analysis (**Fig S23f-g**). Briefly, Ripley's K-function was used to summarise the co-occurrence of specific cell types with all other cell types across increasing distance intervals ^15^. When PD-1+ T cells were set as the reference cell type, these cells were found to co-occur with all three T cell types (i.e. with themselves, CD8+ T cells, and FoxP3+ CD8+ T cells) at a significantly higher rate than a random distribution, across distances between 0 to 400 µm (**Fig S23f**). PanCK+ epithelial cells were found to significantly co-occur with themselves and with double-positive PD-1+ CD68+ immune. Thus, our image-based analysis pipeline has detected biologically-meaningful neighbourhoods of cell types in cancer tissues.
